# Lipoengineering of Biomolecular Condensates Controls Material Properties and Multiphase Hierarchy to Guide Organoid Morphogenesis

**DOI:** 10.64898/2026.04.08.717265

**Authors:** Zhiwei Huang, Md Mahbubul Alam, Mahtab Shokri, Harshavardhan C. Savitrinarayana, Sisila Valappil, Tanushree Agarwal, Rob M. Scrutton, Laiba Maryam, Asma Gulzar, Jinying Wang, Dominic J. Tigani, Liege A. Pascoalino, Akshay V. Jadhav, Albert L. Adhya, Alaji Bah, Zhao Qin, Zheng Shi, Michael R. Blatchley, Jianhan Chen, Tuomas P.J. Knowles, Davoud Mozhdehi

**Affiliations:** Department of Chemistry, Syracuse University, Syracuse, NY 13244, USA; Department of Chemistry, University of Massachusetts, Amherst, MA 01003, USA; Department of Chemistry, Centre for Misfolding Disease, University of Cambridge, Cambridge, UK; Department of Chemistry and Chemical Biology, Rutgers, The State University of New Jersey, NJ 08854, USA; Department of Biomedical and Chemical Engineering, Syracuse University, Syracuse, NY 13244, USA; Department of Civil and Environmental Engineering, Syracuse University, Syracuse, NY 13244, USA; Department of Biochemistry and Molecular Biology, SUNY Upstate Medical University, Syracuse, NY 13210, USA

## Abstract

Cells use post-translational modifications (PTMs) to reconfigure biomolecular condensates across length scales, space, and time.^1,2^ While charged PTMs are well-known electrostatic switches,^3,4^ how ubiquitous neutral PTMs shape condensate plasticity and hierarchy remains unclear. Here, we establish a set of design principles for using site-specific lipidation, a class of neutral hydrophobic PTMs, to rationally control properties and interactions of engineered biomolecular condensates. Through systematic analysis of over 80 lipidated synthetic intrinsically disordered proteins (IDPs), we uncovered two distinct axes of control. First, the interplay between the lipid and the local three-residue sequence of its attachment site acts as a programmable switch for *cohesion*—the homotypic interactions that define the material state of the condensed phase— directing assemblies toward dynamic liquids, arrested gels, or ordered fibrillar solids. Second, the lipid, together with the global properties of the IDP scaffold, tunes *adhesion*—the heterotypic interactions that govern condensate miscibility and hierarchical organization. We harnessed these principles to rationally engineer complex, multi-phase architectures and create hybrid hydrogels with programmed microstructure and material properties that guide the morphogenesis of functional intestinal organoids. These findings establish a new framework for lipoengineering advanced biomaterials and provide a blueprint for dissecting structure–property relationships across diverse classes of PTMs.

## Main

Cells dynamically regulate the formation and material state of biomolecular condensates, transitioning them between fluid, gel-like, and solid states to control biochemical reactions,^5^ regulate transcription,^6^ and respond to stress.^7^ A central challenge is to understand the rules that govern these state transitions^8–11^ and harness them to spatiotemporally program the hierarchical assembly of abiotic matter and cellular organization.^12–16^ While sequence-level engineering of IDPs has advanced our ability to program condensate phase separation^17–20^ and multiphasic assembly,^21,22^ it often necessitates extensive modification of the polypeptide. A less explored but more precise alternative is the use of post-translational modifications (PTMs). By leveraging localized chemical changes to tune molecular interactions, PTMs offer an elegant strategy for controlling material properties and condensate behavior.^1,23^ However, the design principles of PTM-based condensate engineering remains poorly defined due to a lack of systematic investigation.

While charge-altering PTMs (e.g., phosphorylation) are intensely investigated for their effects on condensate dynamics via long-range electrostatics,^3,4,24^ the roles of many ubiquitous PTMs, including glycosylation and lipidation, in modulating short-range interactions and condensate properties remains largely unexplored. Lipidation is a particularly compelling PTM for condensate engineering. Beyond its canonical role in membrane anchoring,^25^ lipidation alters the amphiphilicity of a protein and can regulate self-assembly^26^ and liquid-liquid phase separation (LLPS).^27,28^ Recent studies of N-myristoylation in EZH2 and FSP1 illustrate that lipidation promotes condensate formation yet drives distinct outcomes, including cancer growth or suppression.^29,30^ However, deconvoluting the biophysical contribution of lipidation in native contexts is challenging due to the confounding effects of concurrent PTMs and complex interactomes. Compounding this, lipidation sites are often enriched within intrinsically disordered regions (IDRs).^31^ Here, lipids may act not just as membrane anchors, but also as structural switches that bias the IDR conformational landscape.^32^ To decouple these variables and define the fundamental “molecular grammar” of lipidation for condensate engineering, a reductionist approach is essential.

Here, we address this question by systematically characterizing a library of over 80 myristoylated synthetic IDPs (Fig. 1a). By integrating biophysics, molecular simulations, and machine learning, we deconstructed the design space of site-specific lipidation, revealing that cohesive and adhesive forces—which govern material state and miscibility, respectively—can be tuned via distinct sequence features (Fig. 1b). Together, these results establish lipidation as a versatile molecular handle for engineering the mechanical and interfacial properties of biomolecular condensates and illustrate the broader potential of PTMs to encode emergent functions in engineered biomaterials.

**Fig. 1:**
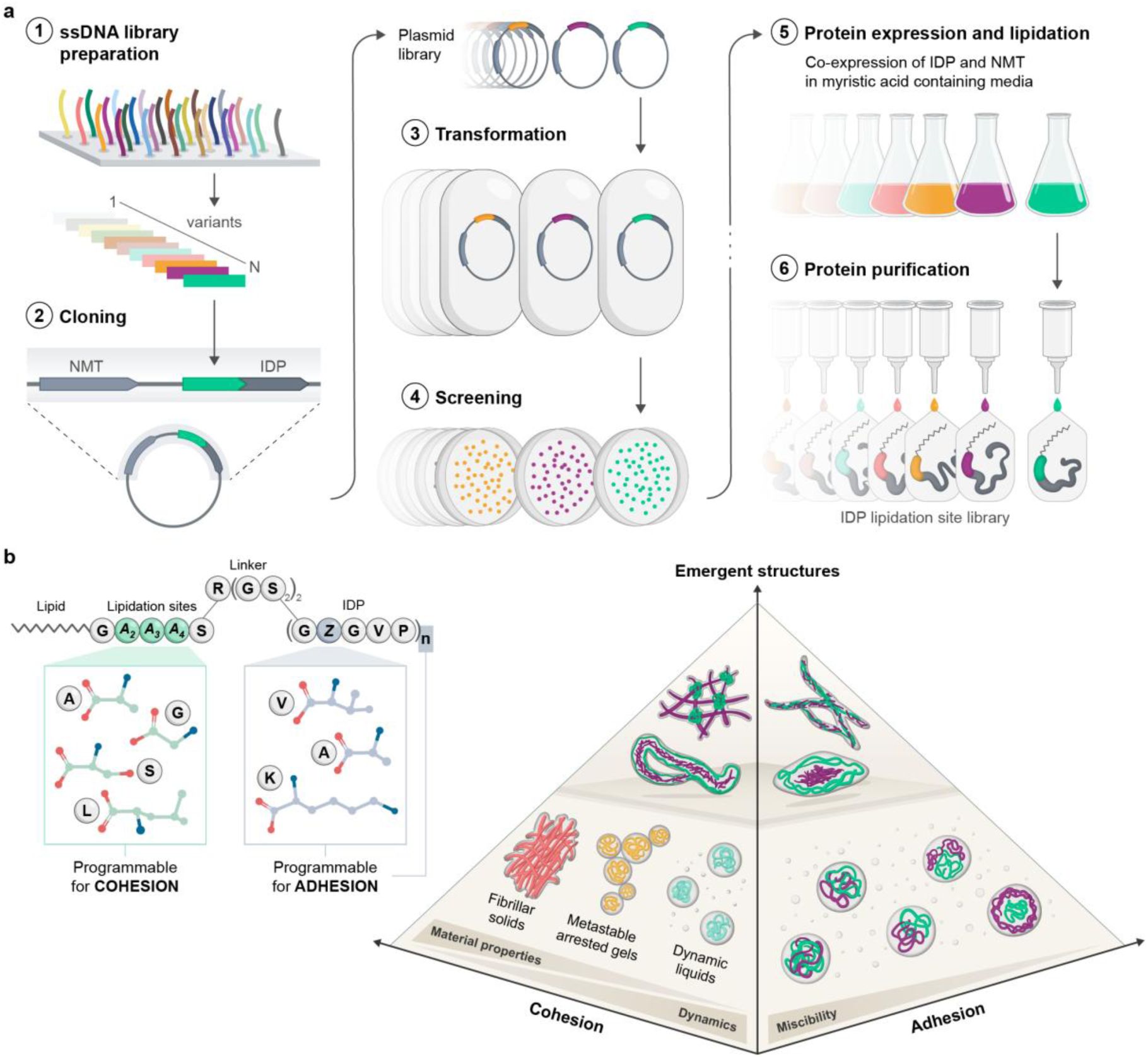
A modular platform for lipidation-based control of biomolecular condensates. **a,** Combinatorial workflow for the construction of the lipidation-site library. A degenerate lipidation-site cassette is fused to an intrinsically disordered elastin-like polypeptide (ELP) scaffold via Gibson assembly. Co-expression with Nmyristoyltransferase (NMT) enables high-throughput recombinant production of myristoylated IDPs bearing systematically varied lipidation-site sequences. **b,** Molecular architecture of lipidated IDPs, highlighting the regions varied in the two-axis design framework. The lipidation-site library explores local sequence space by varying the three residues adjacent to the N-myristoylated glycine (A₂–A₄) using a four-amino-acid alphabet (G, A, S, L). The miscibility library alters the composition (Z) and length (n) of the IDP scaffold. The interplay between the lipid and its local sequence programs cohesion and material properties, whereas the interplay between the lipid and the scaffold composition programs adhesion and miscibility. Tuning these two features yields condensates with emergent hierarchical architectures.

### A Modular System to Define Design Principles of Lipidation

To investigate how lipidation programs the properties of biomolecular condensates, we utilized a modular platform comprising a tunable IDP scaffold fused to a sequence-defined lipidation motif via a short, fixed linker. As the model synthetic IDP, We used elastin-like polypeptides (ELPs)— tropoelastin-derived (GZGVP) repeats, where Z is the variable guest residue—as a tunable intrinsically disordered scaffold, because their LCST phase behavior provides a sensitive readout of condensate formation as a function of temperature and ionic strength.^12^ For lipidation, we chose myristoylation—the acylation of an N-terminal glycine with tetradecanoic acid—catalyzed by N-myristoyl-transferase (NMT), which recognizes the GA_2_A_3_A_4_S motif.^33^ For this library, we varied positions A_2_–A_4_ using a four-letter alphabet—{G, A, S, L}—to create a symmetric design space that explores a diverse range of side-chain properties: flexibility (G), small size/apolarity (A), polar hydrogen-bonding (S), and branched hydrophobicity (L), while remaining compatible with NMT preferences. By installing these sequences onto a fixed ELP scaffold (GVGVP)_30_, we generated a (4^3^) 64-member lipidation-site library to systematically map local sequence context to condensate behavior.

In addition, we constructed a focused Miscibility Library to examine how global scaffold properties—such as chain length and guest residue hydrophobicity—tune the thermodynamic compatibility while keeping the lipidation motif (GLYAS, derived from Arf2 protein) constant. Each construct was produced in both myristoylated and unlipidated form to reveal how the interplay between scaffold properties and lipidation determines condensate miscibility and the formation of multi-phase architectures.

Throughout this manuscript, we use a concise nomenclature to describe each protein variant: m- [SITE]-Z_n_. Here, ‘m’ indicates the N-myristoyl, [SITE] denotes the variable residues of the lipidation site triad, while Z and n are the ELP guest residue and number of repeats, respectively. For example, m-[LAS]-V_30_ refers to a myristoylated GLAS fused to (GVGVP)_30_. Protein sequences, library design and construction details are provided in Supplementary Information, with purity and identity confirmed via HPLC and MALDI-TOF mass spectrometry (Supplementary Tables 1–4; Supplementary Figs. 1–11).

### Lipidation and Lipidation Site Triad Impact Phase Separation Thermodynamics

To quantify the thermodynamic consequences of lipidation on protein condensation, we first mapped the phase behavior of V_30_, and m-V_30_ (the site-specifically myristoylated analogue lacking the lipidation site). Salt-concentration phase diagrams, acquired using a high-resolution microfluidics platform (Supplementary Fig. 12),^34^ revealed that lipidation induces a dramatic shift in the phase boundary to lower salt concentrations, indicating a substantially increased propensity for phase separation (Fig. 2a). Thermal phase diagrams in PBS corroborated this trend (Fig. 2b), demonstrating that lipidation exerts substantial thermodynamic leverage, lowering the transition temperature (*T*_t_) by >20 °C and thereby enhancing LCST phase separation. Beyond this shift, the shape of the temperature-concentration phase boundary for m-V_30_ revealed a more complex, biphasic mechanism. At low protein concentrations, the boundary was steep, consistent with a regime in which predominantly monomeric species must overcome a substantial entropic penalty to demix. However, above ∼ 50 µM, the boundary flattened considerably, indicating a much weaker dependence on bulk protein concentration. This crossover, which is absent in V_30_, is consistent with pre-condensation oligomerization above a critical concentration, wherein self-assembled oligomers (i.e., micelles) serve as nucleating precursors that lower the overall entropic cost of forming a dense phase.^27^

**Fig. 2:**
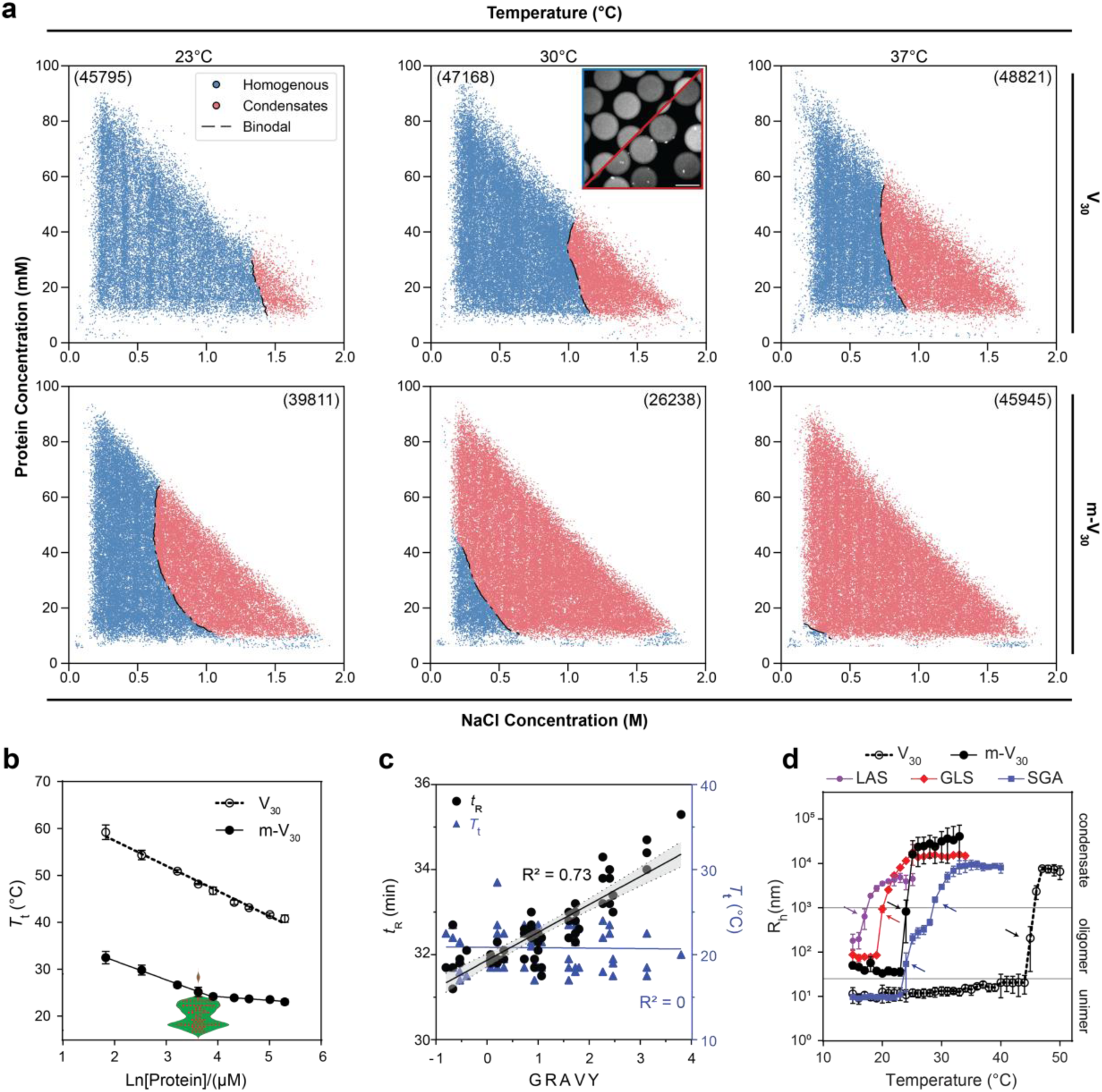
Lipidation enhances phase transition propensity, with secondary modulation from the lipidation site. **a,** Microfluidic phase diagrams of unlipidated V_30_ and myristoylated m-V_30_ as a function of protein and NaCl concentration, obtained via PhaseScan at 25, 30, and 37 °C. Binodals are indicated by dashed lines. Insets show representative micrographs of microdroplets classified as homogenous (blue) or phase-separated (red), scale bar, 100 µm. The total number of microdroplets analyzed is indicated in parentheses. **b,** Partial temperature-concentration phase diagrams of V_30_ and m-V_30_ (symbols/lines, 6–200 µM in PBS) compared with the distribution of transition temperatures (*T*_t_) of the lipidation-site library (violin plot, 50 µM in PBS). Data are mean ± s.d. (n = 3). **c,** RP-HPLC retention time (*t*_R_) and *T*_t_ of the lipidation-site library as a function of theoretical bulk hydrophobicity (GRAVY score) of corresponding A_2_-A_4_. The dashed band represents the 95% confidence interval of the linear regression fit for *t*_R_. **d,** Temperature-dependent hydrodynamic radius (R_h_) profiles for V_30_, m-V_30_, and three representative library variants (measured by DLS at 50 µM protein in PBS). Data are mean ± s.d. (n = 3). Arrows denote *T*_t_, with an additional arrow for m-[SGA]-V_30_ indicating its critical micelle temperature (CMT; Supplementary Fig. 14).

Having shown the dominant role of lipidation, we next investigated whether the local sequence environment—lipidation site triad adjacent to the myristoyl group—could fine-tune this behavior. We measured the *T*_t_ for all 64 variants using variable-temperature dynamic light scattering (VT-DLS) and observed that nearly all variants (63 out of 64) exhibited a lower *T*_t_ than the control construct lacking the triad, with a median shift of -5.0 °C (Fig. 2b, violin plot inset). Crucially, these stabilizations of the dense phase did not correlate with the theoretical bulk hydrophobicity (GRAVY score) of the triad sequence (Fig. 2c), pointing instead to more subtle, position-dependent effects that modulate the cohesive interactions imparted by the lipid (Supplementary Fig. 13). Consistent with this interpretation, VT-DLS showed that variants with lower *T*_t_ tended to form larger oligomers in the soluble phase (Fig. 2d), with polar substitution attenuating cohesion in a subset of variants (e.g., m-[SGA]-V_30_). Together, these results suggest a hierarchical model for the control of phase separation by lipidation. The lipid acts as the primary thermodynamic driver, decreasing protein solubility energetically and promoting oligomerization entropically. Superimposed on this is a secondary layer of control from the lipidation-site triad, which fine-tunes the strength of cohesive forces between chains by modulating the local chemical environment of the anchor and altering the interfacial curvature of pre-condensation assemblies.^35^ This dual framework provides a basis for engineering of condensate properties through site-specific chemical modifications.

### Lipidation Site Sequence Determines Material Properties of Condensed Phase

Having established the phase boundaries of our library, we next investigated how the sequence context of the lipidation site influences the material state of the resulting condensates. Remarkably, we found that minor variations within the lipidation site triad (A_2_-A_4_) were sufficient to drive the emergence of strikingly different material states (Fig. 3a; Supplementary Figs. 15–17): (1) dynamic liquid-like droplets consistent with canonical LLPS (∼53%, 34/64); (2) ordered solid-like fibrous assemblies (38%, 24/64); and (3) metastable viscoelastic intermediates (9%, 6/64). Fluorescence recovery after photobleaching (FRAP) revealed that these classes possessed distinct exchange dynamics, with rapid and nearly complete fluorescence recovery in liquid droplets, negligible recovery in solid fibers, and partial recovery in metastable condensates (Fig. 3b). Corroborating this solid-like character, fibrous assemblies exhibited distinct thermal hysteresis and limited reversibility upon cooling (Supplementary Fig. 18). Similarly, Thioflavin T (ThT) fluorescence further resolved these classes (Fig. 3c; Supplementary Fig. 19), with two distinct populations that mirrored the material states. While droplets remained ThT-negative, fibers displayed high fluorescence consistent with amyloid-like order. Metastable intermediates exhibited intermediate signals that increased over time, paralleling their conversion into fibers (Fig. 3d). Mapping these outcomes across the design space revealed a clear sequence grammar dominated by residues A_3_/A_4_, with the first residue (A_2_) showing minimal impact (Fig. 3e; Supplementary Fig. 20). Specifically, triads with flexible or hydrophobic residues (e.g., Gly, Leu) at both A_3_ and A_4_ almost exclusively formed liquid condensates (14/16), whereas those with small or polar residues (e.g., Ala, Ser) at these positions formed fibers (15/16); mixed combinations produced intermediate behavior, with approximately half of the variants forming droplets. Crucially, these distinct material states were only accessible through site-specific lipidation. Unlipidated constructs (Supplementary Fig. 21) and lipidated proteins lacking the triad (m-V_30_) exclusively formed liquid droplets. Moreover, in-situ myristoylation of unlipidated precursors yielded condensates that were indistinguishable from those formed by the purified lipidated proteins (Supplementary Fig. 22), demonstrating that the observed material states are programmed via the interplay between the lipid and the triad sequence, which collectively governs the cohesive interactions between protein chains.

**Fig. 3:**
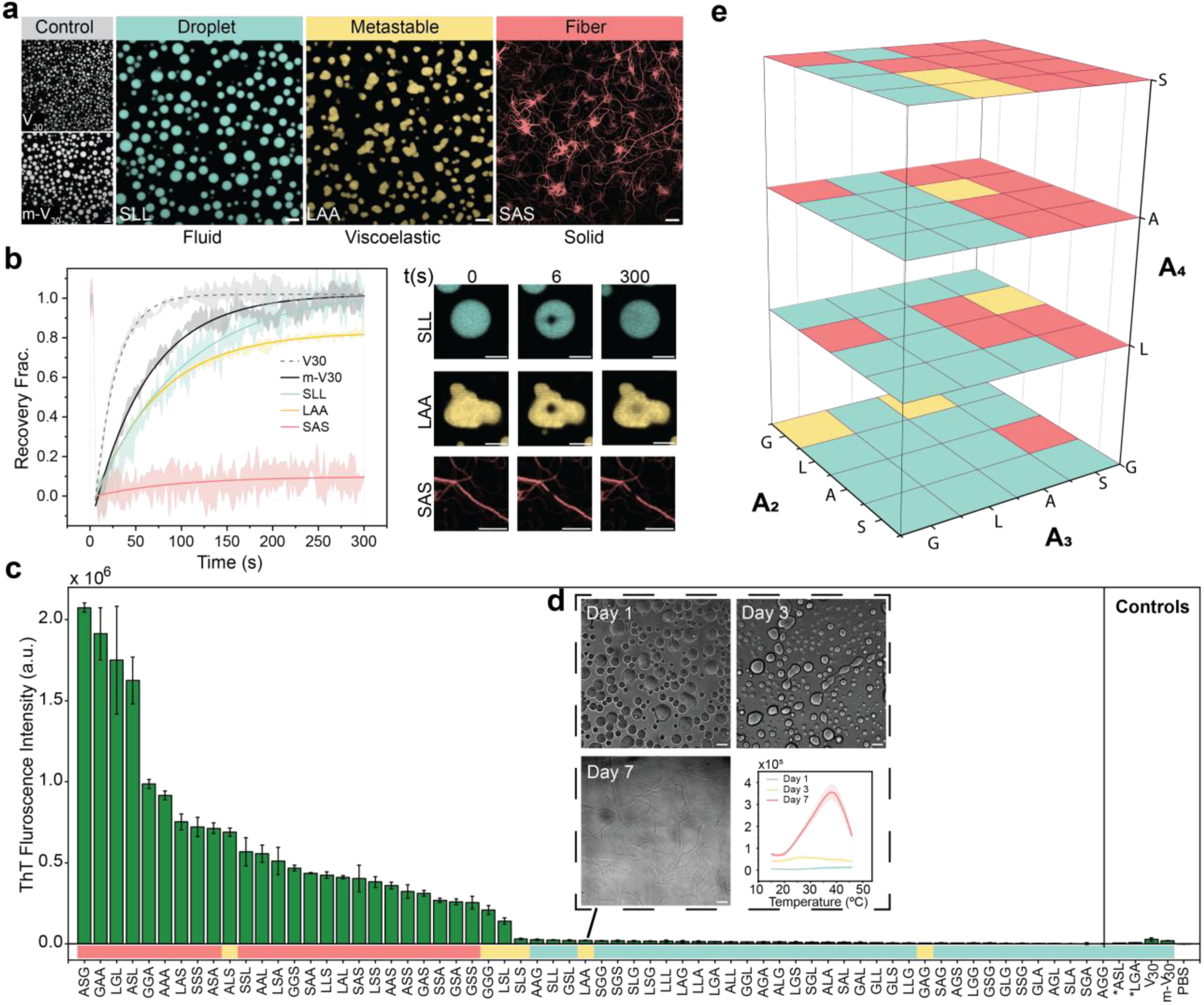
The lipidation site sequence programs a spectrum of condensate material states. **a,** Representative confocal micrographs showing the three distinct material classes identified in the library: dynamic liquid droplets (cyan), metastable viscoelastic gels (yellow), and solid fibrillar (red) structures. Images were acquired at 37 °C with 50 µM protein in PBS, except for V_30_, which required an additional 1M NaCl to trigger phase separation at this temperature. **b,** Quantification of exchange dynamics for representative constructs of each class, V_30_, m-V_30_ using fluorescence recovery after photobleaching (FRAP). Data represent normalized fluorescence intensity (mean ± s.d., n = 3–5), acquired at 37 °C. **c,** Maximum ThT fluorescence for all variants shows a bimodal distribution matching material classes. Values (mean ± s.d., n = 3) correspond to peak fluorescence from samples (50 µM protein, 25 µM ThT) heated from 4–45 °C to induce phase separation. Asterisks indicate non-lipidated control samples. **d,** Maturation of metastable m-[LAA]-V_30_ variant. DIC micrographs and time-dependent ThT emission profiles monitor the transition from liquid to solid states. Images were acquired at 37 °C (50 µM protein in PBS) after incubating the samples for 1, 3, and 7 days at 4 °C. ThT emissions are presented as mean ± s.d., n = 3. **e,** A 3D plot mapping the material state of condensates onto the design space defined by the amino acid identity at each position of the lipidation site, liquid droplet (cyan), fibrillar solid (red), and metastable gels (yellow). Scale bars, 10 µm (**a, d**), 5 µm (**b**).

### Lipidation Amplifies Sequence-Encoded Structural Propensities That Regulate Condensate Properties

To elucidate the molecular mechanism underpinning the interplay between the lipid and triad sequence, we used NMR spectroscopy to probe local structure and dynamics (Fig. 4a–c). We focused on two representative constructs: the fiber-forming m-[SAA]-V_30_ and the droplet-forming m-[SAL]-V_30_. Because alanine appears only at the lipidation site, it provides a direct spectroscopic handle on its local environment. NMR revealed two critical insights. First, the identity of A_4_ modulates the conformational dynamics of the lipidation site (Fig. 4a). The droplet-forming m-[SAL]-V_30_ displayed clear Ala (Hα–Hβ) and lipid-specific NOEs, whereas these signals were attenuated in the fiber-forming m-[SAA]-V_30_, consistent with a rigidified local environment limiting NOE transfer.^36^ Second, although both constructs are globally disordered (Supplementary Fig. 23, 24), differences at the lipidation site subtly bias the ELP conformational ensembles. While both proteins exhibited short-range NOEs characteristic of transient β-turns,^37^ m-[SAL]-V_30_ showed significantly stronger relative intensities, indicative of a more compact, turn-enriched ensemble (Fig. 4b,c). Conversely, m-[SAA]-V_30_ adopted a more extended conformation, which could facilitate the intermolecular associations for fiber nucleation and may concomitantly reduce the population of NMR-visible unimers through the formation of larger pre-condensation assemblies, thereby reinforcing the observed attenuation in NOE signals.

**Fig. 4:**
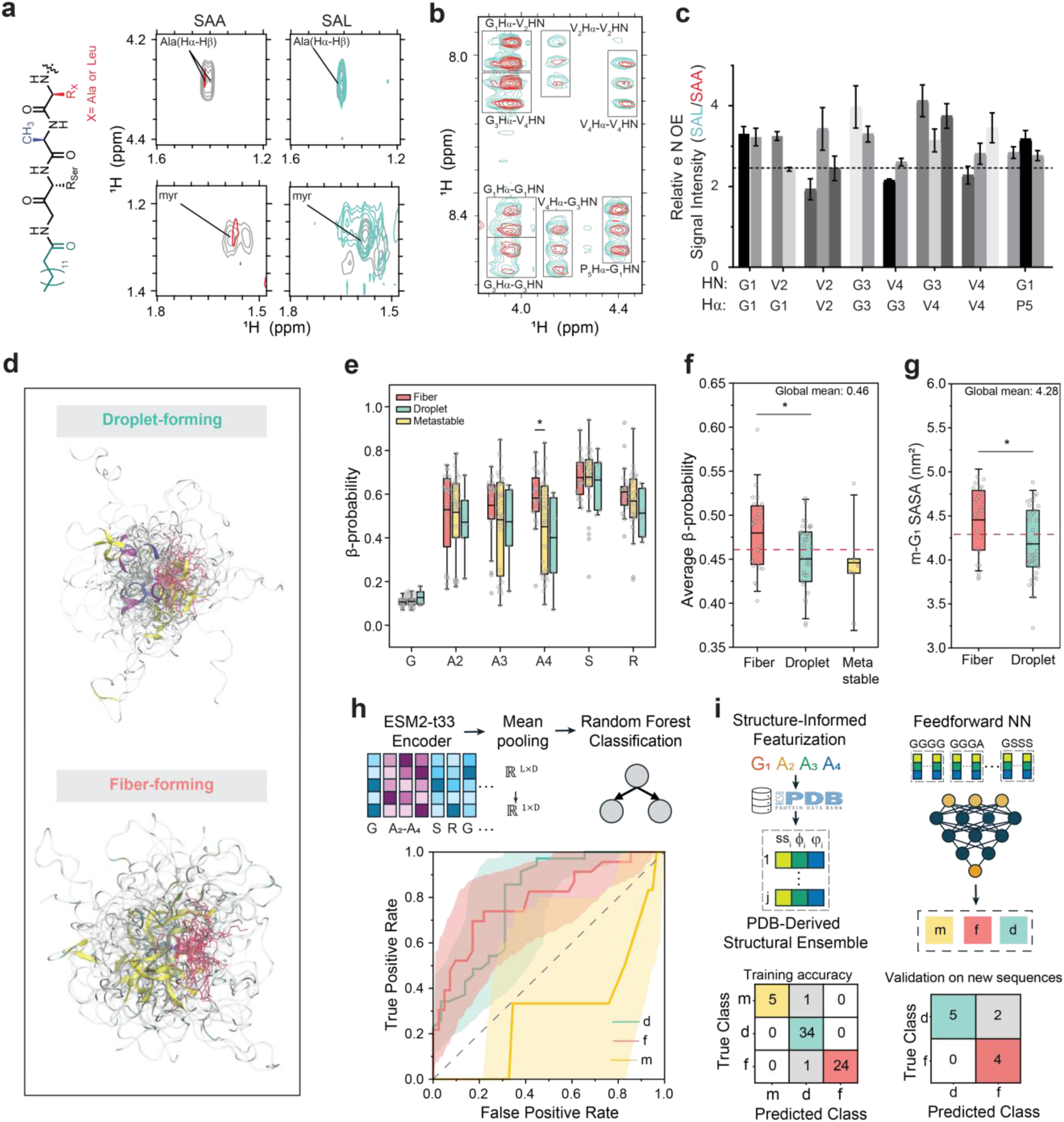
Lipidation amplifies sequence-encoded structural propensities that govern condensed-phase material states. **a,** Chemical structure and overlay of TOCSY (grey) and NOESY spectra of m-[SAA]-V_30_ (fiber-forming, red) and m-[SAL]-V_30_ (droplet-forming, cyan). **b,c,** Analysis of transient β-turns populated by GVGVP repeats. **b,** NOESY regions showing diagnostic Hα–HN correlations for transient β-turns. **c,** Relative NOE intensities (peak height ± uncertainty); dashed line shows the weighted-average NOE ratio (inverse-variance weighting, 1/σ²). **d,** Structural ensembles of droplet-forming (LGG) vs. a fiber-forming (LAL) with five GVGVP repeats, extracted from 1-µs atomistic simulations and aligned on the G1 Cα/N atoms; β-Strands (yellow), helices (purple/blue), and lipid tails (red). **e–g,** Single-chain structural properties derived from atomistic simulations. Box plots show the median (center line), mean (square symbol), quartiles (box limits), and whiskers to 1.5 × the interquartile range. Individual data points are overlaid. **e,** β-structure probability for the first six residues. **f,** Comparison of average β-probability sampled by the first 11 residues (lipidation site+linker). Red line is global mean. **g,** Solvent-accessible surface area (SASA) of the myristoylated glycine, showing increased lipid exposure in fiber-forming sequences compared to droplet-forming sequences. **h,** Sequence-based classification pipeline using ESM2 embeddings and Random Forest. ROC curves show discrimination of droplet (d) and fiber (f) states; metastables (m) are near-random. **i,** Structure-informed neural network trained on secondary structure (*ss*) and dihedral angle (*φ, ψ*) statistics for four-residue motifs derived from PDB. Bottom, confusion matrices for the training (left) and independent validation on new sequences (right).

To define the atomic-level biases that underlie these divergent behaviors, we complemented the NMR measurements with all-atom molecular dynamics simulations. For computational efficiency, we simulated all members of the lipidation site library as truncated constructs (lipidation site + five GVGVP repeats) using the CHARMM36mw force field for 1 μs (Supplementary Figs. 25-33). These simulations revealed that, while the lipidation site remained conformationally flexible, fiber-forming sequences possess a higher intrinsic β-strand propensity across the N-terminal region, particularly at residue A_4_ (Fig. 4d–f), consistent with FT-IR spectra showing a pronounced β-sheet amide I band in fibers that was attenuated in droplet-formers (Supplementary Fig. 34). This local ordering also increased lipid accessibility. The myristoylated glycine was more solvent-exposed in fiber-formers, consistent with increased lipid accessibility that may promote intermolecular association and fibril nucleation (Fig. 4g), whereas it was less solvent-exposed in droplet-formers, consistent with partial lipid burial that may favor intramolecular contacts. Fiber-forming sequences also exhibited a marginal preference for sampling more extended conformations (Rg and E2E), despite broad distributional overlap with droplet-forming variants (Supplementary Figs. 35). Collectively, these data show that the sequence-encoded dynamics at the lipidation site act as a molecular control point linking local flexibility and ensemble bias. By promoting oligomerization and restricting conformational sampling,^35^ lipidation can lower the entropic cost of ordering and amplify these intrinsic structural biases, thereby tuning the emergent condensed-phase material state.

To determine whether local sequence context encodes sufficient information to distinguish condensate material states, we first used latent embeddings from the ESM2 to train a Random Forest classifier (Fig. 4h).^38^ This approach effectively discriminated liquid droplets from solid fibers, but performed poorly on metastable behavior (overall accuracy, 75%; Supplementary Fig. 36), suggesting that metastability is likely governed by kinetics competition between assembly pathways not easily captured by static sequence descriptors. Having established that discriminative information is present in sequence, we next sought a representation that is mechanistically aligned with the conformational “grammar” revealed by NMR and MD analyses. We trained a neural network on physics-informed conformational descriptors—secondary-structure and ϕ/ψ propensities—mined from >200,000 resolved sequence contexts in the Protein Data Bank. This maps the lipidation-site sequence context (the triad and preceding glycine) into a physically defined structural space that reports intrinsic conformational preference (Fig. 4i).^39^ Incorporating these descriptors enabled the model to capture subtle determinants of assembly, reproducing experimentally observed morphologies across the training library with ∼97% accuracy (Supplementary Fig. 37, 38), consistent with learning the biophysical constraints of the dataset. To evaluate generalization beyond the initial screen, we designed and characterized 11 additional constructs containing residues absent from the training set, including bulky hydrophobic amino acids such as Val and Trp. The model correctly predicted 9 of 11 morphologies (82%) (Fig. 4i, Supplementary Fig. 39), supporting both robustness and predictive utility. Together, these results indicate that condensate materiality is dictated by sequence-encoded structural preferences at the lipidation site, providing a foundation for high-throughput in-silico screening/design of lipidation sites as experimental training sets expand and artificial intelligence continues to accelerate the discovery of supramolecular assembly codes.^40,41^

### Lipidation Modulates Miscibility and Architecture of Multi-Component Condensates

Beyond bulk and emergent material properties such as complex aging^42^ and transport kinetics,^43^ biomolecular condensates also derive function from multiphasic architectures that maintain distinct chemical environments (e.g., electric potentials^44,45^ and pH gradients^46^). While the physical principles governing multiphase architectures in charge-driven coacervates have been extensively characterized,^47–50^ whether lipidation can drive these higher-order architectures remains unclear. Having shown that lipidation tunes the cohesive interactions within condensates, we next investigated how it modulates the adhesive (i.e., inter-condensate) interactions that govern the miscibility of multi-component condensates when either one or both IDP scaffolds are lipidated (Fig. 5a). We prepared a focused Miscibility Library comprising of six ELPs with varying lengths and guest residue hydrophobicity: a short, hydrophobic ELP (V_40_), a longer hydrophobic ELP (V_80_), and a long, hydrophilic ELP (V/A/K)_80_, each with and without N-myristoyl group. For this library, we kept the lipidation site constant using a neutral GLYAS sequence (derived from Arf2 protein, referred hereafter as [LYA]). Consistent with the lipidation-site library, myristoylation increased phase-separation propensity, while all variants formed liquid condensates (Supplementary Fig. 40).

**Fig. 5:**
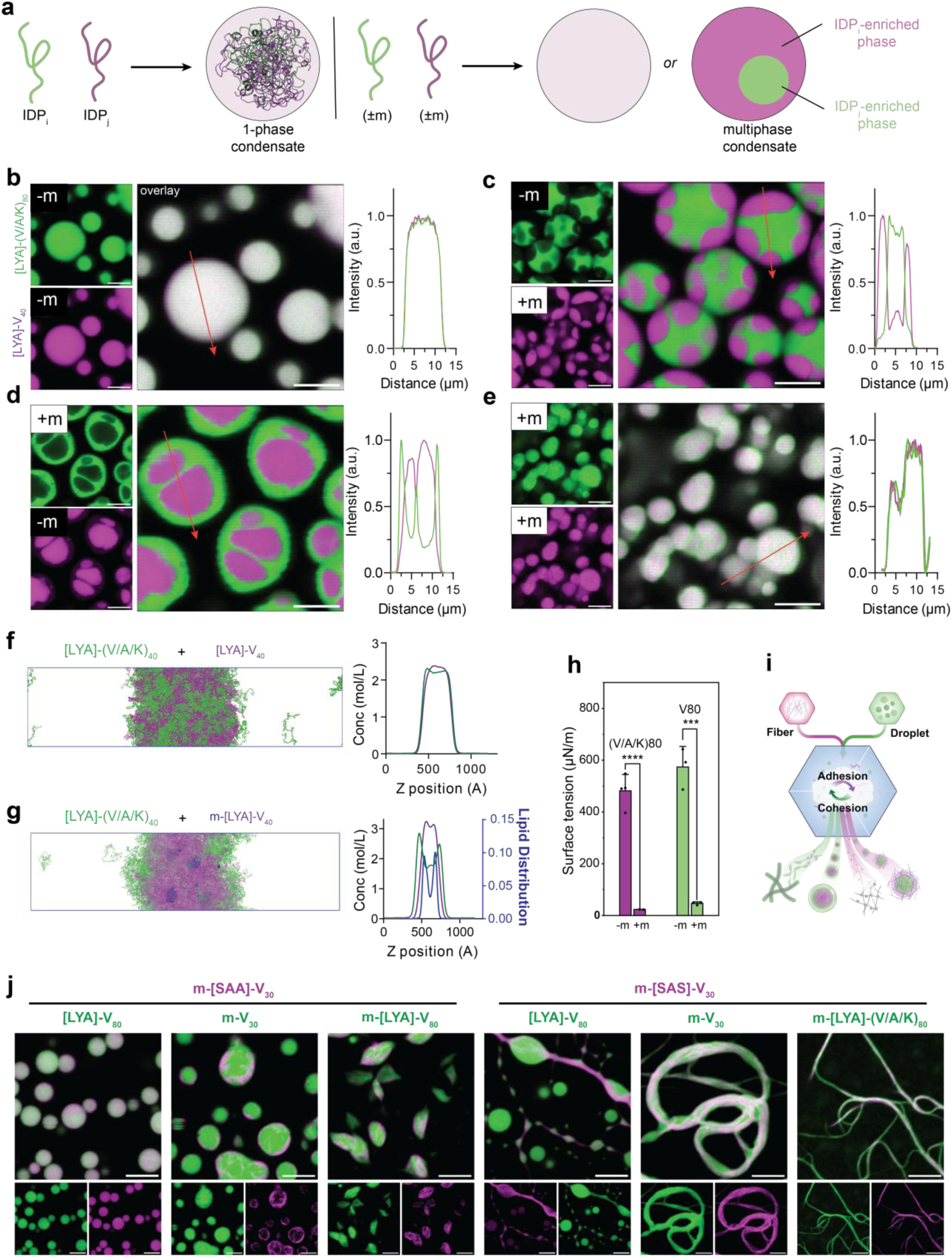
Lipidation tunes inter-condensate adhesion and programs hierarchical architecture by modulating interfacial tension. **a,** Schematic of hierarchical architectures (e.g., homogeneous, core-shell) accessible to binary IDP mixtures. Lipidation pattern alters thermodynamic compatibility between components, determining the final outcome. **b–e,** Confocal micrographs and intensity profiles (along red arrow) of binary mixtures containing [LYA]-(V/A/K)_80_ (green) and [LYA]-V_40_ (magenta). Proteins were mixed at equal mass ratio in 1.7 M NaCl (room temperature). **b,** Unlipidated mixtures form homogenous single-phase condensates. **c,d,** Asymmetric lipidation drives demixing into multiphase core-shell architectures. **e,** Symmetric lipidation of both components restores miscibility. Scale bar, 10 µm. **f,g** Coarse-grained molecular dynamics simulations of binary mixtures. **f**, Unlipidated mixtures show overlapping density profiles, indicating a homogeneous state. **g**, The addition of a myristoyl tail to one component (m-V_40_) induces molecular clustering and formation of an enriched phase. Green/Magenta lines track protein density; lipids are shown in blue. **h,** Quantification of surface tension using micropipette aspiration (MPA). Data are mean ± s.d. of n = 3-4 independent experiments. *P* values were determined by a two-way ANOVA with Tukey’s post-hoc test. **i,** Schematic for generating multi-component complex architectures by mixing fiber-forming (magenta) and droplet-forming (green) constructs. Final morphologies are dictated by the competition between cohesive strength (lipidation site) and adhesive compatibility (IDP scaffold). **j,** Gallery of emergent architectures programmed through wetting transitions: droplets dissolving fibers, capillary-driven encapsulation of fibers within deformed spindle-like droplets,

To investigate how lipidation affects condensate miscibility, we systematically mixed pairwise combinations of library members labeled with distinct fluorophores (AF488, green; Cy5, magenta), revealing lipidation as a potent, switch-like regulator of (de)mixing and internal architecture (Fig. 5b–e). Across this library, a simple rule emerged for compositionally distinct scaffolds: non-lipidated pairs formed a single phase, asymmetric lipidation drove two-phase demixing, and symmetric lipidation restored mixing. For example, non-lipidated (V/A/K)_80_ and V_40_ were fully miscible (partition coefficient (K) ≈ 1; Fig. 5b; Supplementary Fig. 41a), whereas mixtures of m-V_40_ and (V/A/K)_80_ demixed into a condensate with two distinct phases, with each protein preferentially partitioning into one phase (Fig. 5c; K(V/A/K)_80_ ≈ 4.2; K(m-V_40_) ≈ 4.5). A similar effect was observed when the lipid was added to (V/A/K)_80_ scaffold (Fig. 5d). Strikingly, miscibility was restored when both ELPs were myristoylated (K ≈ 1; Fig. 5e).

Coarse-grained simulations using the HPS–Urry model^51^ recapitulated this behavior and provided insight into its molecular origin (Fig. 5f,g; Supplementary Figs. 42, 43): lipidated chains oligomerize through lipid-driven associations, forming dynamic micellar patches within the condensates (Supplementary Fig. 44) that disrupt entropic mixing and drive immiscibility when only one component is lipidated. This behavior generalized across scaffold pairs (e.g., (V/A/K)_80_/V_80_; Supplementary Fig. 41b) and was insensitive to incubation time or mixing order, consistent with a thermodynamic ground state (Supplementary Figs. 45, 46). In contrast, mixtures lacking a scaffold mismatch, such as an ELP with its myristoylated counterpart (e.g., V_40_/m-V_40_) or constructs with matched guest chemistry and different length (e.g., V_40_/m-V_80_), remained fully miscible under all conditions (Supplementary Fig. 41c). co-existing droplets and fibers, droplet forming coaxial sheaths around fibers, and co-assembled fibrillar network. Equal volumes of each protein (0.7 mg/mL in PBS) were mixed (4 °C, 1 h), adjusted to 1 M NaCl to induce phase transition, and matured (1 h, 37 °C) before imaging. Scale bars, 5 μm.

Beyond miscibility, we also observed a striking rule governing the architecture of the multiphase condensates: the lipidated component consistently formed an outer shell that enveloped the nonlipidated phase (Fig. 5c,d; Supplementary Fig. 41b), independent of scaffold hydrophobicity (e.g., m-V_40_ or m-(V/A/K)_80_). This organization is counterintuitive given that myristoylation introduces a hydrophobic moiety that might be expected to favor the core.^52^ Although simulations did not fully reproduce this architecture, they suggest a mechanism in which lipid-driven micellization may effectively increase the hydrophilicity of the lipidated condensate by increasing backbone solvent exposure within micellar patches (Supplementary Fig. 44). To test this hypothesis, we used micropipette aspiration^53^ and contact angle measurements characterize the material properties of the condensates (Fig. 5h, Supplementary Fig. 47, 48). Myristoylation dramatically reduced the surface tension, from 478.7 µN/m to 23.4 µN/m for (V/A/K)_80_, and from 573.7 µN/m to 46.8 µN/m for V_80_, consistent with lipidated proteins acting as polymeric surfactants that accumulate at the condensate–water interface and thereby drive shell formation.^54,55^ Moreover, contact angle measurements (Supplementary Fig. 48) showed that lipidated variants exhibit significantly higher wetting on hydrophilic glass surface, consistent with the increased effective hydrophilicity of the lipidated condensed phase predicted by simulation. Overall, these results establish lipidation as a general strategy to tune condensate interfacial energetics and thereby program hierarchical multiphase architecture without changing the underlying protein sequence. This is significant because reports of PTM-driven demixing remain exceptionally rare and, to our knowledge, are largely restricted to electrostatic, phosphorylation-dependent switches such as FMRP/CAPRIN1 in ribonucleoprotein granules^3^ and SAPAP/PSD-95 in the postsynaptic density.^56^ By contrast, our results show that neutral, hydrophobic PTMs can encode distinct demixing and wetting behaviors, with the potential to expand the design space for engineered multiphasic condensates beyond canonical core–shell architectures.

To explore this design space, we recognized that the mesoscale organization of multicomponent condensates is governed by the interplay between cohesive interactions that drive self-association and adhesive interactions that mediate mixing. Because these forces are regulated by distinct sequence features—lipidation-site grammar and the IDP scaffold—we hypothesized that combining components with different cohesive and adhesive profiles would bias the free-energy landscape toward distinct co-assembly pathways and emergent mesoscale architectures (Fig. 5i). To test this and establish the governing rules, we first examined the high-adhesion limit using fiber- and droplet-forming constructs with matched lipid/IDP backbones, spanning a range of cohesive strengths informed by our lipidation-site heuristics (Fig. 3e). Here, the more cohesive component dictated the material state, spanning templated droplet-to-fiber conversion to droplet-dominated phases (Supplementary Fig. 49a). When adhesive compatibility was reduced by mismatching IDP scaffolds, miscibility controlled whether these interactions produced unified phases (templated conversion or dissolution) or enforced spatial segregation (Supplementary Fig. 49b). Together, these principles define a two-axis design framework for bottom-up programming of multiphasic lipidated condensates structures, yielding emergent architectures exemplified by droplets dissolving fibers, fiber-wrapped droplets, capillary-driven spindle encapsulation, droplet-fiber coexistence, coaxial sheaths, and co-assembled fibrillar networks (Fig. 5j). We anticipate that systematic, combinatorial exploration of this design space will unlock still richer emergent structural complexity.

### Lipidated Condensate Material Properties Regulate Organoid Morphogenesis

Finally, to translate these design rules into functional biomaterials, we took inspiration from elastogenesis, in which elastin coacervates are templated by microfibrils to form elastic fibers.^57^ We reasoned that lipidated ELPs programmed to form fibrous networks could similarly provide biophysical cues that support higher-order tissue morphogenesis and more closely mimic the native ECM.^58–60^ To evaluate this, we engineered hybrid hydrogels by co-assembling lipidated ELPs with Matrigel (MG), a widely used basement-membrane mimic that lacks the fibrillar architecture of interstitial ECM proteins such as collagen-I and instead exhibits a sheet-like laminin-driven organization.^61^ We therefore asked whether introducing condensate-programmed architectures into MG could compensate for this structural limitation and guide enhancements in the emergence of complex, functional tissue architectures.

We first assessed whether temperature-triggered phase separation of lipidated ELPs is orthogonal to MG gelation and whether the resulting micro-architecture alters bulk mechanics. Solutions of either a fiber-forming (m-[AAS]-V_30_) or a droplet-forming (m-[AAG]-V_30_) variant (1% w/v) were mixed 1:1 with MG (1% w/v) at 4 °C and gelled at 37 °C to yield 0.5% MG / 0.5% ELP hybrids. Confocal microscopy confirmed that condensate formation proceeds independently of MG gelation in both the absence (Supplementary Fig. 50) and presence of cells (Fig. 6a), generating composites in which droplets or fibrils were uniformly integrated around embedded intestinal stem cells. Storage modulus (Gʹ) and stress relaxation were then analyzed to decouple their known effects on intestinal organoid morphogenesis^62–67^ from the effect of lipidated ELP microarchitecture. Incorporation of ELPs did not perturb the evolution of the storage modulus; MG concentration remained the dominant contributor to stiffness. While 100% MG formed the stiffest network (G’ = 97 ± 1.3 Pa), the hybrids exhibited intermediate stiffness (29 ± 3.1 Pa for AAG hybrids, 43 ± 4.8 Pa for AAS hybrids), slightly higher than the 50% MG control (22 ± 2.6 Pa) (Fig. 6b). Stress-relaxation profiles were conserved across all conditions (Fig. 6c). Together, these results establish the orthogonality of condensate formation and MG gelation—an operational advantage that enables the programming of distinct mesoscale architectures without significant alterations to the bulk viscoelastic response.

**Fig. 6:**
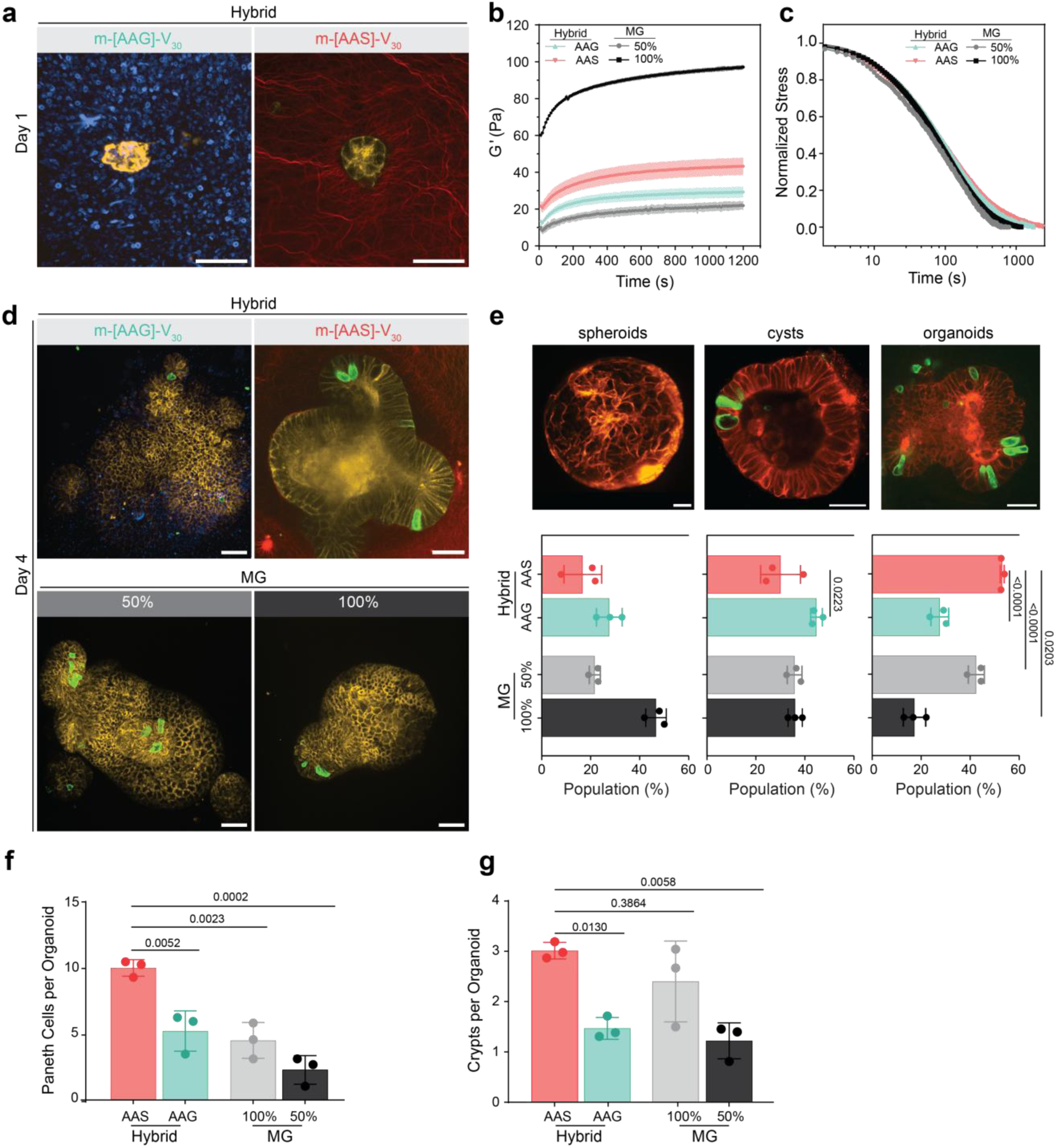
Condensate micro-architecture directs intestinal organoid morphogenesis. **a,** Confocal images of murine Defa4-mTmG intestinal organoids embedded in hybrid composites (0.5 w/v% lipidated IDP+ 0.5 w/v% Growth Factor Reduced Matrigel (MG) at Day 1. The phase-separation of IDP is orthogonal to MG gelation, patterning the microarchitecture of the gel: droplet-forming m-[AAG]-V_30_ forms puncta (pseudo-colored blue), whereas fiber-forming m-[AAS]-V_30_ (red) forms a fibrillar network around cells. **b,c,** Rheological characterization of hybrid composites and MG controls. **b,** Time-dependent evolution of storage moduli (*G’*) measured at 1 Hz, 2% strain) show MG concentration is the primary determinant of bulk storage moduli with secondary effect from IDP microarchitecture. **c,** Normalized stress-relaxation profiles (10% strain) reveal that relaxation kinetics are conserved across all hydrogel conditions. **d,** Representative confocal images of organoids grown in hybrid composites or control gels at Day 4. The fiber-infiltrated composites promote increases in robust symmetry breaking, complex budding, and Paneth cell emergence (green, Defa4-EGFP). **e–g,** Quantitative analysis of organoid morphogenesis and differentiation at Day 4. **e,** Representative structure of intestinal organoid classifications (top; images) and Population distribution of morphological phenotypes (bottom; graphs); the fibrillar AAS matrix yields the highest proportion of mature, budded organoids (∼53%). **f,** Quantification of Paneth cell differentiation based on Defa4-EGFP fluorescence. **g,** Quantification of crypt formation per organoid. Data represent mean ± s.d. from n = 3 independent biological experiments (each with three technical replicates). *P* values are determined by one-way ANOVA with Tukey’s post-hoc test, and comparisons to AAS are shown. Scale bars, 30 µm.

We next evaluated how these architectures influence the development of murine intestinal organoids (Fig. 6d; Supplementary Fig. 51). Organoids cultured in 100% MG frequently stalled as spheroids or cysts, and the droplet-forming m-[AAG]-V_30_ hybrid showed similar limitations. In contrast, the fiber-forming m-[AAS]-V_30_ composites markedly accelerated morphogenesis, yielding the highest fraction of mature, budded organoids (∼53%; Fig. 6e). This enhancement extended to functional niche formation: m-[AAS]-V_30_ supported abundant, crypt-localized Paneth cells and the highest numbers of crypts and Paneth cells per organoid (Fig. 6f,g).

Softening the matrix with 50% MG increased crypt number relative to 100% MG and the droplet-forming hybrid but did not achieve the level of Paneth cell specification observed in the fibrillar composites. Thus, while mechanical softening can partially enhance cryptogenesis, the superior performance of the fibrillar condition cannot be attributed solely to changes in stiffness and instead points to a specific contribution from the supramolecular architecture. Together, these results establish lipidation-site grammar as a handle for biomimetic matrix design, enabling fibrillar microstructure that bias tissue morphogenesis and stem-cell niche organization in vitro.

## Conclusion

In this work, we establish that lipidation can be used to program the formation, material state, and architecture of engineered condensates. Our systematic analysis supports a thermodynamically grounded model in which the lipid-driven oligomerization promotes condensation while simultaneously amplifying the sequence grammar at the attachment site. By restricting conformation sampling, this oligomerization lowers the entropic cost for ordering and enables the lipidation site to form persistent cohesive contacts that determine whether a condensate behaves as a dynamic liquid, an arrested gel, or an ordered fibrillar solid. In parallel, the emergent amphiphilicity of the lipidated IDP modulates the interfacial tension (adhesion) with the solvent and with co-existing condensed phases. Because these two axes are controlled by orthogonal genetic elements (lipidation site + IDP scaffold), this platform provides a foundation for systematic, combinatorial exploration of PTM-inspired design rules for soft, adaptive biomolecular materials.

The ability to program fibrillar micro-architectures via lipidation has clear implications for tissue engineering, synthetic biology, and artificial cells. By recapitulating a key feature of the native ECM, these engineered fibrillar networks can provide instructive cues to guide multicellular self-organization process,^68,69^ as illustrated by morphogenesis of intestinal organoid in our study.

Furthermore, because lipidation can both promote phase-separation and program the material state of condensed phase, it offer a powerful strategy for engineering synthetic intracellular and abiotic condensates and materials.^70–73^ From a biological perspective, our findings suggest that lipidation could be a widespread, yet currently underappreciated, mechanism for the regulation of cellular condensates. The exquisite sensitivity of the system—where a minor change in sequence context drives a shift in material state from a liquid to a solid— makes it an ideal candidate for a dynamic regulatory switch. We further speculate that reversible lipidations (e.g., palmitoylation) could dynamically remodel condensate properties at membrane interfaces in response to signaling inputs or environmental stress. Future work will be needed to explore this possibility in living cells and to determine the full scope of PTM-based regulation of biomolecular condensates.

## Methods

Detailed experimental procedures are provided in the Supplementary Information.

## Data availability

All relevant data are included in the manuscript. Source data are provided with this paper.

## Code availability

The code used for phase scan image processing is available at https://github.com/rqi14/PhaseScan. Full implementation details of sequence-based transfer learning using ESM2 are available at https://github.com/Rob-Scrutton/LipidationPredictions. Scripts for high-throughput analysis of protein library HPLC and MALDI data, as well as the Fiji macro for confocal image analysis, are available at https://github.com/ZhiweiHuang-source/lipidation-site.

## Acknowledgements

We thank A. A. Aljiboury, A. L. Dovciak, and A. E. Patteson for assistance with microscopy, FTIR spectroscopy, and rheology. This work was supported by the National Institutes of Health (R35GM142899 (D.M.); R35GM144045(J.C.); R35GM138097(A.B.); K99/R00 DK135907(M.R.B.); R35GM147027(Z.S.)) and the National Science Foundation (CAREER 2146168 to D.M.). Additional support was provided by the Pew Charitable Trusts (A.B.), the European Research Council (DiProPhys 101001615 to T.P.J.K.), and a FLAD R&D@USA research internship grant (L.A.P.).Organoids were provided by the CU Anschutz Organoid and Tissue Modeling Shared Resource at the University of Colorado Anschutz Medical Campus, which is supported, in part, by the Gates Institute, Diabetes Research Center (grant P30 DK116073), and Cancer Center Support Grant (P30CA046934). pET-24a-ELP[V-30] was a gift from Ashutosh Chilkoti (Addgene plasmid # 67014).

## Author contributions

D.M. conceived the project, designed the experiments, and directed the research. Z.H., M.M.A., and M.S. generated and characterized the lipidation-site library. Z.H. characterized the miscibility library, performed biophysical measurements (FT-IR, MPA, rheology), and led the data analysis. M.A. performed ThT assays. M.S. and A.B. conducted NMR experiments and analyzed results with some sample isotopically labelled proteins prepared by L.A.P.. H.C.S. performed and analyzed molecular simulations under the supervision of J.C.. T.A. performed PhaseScan experiments, and R.S. conducted ESM2/RF classification under the supervision of T.P.J.K.. S.V. and D.T. performed organoid culture experiments with guidance from M.R.B.. A.G. and L.M. conducted fuel-driven LLPS experiments. J.W. and Z.S. contributed to the design and analysis of MPA experiments. Z. Q. and A.V.J. contributed to the development of the FNN model and subsequent analysis. A.L.A. assisted with technical characterization including MALDI-TOF and DLS. All authors discussed the results and commented on the manuscript.

## Competing interests

A patent application related to this work has been filed.

## Notes

https://github.com/rqi14/PhaseScan

https://github.com/Rob-Scrutton/LipidationPredictions

https://github.com/ZhiweiHuangsource/lipidation-site

