## Supplementary Information for "Lipoengineering of Biomolecular Condensates Controls Material Properties and Multiphase Hierarchy to Guide Organoid Morphogenesis"

SUPPORTING INFORMATION

Zhiwei Huang,[1] Md Mahbubul Alam,[1] Mahtab Shokri,[1] Harshavardhan C. Savitrinarayana,[2] Sisila Valappil,[1] Tanushree Agarwal,[3] Rob M. Scrutton,[3] Laiba Maryam,[1] Asma Gulzar,[1] Jinying Wang,[4] Dominic J. Tigani,[5] Liege A. Pascoalino,[1,†] Akshay V. Jadhav,[6] Albert L. Adhya,[1] Alaji Bah,[7] Zhao Qin,[6] Zheng Shi,[4] Michael R. Blatchley,[5] Jianhan Chen,[2] Tuomas P.J. Knowles,[3] and Davoud Mozhdehi[1*]

[1] Department of Chemistry, Syracuse University, Syracuse, NY 13244, USA.
[2] Department of Chemistry, University of Massachusetts, Amherst, MA 01003, USA
[3] Department of Chemistry, Centre for Misfolding Disease, University of Cambridge, Cambridge, UK
[4] Department of Chemistry and Chemical Biology, Rutgers, The State University of New Jersey, NJ 08854, USA.
[5] Department of Biomedical and Chemical Engineering, Syracuse University, Syracuse, NY 13244, USA
[6] Department of Civil and Environmental Engineering, Syracuse University, Syracuse, NY 13244, USA.
[7] Department of Biochemistry and Molecular Biology, SUNY Upstate Medical University, Syracuse, NY 13210, USA.
*

### 1. Materials

DNA oligonucleotides were purchased from Integrated DNA Technologies (Coralville, IA). Chemically competent *Escherichia coli* strains (NEB5α and BL21(DE3)), Q5 High-Fidelity DNA Polymerase, NEBuilder HiFi DNA Assembly master mix, and kits for plasmid miniprep, gel extraction, NdeI, XhoI, KLD Enzyme Mix and PCR cleanup were purchased from New England Biolabs (Ipswich, MA).

Myristic acid, ampicillin, kanamycin, phosphate-buffered saline (PBS) tablets, HPLC-grade solvents (acetonitrile, isopropanol, acetone), trifluoroacetic acid (TFA), anhydrous dimethylformamide (DMF), dimethyl sulfoxide (DMSO), buffer components (sodium chloride, sodium phosphate dibasic anhydrous, ammonium chloride, potassium phosphate monobasic), yeast extract, tryptone, agarose, agar powder, ethanol, biotin, calcium chloride anhydrous, magnesium sulfate anhydrous, dextrose anhydrous, thiamine hydrochloride, Alexa Fluor 488 cadaverine, TrypLE Express, Alexa Fluor 488 NHS Ester, and Cy5 NHS Ester were purchased from Thermo Fisher Scientific (Waltham, MA). Isopropyl β-D-1-thiogalactopyranoside (IPTG) was obtained from Gold Biotechnology (St. Louis, MO). Fluorescent dyes including AZDye 488 Cadaverine, AZDye 647 Amine were procured from Vector Labs (Newark, CA). HFE-7500 oil containing 1.2% (w/v) fluorosurfactant was purchased from RAN Biotechnologies (Beverly, MA).

Apomyoglobin, cytochrome c, adrenocorticotropic hormone (ACTH) fragment 18–39, bradykinin fragment 1–7, sinapinic acid, α-cyano-4-hydroxycinnamic acid (CHCA), LC-MS grade ammonium bicarbonate, iron(III) chloride, deuterium oxide, Pluronic F-127, Sigmacote, myristoyl-CoA, *N*-acetyl-L-cysteine, and valproic acid sodium salt, ProteoMass Protein MALDI-MS Calibration Kit, 1-ethyl-3-(3-dimethylaminopropyl)carbodiimide (EDC) and N-hydroxysuccinimide (NHS) were purchased from Sigma-Aldrich (St. Louis, MO). Consumables such as syringe filters, membrane filters (0.22 µm, cellulose acetate and PVDF), ZipTip C18 pipette tips, and 3 mm NMR tubes were also from Sigma-Aldrich. $^{15}$N-ammonium chloride was purchased from Cambridge Isotope Laboratories (Tewksbury, MA).

Protein electrophoresis materials (Mini-PROTEAN TGX 5–20% stain-free precast gels, 4× Laemmli sample buffer, protein standards, SimplyBlue SafeStain, and 50× TAE buffer) were procured from Bio-Rad Laboratories (Hercules, CA). Sequencing-grade modified trypsin was obtained from Promega Corporation (Madison, WI). 384-well glass-bottom plates (#1.5 cover glass) were purchased from Cellvis (Mountain View, CA). Deionized water was purified using a Milli-Q system (Millipore, Molsheim, France).

*Organoid and Cell Culture Materials:* The mouse intestinal organoid line Defa4-mTmG (Reporter: *Defa4*^CreNeo/+; *Rosa26*^mTmG) and R-spondin conditioned medium were obtained from the CU Anschutz Organoid and Tissue Modeling Shared Resource (Aurora, CO). Growth Factor-Reduced Matrigel (phenol red-free, LDEV-free) and cell strainers (40 µm) were purchased from Corning (Corning, NY). Intestinal cell base medium (BM), Dulbecco's Phosphate-Buffered Saline (DPBS), and fetal bovine serum (FBS) were procured from Gibco (Waltham, MA). Recombinant murine epidermal growth factor (EGF) and recombinant murine Noggin were purchased from PeproTech (Rocky Hill, NJ). Small molecules Y-27632 and Thiazovivin were purchased from Selleckchem (Houston, TX), DNase I from Roche (Indianapolis, IN), and CHIR99021 from Tocris Bioscience (Minneapolis, MN). All reagents and chemicals were used as received without further purification.

### 2. Methods

#### 2.1 Library Design and Construction

To systematically probe local sequence effects on the condensate properties of N-myristoylated proteins, a focused library was designed for the consensus motif G-$A_2$-$A_3$-$A_4$-S. The amino acid alphabet at the variable positions ($A_2$–$A_4$) was restricted to {G, A, S, L} to exclude residues known to inhibit NMT catalysis (e.g., charged side chains, proline, and bulky aromatics),[1,2] while maintaining diverse physicochemical properties.

To implement this design, a two-step semi-combinatorial assembly strategy was employed wherein the $A_2$ position was fixed to create distinct parent vectors, while positions $A_3$ and $A_4$ were randomized using degenerate KBG codons (K = G/T; B = C/G/T).[3] Parent vectors encoding a fixed $A_2$ residue upstream of the $V_{30}$ ELP scaffold were constructed by three-fragment Gibson assembly. An in-house pETDuet_NMT vector[4] was linearized (*NdeI*/*XhoI*) and assembled with the $V_{30}$ gene (PCR-amplified from Addgene #67014) and a synthetic gBlock encoding the fixed lipidation motif using NEBuilder HiFi DNA Assembly Master Mix (50 °C, 30 min; 1:2:2 molar ratio). Library diversity was subsequently introduced by linearizing the parent vectors via inverse PCR (Q5 Hot Start High-Fidelity Master Mix) using primers flanking the $A_3$/$A_4$ sites (Supplementary Table 1). The linearized backbones were confirmed by electrophoresis (1% agarose gel) and purified via PCR clean up kit and assembled with pools of ssDNA bridge oligonucleotides containing the degenerate KBG codons using NEBuilder HiFi DNA Assembly Master Mix (50 °C, 1 h; 1:200 vector:insert molar ratio). The assembled plasmids were transformed into chemically competent *E. coli* NEB5α cells and screened by Sanger sequencing. Any sequences absent from the pooled library were synthesized individually using a deterministic ssDNA bridge to ensure 100% coverage.

To evaluate the contribution of myristoylation, control vectors lacking the NMT gene (ΔNMT) were generated. Inverse PCR was performed on select library constructs using Q5 Hot Start High-Fidelity Master Mix and primers flanking the NMT coding region. The resulting amplicons were circularized using the KLD Enzyme Mix, transformed into *E. coli* NEB5α cells, and verified by nanopore sequencing.

#### 2.2 Protein Expression and Purification

A single colony of freshly transformed *E. coli* BL21(DE3) cells harboring the desired expression vector was cultured in 2×YT medium supplemented with ampicillin (150 mg $L^{-1}$) at 37 °C with shaking (220 rpm). Protein expression was induced at an optical density ($OD_{600}$) of ~0.7 by the addition of IPTG (1 mM). For lipidated constructs, the culture medium was supplemented with myristic acid (100 µM; added from a 100 mM stock in DMSO) at the time of induction. Following 3 h of expression at 37 °C, cells were harvested by centrifugation (5,000 × *g*, 30 min, 4 °C) and stored at −80 °C.

Proteins were isolated by previously reported organic extraction protocol[5] with slight modifications. Briefly, cell pellets were resuspended in isopropanol (4 mL/g of wet biomass), dispersed via vortexing and sonication. The lysate was clarified by centrifugation (16,000 × *g*, 5 min, 25 °C). The supernatant was collected, adjusted to 70% (v/v) acetonitrile, and incubated on ice for 1 h to induce precipitation of proteins. The precipitate was recovered by centrifugation (16,000 × *g*, 10 min, 4 °C), resuspended in 50% ethanol (5 mL/g of wet cell weight), and filtered (0.22 µm PVDF). The crude product was purified by semi-preparative reverse-phase high-performance liquid chromatography (RP-HPLC) to >95% purity by using a Prominence HPLC system (Shimadzu) with a PDA detector, equipped with a Phenomenex Jupiter® C18 column (5 µm, 300 Å, 250 x 10 mm), at a flow rate of 4.2

mL min$^{-1}$. The mobile phases were (A) water with 0.1% (v/v) TFA and (B) acetonitrile with 0.1% (v/v) TFA. The gradient consisted of a 5-minute isocratic run at 0% B, followed by a linear increase to 90% B over 40 min (injection volume: 4 mL). In this gradient system, the lipidated fragments eluted around 67% B while the unlipidated ones around 54% B. Chromatograms were recorded at 210 nm. Eluted protein fractions were pooled, flash frozen in liquid nitrogen, and lyophilized (FreeZone 4.5, Labconco, Kansas City, MO). Lyophilized proteins were stored at −20 °C.

For NMR studies, proteins were expressed in M9 minimal medium to achieve uniform $^{15}$N-labelling. Starter cultures were prepared in 2×YT medium containing the appropriate antibiotic (ampicillin: 150 mg L$^{-1}$; kanamycin: 45 mg L$^{-1}$ for control plasmids) and used to inoculate 1 L of M9 minimal medium containing $^{15}NH_4Cl$ (3 g L$^{-1}$; >98% $^{15}$N, Cambridge Isotope Laboratories) as the sole nitrogen source. The medium was prepared from nitrogen-free salts (43.5 mM $Na_2HPO_4$, 22 mM $KH_2PO_4$, 8.6 mM NaCl) and supplemented with glucose (4 g L$^{-1}$), thiamine (10 mg L$^{-1}$), biotin (10 mg L$^{-1}$), $MgSO_4$ (2 mM), $CaCl_2$ (4 μM), and $FeCl_3$ (10 μM). Culture growth, induction, and protein purification were performed as described above, with the exception that post-induction expression proceeded overnight (~14 h). $^{15}$N isotopic incorporation was verified by Matrix-assisted laser desorption/ionization time-of-flight (MALDI-TOF) mass spectrometry by comparing the observed molecular weight with the theoretical mass calculated using the NIST Mass and Fragment Calculator.[6]

#### 2.3 Protein Characterization

##### 2.3.1 Analytical HPLC

RP-HPLC was performed on a Shimadzu LC-2030 system equipped with a UV–vis detector. Separation was achieved on a Phenomenex Jupiter C18 column (5 μm, 300 Å, 250 × 4.6 mm) maintained at ambient temperature. Samples (50 μM) were filtered through 0.22 μm PVDF syringe filters prior to analysis. The mobile phases were (A) water with 0.1% (v/v) trifluoroacetic acid (TFA) and (B) acetonitrile with 0.1% (v/v) TFA. A linear gradient of 0–90% solvent B over 40 min was used at a flow rate of 1.0 mL min$^{-1}$ (injection volume: 50 μL). Chromatograms were recorded at 210 nm.

##### 2.3.2 MALDI-TOF Mass Spectrometry

MALDI-TOF mass spectrometry were acquired on a Bruker Microflex LRF system equipped with a microScout ion source and a 337 nm nitrogen laser, operated in linear positive-ion mode.

*Intact Protein Analysis:* Samples were prepared using the dried-droplet method with sinapinic acid (10 mg mL$^{-1}$ in 50% acetonitrile, 0.1% TFA) as the matrix. Protein solutions (50 μM in PBS) were mixed 1:1 (v/v) with matrix and spotted onto the target plate. Signal intensity was optimized via serial dilution on-plate. Spectra with a signal-to-noise (S/N) ratio >10 were calibrated externally using cytochrome *c* ($[M+H]^+$ = 12,361.96 Da) and apomyoglobin ($[M+H]^+$ = 16,952.27 Da). Data were analyzed using Bruker flexControl 3.4. Theoretical and observed masses are summarized in Supplementary Table 3.

*Tryptic Digest Analysis:* To confirm N-terminal myristoylation, constructs were digested with trypsin to release the lipidated N-terminal fragment. Protein samples (25 μM) were incubated with sequencing-grade trypsin (1:20 v/v ratio) in 100 mM ammonium bicarbonate (pH 7.8) for 3 h at 37 °C. Peptides were desalted and concentrated using C18 ZipTips according to the manufacturer's protocol. The eluted peptides were analyzed by MALDI-TOF mass spectrometry using CHCA as the matrix. Spectra were calibrated against bradykinin fragment 1–7 ($[M+H]^+$

= 756.3997 Da) and adrenocorticotropic hormone fragment 18–39 ($[M+H]^+$ = 2,464.1989 Da). Theoretical and observed masses are summarized in Supplementary Table 4.

#### 2.4 Fluorescent Labeling of Proteins

Proteins were fluorescently labeled at a substoichiometric level (<1%) to enable condensate visualization. Depending on construct design, one of the following two chemoselective labeling strategies was employed. Control experiments confirmed that fluorophore identity and labeling sites did not measurably affect condensate physicochemical properties.

*Amine-Reactive Labeling (NHS-Ester):* Constructs with accessible primary amines were labeled using NHS-ester activated fluorophores (e.g., Alexa Fluor 488 or Cy5 NHS Ester). Protein stocks (10 mg $mL^{-1}$) were prepared in 0.2 M $NaHCO_3$ (pH 8.2) and mixed with fluorophores at a feed ratio of 0.2:1 (dye:protein) overnight at 4 °C with gentle agitation.

*Carboxyl-Reactive Labeling (EDC/NHS Coupling):* For constructs lacking accessible amines, C-terminal carboxyl groups were activated via EDC/NHS chemistry followed by coupling of amine-functionalized fluorophores (e.g., AZDye 488 Cadaverine or AZDye 647 Amine). Proteins were dissolved in MES buffer (0.2 M, pH 5.0), and carboxyl groups were activated by the addition of 1-ethyl-3-(3-dimethylaminopropyl)carbodiimide (EDC, 5 equivalent) and N-hydroxysuccinimide (NHS, 10 equivalent) at 4 °C. After 20 min, the pH was adjusted to 7.5–8.0 using NaOH. Amine-functionalized fluorophores were added at a feed ratio of 0.15:1 (dye:protein). The reaction proceeded overnight at 4 °C.

For both strategies, unreacted dye was removed by precipitating the protein with NaCl (2 M) followed by centrifugation (16,000 × g, 5 min, 40 °C). The resulting pellet was resuspended in water and further purified via preparative RP-HPLC and lyophilized for storage at −20 °C.

#### 2.5 Phase-Boundary

##### 2.5.1 Phase Scan

*Automated phase diagram mapping*. Phase diagrams were acquired using the droplet-based microfluidic PhaseScan platform as previously established.[7,8] Protein inputs consisted of 110 µM $V_{30}$ or m-$V_{30}$ in 1× PB, labeled with AZDye 488 (1%) to enable fluorescence detection. Phase separation was triggered by mixing the protein stream with a salt solution containing 2 M NaCl buffered in 1× PB barcoded with AlexaFlour 647 (Thermo Fisher Scientific). These aqueous components were loaded into the device alongside HFE-7500 oil. Flow rates were managed by a pressure-driven controller (Fluigent) to generate combinatorial concentration gradients.

For 2D phase diagrams, the total aqueous flow rate was maintained at 60 µL $h^{-1}$ (individual components varied 5–50 µL $h^{-1}$). The oil phase was injected at 50–80 µL $h^{-1}$ to ensure stable droplet formation. Droplets were imaged using an epifluorescence microscope (Cairn Research) with a 10× objective (Nikon CFI Plan Fluor, NA 0.3).

*Data analysis.* A custom Python pipeline was used to process raw images (code is available at GitHub (https://github.com/rqi14/PhaseScan). Droplets were identified, and their fluorescence intensities were converted into protein concentrations using calibration curves derived from reference standards. Phase separation was determined by the visual presence of condensates within the droplets. The final phase diagrams were plotted using the average concentrations for each classified condition.

##### 2.5.2 Dynamic Light Scattering

Dynamic light scattering (DLS) measurements were performed on a Zetasizer Nano (Malvern Panalytical, UK) equipped with a 173° backscattering detector. Protein samples were prepared at 50 µM in PBS (pH 7.4) at 4 °C and filtered (0.22 µm PVDF). Temperature-dependent changes in hydrodynamic radius ($R_h$) were monitored using a thermal ramp with 1 °C increments and 2 min equilibration at each step. The phase transition temperature ($T_t$) was defined as the midpoint between the two consecutive temperatures, showing the largest increase in $R_h$.

##### 2.5.3 Critical Salt Concentrations

Critical salt concentrations (CSC) were determined via turbidity measurements using a BioTek Synergy H1 microplate spectrophotometer (BioTek, Winooski, VT). Experiments were performed in 384 well glass-bottom plates. Protein solutions were prepared at three concentrations (0.66, 1.00, and 1.34 mg/mL) in PBS. For each protein condition, NaCl was titrated across 16 concentrations (in 0.025 M increments) spanning the expected phase transition boundary. The final sample volume was maintained at 30 µL per well. Plates were equilibrated at 25 ± 1 °C with orbital shaking for 5 min prior to measurement. Turbidity was monitored by absorbance at 350 nm ($A_{350}$). The CSC was defined as the NaCl concentration at which the turbidity surpassed a threshold of $A_{350} = 0.10$.

##### 2.5.4 Variable-Temperature Turbidimetry

Temperature-composition phase diagrams and thermal reversibility profiles were obtained using a Cary 300 UV–vis spectrophotometer (Agilent Technologies, Santa Clara, CA) equipped with a Peltier temperature controller. Sample turbidity was monitored at 350 nm while the temperature was ramped at a rate of 1 °C min$^{-1}$.$^{-1}$. For reversibility measurements, samples were subsequently cooled at the same rate.

#### 2.6 Microscopy

##### 2.6.1 Substrate Treatments

The glass bases of microplates were treated with Pluronic F-127 to minimize surface wetting following established protocols.[9] First, the glass surfaces were immersed in 1 M KOH for 10 min, rinsed thoroughly with DI water, and dried under a nitrogen stream. Subsequently, 50 µL of Sigmacote was dispensed into each well, incubated for 10 min, and rinsed with isopropanol. Finally, prior to imaging, wells were blocked with a 0.5% (w/v) Pluronic F-127 solution for 1 h, followed by five washes with PBS. Protein solutions (30 µL) were then introduced into the wells for analysis.

##### 2.6.2 Confocal and DIC Imaging

Imaging was performed on a Zeiss LSM 980 laser scanning confocal microscope (Carl Zeiss, Jena, Germany) equipped with a 63×/1.4 NA Oil C Plan-Apochromat objective and controlled by ZEN Blue 3.3 software.

*Differential Interference Contrast (DIC):* Samples (50 µM protein in PBS) were illuminated with a 639 nm laser, and transmitted light was detected using a photomultiplier tube (T-PMT). Image acquisition parameters (2048 × 2048 pixels) were standardized across all samples, with gain and offset adjusted for optimal contrast.

*Fluorescence Imaging:* The focal plane was positioned ~4 µm above the substrate surface. General condensate morphology was assessed in Airyscan mode (3072 × 3072 pixels; three regions per specimen). For high-resolution analysis, selected condensates were imaged in Airyscan mode at 1024 × 1024 pixels with 6× optical zoom. Laser power and gain settings were held constant across all samples.

#### 2.7 Fluorescence Recovery After Photobleaching (FRAP)

Samples were equilibrated for at least 1 h prior to analysis to ensure steady-state conditions. FRAP experiments were performed on spherical condensates (8–15 μm diameter) with the focal plane set ~4 μm above the substrate. Photobleaching was performed on a circular region of interest (ROI, diameter = 1 μm) using 60% laser power (488 nm or 639 nm, corresponding to the fluorophore excitation maximum). A minimum of three distinct condensates ($N \geq 3$) were analyzed per condition.

Image acquisition spanned 5 min, with frames captured every 2 s. Bleaching was initiated after the third pre-bleach frame. To correct for sample drift, image stacks were aligned using the StackReg plugin in Fiji (ImageJ).

Recovery curves were fitted to the single-exponential model: $I(t) = a - b \cdot e^{\frac{-t}{\tau_f}}$, where $I(t)$ is the normalized fluorescence intensity, $\tau_f$ is the fluorescence recovery time constant, and $a$ and $b$ are fitting parameters corresponding to the mobile and immobile fractions, respectively.[10]

#### 2.8 Thioflavin T (ThT) Assay

ThT fluorescence assays were performed using a QuantStudio 3 Real-Time PCR System, following a previously published protocol.[4,11] A fresh stock of ThT (500 μM in PBS) was prepared and protected from light. Reaction mixtures (25 μL) were prepared with 48 μM protein and 24 μM ThT (2:1 molar ratio). Fluorescence was monitored using the FAM filter set ($\lambda_{ex} = 450$ nm, $\lambda_{em} = 510$ nm) while the temperature was ramped from 5 °C to 60 °C at a rate of 1 °C min$^{-1}$. All measurements were performed in triplicate.

#### 2.9 Multiphase Condensates

To assess the effect of lipidation on condensate miscibility, stock solutions of all miscibility-library proteins were prepared at 3 mg/mL in PBS. Two protein solutions were combined at a 1:1 volume ratio, mixed thoroughly (~10 pipetting cycles), and phase separation was triggered by adding 5.1 M NaCl (final conditions: 2 mg mL$^{-1}$ total protein, 1.7 M NaCl). Samples were incubated in the dark for 30 min prior to imaging.

Emergent structures formed by fiber and droplet-forming constructs (Fig. 7) were prepared by mixing stock solutions (0.72 mg mL$^{-1}$ in PBS) at a 1:1 volume ratio. The mixture was incubated at 4 °C for 1 h. Phase separation was triggered by adding 10 μL of 5.1 M NaCl to 50 μL of mixture , followed by gentle mixing. Samples were incubated at 37 °C for 1 h to allow maturation.

#### 2.10 Partition Coefficient Analysis

Partition coefficients were calculated from fluorescence images containing ≥ 20 droplets. For each channel, a binary mask was generated via Otsu thresholding in Fiji to define the dense phase. The dilute phase area for Protein 1 was defined by the mask of Protein 2, and vice versa. The partition coefficient (P) of Protein 1 was calculated as P1 = Idense/Idilute, where Idense and Idilute represent the mean pixel intensities of the fluorophore in the dense and dilute phases, respectively.

#### 2.11 Biophysical Characterization of Condensates

##### 2.11.1 Micropipette Aspiration (MPA) Setup

MPA experiments were performed on a Nikon Ti2A inverted fluorescence microscope (Nikon, Japan) equipped with a motorized stage and two motorized four-axis micromanipulators (PatchPro 5000, Scientifica, UK). Micropipettes were fabricated from borosilicate glass capillaries (KWIK-Fil TW100-4; World Precision Instruments [WPI], Sarasota, FL) using a pipette puller (PUL-1000, WPI). The tips were cut to an inner diameter of 3–5 μm and bent to an angle of 40° using a microforge (DMF1000, WPI). The proximal end of each micropipette was connected to a FlowEZ microfluidic flow controller (Fluigent SAS, France) with the vacuum source was ranging from 0−69 mbar. The micropipette holder was positioned such that the bent tip was parallel to the imaging plane. The zero-pressure point ($P_0$) was calibrated before each experiment following established protocols,[12] defined as the minimal pressure change (<2 Pa) required to produce observable directed flow near the tip.

##### 2.11.2 Sample Preparation

Phase separation was initiated by mixing 20 μL of protein stock (3 mg mL$^{-1}$) with 10 μL of NaCl at a concentration adjusted to reach the specific critical salt concentration for each variant (Supplementary Figure 40). The sample was mixed by pipetting up and down approximately five times. Prior to measurement, small condensates were coalesced into larger droplets (~10 μm diameter) using an optical trap (Tweez305, Aresis, Ljubljana, Slovenia). An individual condensate was positioned at the pipette tip and captured by applying a negative pressure of 5 Pa. To minimize nonlinear dissipation effects, the volume of condensate aspirated into the pipette was maintained at ≤5% of the total condensate volume.

##### 2.11.3 Static Surface Tension and Contact Angle

For surface tension ($\gamma$) measurements, the aspiration pressure was increased stepwise until the condensate deformation matched the inner radius of the micropipette. Pressure increments were set to 40 Pa for non-lipidated condensates and 5 Pa for lipidated condensates. The critical pressure difference ($\Delta P$) required for this deformation was used to calculate surface tension via the Laplace equation:

$$\gamma = \frac{\Delta p}{2} / (\frac{1}{R_p} - \frac{1}{R_c})$$

Here $R_p$ is the pipette radius, $R_c$ is the condensate radius. Contact angles were estimated by gently adhering ~10 μm condensates to the outer wall of the micropipette. Static shapes were imaged and analyzed in Fiji using circular/elliptical fitting to determine the droplet footprint and meniscus geometry.

##### 2.11.4 Viscosity and Dynamic Surface Tension Measurements

Following static surface tension determination, an aspiration pressure ($\Delta P_a$) exceeding the critical Laplace pressure ($2\gamma/R_p$) was applied to induce flow of the condensate into the micropipette. The condensate was aspirated until the aspiration length ($L_p$) reached approximately 40 μm. Subsequently, stepwise changes in aspiration pressure were applied to probe the deformation rate under varying shear stresses, maintaining $L_p$ between 5—40 μm. Time-lapse images were acquired at 0.5 Hz using a 60× objective and an ORCA-Flash4.0 sCMOS camera (Hamamatsu, Shizuoka, Japan). Geometric parameters including $L_p$, $R_p$ and $R_c$ were quantified using Fiji. The dynamic viscosity (η) was derived from the linear relationship between the applied pressure and the rate of deformation ($d(L_p/R_p)^2/dt$), assuming a Newtonian fluid model.[13]

$$\Delta P_a = 4\eta\, d\frac{\dot{L}_{\mathrm{p}}^2}{R_{\mathrm{p}}^2}/dt + P_\gamma$$

$P_\gamma$ is the pressure needed to overcome the condensates surface tension.

#### 2.12 Nuclear Magnetic Resonance (NMR) Spectroscopy

All NMR experiments were performed with a protein concentration of 50 µM in PBS buffer containing 2.5% $D_2O$. Samples were measured in 3 mm tubes with a final volume of 200 µL. All spectra were recorded on a Bruker 800 MHz spectrometer equipped with a cryogenic probe. The spectra were processed with NMRpipe[14] and analyzed with SPARKY[15].

Gradient-selected 2D $^{1}H$–$^{15}N$ HSQC spectra were acquired using the Bruker pulse program fhsqcf3gpph at 15, 25, 35, and 45 °C. The $^{1}H$ spectral width was 16.0 ppm (12821 Hz) with 2048 points in the direct ($t_2$) dimension. The indirect $^{15}N$ dimension contained a 64 $t_1$ increments covering the amide $^{15}N$ region (100–135 ppm). A relaxation delay of 2.0 s and 80 scans per $t_1$ increment was used.

2D $^{1}H$–$^{1}H$ TOCSY spectra were acquired at 15 °C using the Bruker pulse program mlevgpph19.UW. The spectral width in the $^{1}H$ dimension was set to 11.97 ppm (9579 Hz), with 2048 complex points collected in the direct ($t_2$) dimension and 256 increments in the indirect ($t_1$) dimension. The carrier was positioned at 4.70 ppm, and a relaxation delay of 2.0 s with 24 scans per $t_1$ increment was used. An isotropic mixing time of 80 ms was applied.

2D $^{1}H$–$^{1}H$ NOESY spectra were acquired at 15 °C using a gradient-selected NOESY pulse sequence (Bruker pulse program noesygpph19.UW). The spectral width in both dimensions was set to 11.97 ppm (9579 Hz) with 2872 points collected in the direct ($t_2$) dimension and 256 increments in the indirect ($t_1$) dimension. The $^{1}H$ carrier was positioned at 4.70 ppm. A relaxation delay of 3.0 s and 16 scans per $t_1$ increment were used. An NOE mixing time of 100 ms was employed.

#### 2.13 Fourier Transform Infrared (FTIR) Spectroscopy

Attenuated total reflectance (ATR)-FTIR spectra were acquired at room temperature using a Nicolet iS50 spectrometer (Thermo Scientific, Waltham, MA) equipped with an iD7 ATR accessory (diamond crystal). Protein samples (10 mg $mL^{-1}$ in deuterated PBS) were allowed to phase separate at room temperature and incubated for 1 h prior to measurement. Spectra were collected over the range of 400-4000 $cm^{-1}$with a resolution of 0.2 $cm^{-1}$, averaging 64 scans per sample after background subtraction.

#### 2.14 Fuel-driven Liquid-Liquid Phase Separation

Unlipidated proteins ([LLL]-$V_{30}$, [SSS]-$V_{30}$, [ASL]-$V_{30}$, [SAS]-$V_{30}$ and [SLL]-$V_{30}$) were expressed from plasmids lacking the NMT gene and purified as described in section 2.2.

Recombinant hexahistidine-tagged NMT (6×His-NMT) was expressed and purified following established protocols with slight modifications.[16] Briefly, *E. coli* BL21(DE3) cells harboring the expression plasmid were cultured in 2×YT medium (containing 100 µg $mL^{-1}$ ampicillin) at 37 °C. Upon reaching an $OD_{600}$ of 0.7, the temperature was lowered to 28 °C, and expression was induced with 1 mM IPTG for 14 h. Cells were harvested, resuspended in PBS, lysed by sonication, and clarified by centrifugation. The lysate was supplemented with 10 mM imidazole and purified via immobilized metal affinity chromatography using standard buffers (50 mM phosphate buffer, 300 mM NaCl, pH = 7.4; 10 mM imidazole for wash buffer; and 300 mM imidazole for elution buffer). Eluted fractions were dialyzed against water, lyophilized, and stored at −80 °C.

NMT-catalyzed myristoylation and the associated phase separation were monitored in reaction mixtures containing 100 µM unlipidated ELP, 200 µM myristoyl-CoA, and varying concentrations of NMT (0.025–20 µM). Reactions were assembled on ice in PBS and incubated at 30 °C with gentle rotation (80 rpm). Condensate formation was assessed by DIC microscopy. To quantify conversion, aliquots were analyzed after 1 h by RP-HPLC. The kinetics of phase separation were monitored by measuring turbidity (absorbance at 350 nm) in 384-well plates (100 µL per well). Measurements were taken at 1-min intervals for 120 min at 30 °C. The plate was shaken for 15 s prior to each reading to maintain suspension, and measurements were performed without a lid to prevent condensation artifacts. All conditions were analyzed in quadruplicate.

#### 2.15 Fabrication and Characterization of ELP-Matrigel Composite Hydrogels

##### 2.15.1 Synthesis of ELP–Matrigel Composites

Composite hydrogels were prepared using fiber-forming (m-[AAS]-$V_{30}$) and droplet-forming (m-[AAG]-$V_{30}$) constructs labeled with AZDye 647 amine. Lyophilized proteins were resuspended in pre-chilled Dulbecco's phosphate-buffered saline at a concentration of 10 mg $mL^{-1}$ and rotated at 4 °C overnight to ensure complete dissolution. For confocal imaging, proteins were labeled with AZDye 647 Amine as described in Section 2.5.

ELP solutions were mixed with Growth Factor-Reduced Matrigel at a 1:1 volume ratio. Fiber-forming constructs were mixed directly, whereas droplet-forming constructs were first incubated at 37 °C for 30 min to induce the formation of larger condensates prior to mixing. Composite droplets (10 µL) were then polymerized in inverted 48-well plates at 37 °C for 15 min.

##### 2.15.2 Characterization of ELP-Matrigel hydrogels

*Confocal Imaging:* The composite morphology was verified using a Zeiss LSM 980 Airyscan 2 Multiplex microscope. Images were acquired with a 40× objective to visualize the distribution of ELP fibers or droplets within the Matrigel matrix.

*Rheology:* Rheological measurements were performed on a Kinexus Ultra+ rheometer (Malvern Panalytical, Malvern, UK) equipped with a Peltier-controlled lower plate and an 8 mm parallel plate geometry (PC08). Experiments were conducted at 37 °C with a gap size of 0.400 mm. To minimize evaporation, a ring of mineral oil was applied around the sample perimeter.

*Sample Loading:* ELP solutions (10 mg $mL^{-1}$) were gently combined with Matrigel at a 1:1 volume ratio using slow pipette aspiration to avoid bubble entrapment. Droplet-forming construct (AAG) was pre-incubated at 37 °C for 30 min prior to mixing to induce condensate formation, whereas fiber-forming construct (AAS) was mixed directly without pre-incubation. A 25 µL aliquot of the mixture was dispensed onto the lower plate, the geometry was lowered, and the sample was allowed to equilibrate for 1 min. Matrigel-only controls were prepared identically.

*Time Sweep (Gelation):* Gelation kinetics were monitored in oscillatory strain-controlled mode (1 Hz, 2% strain) with data points collected every 10 s. Gelation was considered complete when the storage modulus ($G'$) exhibited <5% relative change over a continuous 5 min window.

*Stress Relaxation:* Following the time sweep, stress relaxation was characterized using a small step strain ($\gamma_0 = 10$; rise time $\approx 0.10$ s). The strain was held constant for 40 min at 37 °C while maintaining zero normal force (±0.02–0.05 N tolerance). The time-dependent shear stress $\sigma(t)$ was recorded and converted to the relaxation modulus $G(t) = \sigma(t)/\gamma_0$. Characteristic relaxation times were extracted from smoothed curves after removing inertial artifacts from the first 2 s.

#### 2.16 Mouse Intestinal Organoid Culture

##### 2.16.1 Organoid Maintenance and Passaging

Mouse intestinal organoids derived from the Defa4-mTmG line were maintained in growth factor–reduced Matrigel at 37 °C in a humidified incubator with 5% $CO_2$. Cultures were grown in Intestinal Cell Base Medium (BM) supplemented with R-spondin–conditioned medium (5% v/v), recombinant murine EGF (50 ng $mL^{-1}$), and recombinant murine Noggin (100 ng $mL^{-1}$), referred to as ENR medium.

*Mechanical Passaging:* Matrigel domes were recovered in cold BM and pelleted (200 × *g*, 5 min). Organoids were fragmented by mechanical trituration (~45 strokes with a P1000 pipette tip) and re-pelleted. Fragments were resuspended in cold Matrigel (final concentration 8 mg $mL^{-1}$), dispensed as 10 µL droplets into pre-warmed 48-well plates, polymerized at 37 °C for 15 min, and overlaid with ENR medium.

*Single-Cell Dissociation:* Organoids were dissociated using TrypLE Express supplemented with DNase I, *N*-acetyl-L-cysteine (1 mM), and Y-27632 (10 µM). Incubation consisted of two 4-min cycles at 37 °C with trituration. The reaction was quenched with BM containing 10% FBS, and the suspension was filtered through a 40 µm cell strainer. Cells were counted and embedded in Matrigel at a density of $5 \times 10^5$ cells $mL^{-1}$. Polymerized domes were overlaid with ENR medium supplemented with CHIR99021 (3 µM), valproic acid (1 mM), *N*-acetyl-L-cysteine (1 mM), and Thiazovivin (2.5 µM) to support single-cell outgrowth.

##### 2.16.2 Encapsulation of Organoids in ELP–Matrigel Hybrid Hydrogels

Single cells were pelleted (400 × g, 5 min) and resuspended in the ELP-Matrigel (1:1; w/w) composites. 5 mg (50%, 1:1 Matrigel:PBS) and 10 mg (100%, Matrigel alone) of Matrigel were used as controls. Approximately 5000 cells were seeded per 10 µL droplet, deposited into the center of pre-chilled 48-well plates, and polymerized for 15 min at 37 °C. Following gelation, cultures were overlaid with complete ENR medium supplemented with CHIR99021 (3 µM), valproic acid (1 mM), N-acetyl-L-cysteine (1 mM), and Thiazovivin (2.5 µM) for 48 h to promote single-cell survival. The medium was replaced with ENR medium alone to induce differentiation and crypt formation for the next 48 hours. Samples were fixed in 3.2% paraformaldehyde and 0.1% glutaraldehyde in PBS for 20 min at room temperature. Samples were washed three times with PBS (5 min each) prior to imaging.

##### 2.16.3 Imaging and Analysis

Organoid growth was monitored daily using an Echo Rebel microscope (Discover Echo Inc.). Quantitative analysis of organoid morphology was performed on the final day using automated image analysis software (TellU).[17] Organoids that were not detected by the automated pipeline were counted manually to ensure accurate classification of the different morphologies present in each condition. For high-resolution assessment of crypt formation and epithelial morphology (e.g., Paneth cells), samples were fixed and imaged using a Dragonfly confocal microscope (Andor Technology)) to acquire 3D z-stack images. In mTmG-Defa4 organoids, EGFP-positive paneth cells were localized to crypt regions and quantified per crypt and per organoid from 3D z-stacks using Imaris (Oxford Instruments, Abingdon, UK).

#### 2.17 Atomistic Simulations

##### 2.17.1 Simulation Setup

Sixty-four protein constructs with distinct lipidation sites were prepared for atomistic molecular dynamics (MD) simulations. Initial peptide coordinates were generated using PeptideBuilder[18], and N-terminal myristoylation of glycine was introduced using an in-house CHARMM script. Each fully extended initial structure was subjected to a 100 ps NVT simulation in vacuum at 500 K, to generate a partially collapsed and randomized conformation. The resulting structures were solvated in a $10 \times 10 \times 10$ nm$^3$ cubic box with water and 0.15 M NaCl (Supplementary Figure 25). Energy minimization was performed using the steepest descent algorithm until forces were below 1000 kJ mol$^{-1}$ nm$^{-1}$, followed by a 125 ps NVT relaxation at 303.15 K.

Production simulations were carried out for 1.0 μs in the NPT ensemble with a 2 fs timestep, maintaining temperature and pressure at 303.15 K and 1 bar using V-rescale and C-rescale[19] coupling, respectively. Long-range electrostatics were treated with PME[20], van der Waals interactions employed a force-switch scheme (1.0–1.2 nm), and all bonds involving hydrogen atoms were constrained using LINCS[21].

##### 2.17.2 Trajectory Analysis

Production trajectories were analyzed using MDAnalysis[22] and GROMACS (2024)[23] after discarding the first 100 ns as equilibration. The end-to-end distance (E2E) was computed as the Euclidean distance between the N-terminal myristoylated glycine and the C-terminal $C_\alpha$ atoms. The radius of gyration was calculated using all backbone heavy atoms together with the aliphatic carbon atoms (C1–C14) of the myristoyl group. The solvent-accessible surface area (SASA) of the myristoylated glycine (GLYM) and Ramachandran angle distributions were computed using GROMACS.

##### 2.17.3 Convergence Metrics

Convergence of structural properties (Rg, E2E, SASA, φ, ψ) was assessed by comparing block-averaged distributions from the first (100–550 ns) and second (550–1000 ns) halves of each trajectory (Supplementary Figure 26–34). For each observable, normalized histograms were constructed for two-time windows, and convergence was quantified using relative root-mean-square deviation (rel-RMSD) between corresponding histograms:

$$rel_{RMSD} = \frac{\sqrt{\left(\frac{1}{N}\right)\Sigma\left(h_1^i - h_2^i\right)^2}}{\left\|\frac{(h_1 + h_2)}{2}\right\|_2 + \epsilon}$$

Here, $h_1^i$ and $h_2^i$ denote the heights of the $i$th bin in the first and second-half histograms, respectively; $N$ is the total number of bins; $\|\cdot\|_2$ represents the Euclidean norm; and $\varepsilon$ (set to $10^{-8}$) is a small constant introduced to avoid division by zero. $rel_{RMSD}$ is reported as a percentage.

β-strand propensities were extracted from Ramachandran distributions using φ/ψ filter defining β regions (–160° ≤ φ ≤ –50°, 90° ≤ ψ ≤ 180°). Residue-wise β-strand probabilities were computed for residues 1–11 for each sequence and grouped by class labels (Fiber, Droplet, Metastable). Statistical comparisons were performed using box-and-strip plots, showing the per-residue distribution of β-strand probabilities across classes.

#### 2.18 Coarse Grained Simulations

##### 2.18.1 Simulation Setup

Coarse-grained molecular dynamics simulations were performed using the HPS–Urry model[24] implemented in the openABC package,[25] in which each amino acid is represented by a single bead with residue-specific hydropathy-based Lennard–Jones interactions. System was built using openABC package. Simulations were carried out on binary mixtures of 200 [LYA]-$(V/A/K)_{40}$ and 200 [LYA]-$V_{40}$ chains in a 20 × 20 × 120 $nm^3$ rectangular slab under periodic boundary conditions, with initial configurations prepared to be in a condensed geometry to accelerate equilibration and to improve sample dense-phase behavior.

All simulations were performed under the NVT conditions at 300 K using a Langevin integrator with a friction coefficient of 0.01 $ps^{-1}$ and a 10 fs timestep for 3.0 μs each. Nonbonded interactions were truncated at a cutoff distance equal to four times of maximum Lennard–Jones σ parameter in the system.

##### 2.18.2 Model and Parameter Choices for Lipid Tail and Lipidation Site Beads

Lipidation was modeled explicitly by representing the myristoyl group as a linear chain of four coarse-grained beads, corresponding to an effective ~3:1 mapping of carbon atoms per bead. The representation preserves the length and excluded volume characteristics of the C14 aliphatic chain at a coarse-grained level. Each lipid bead was assigned Leu-like Lennard–Jones σ parameters, reflecting the aliphatic, nonpolar character of lipid hydrocarbons and the structural and chemical similarity to the LEU side chain.

Consistent with this mapping, lipid-amino acid interactions were treated identically to Leu–amino acid interactions, such that lipid beads act as generic hydrophobic moieties. Lipid–lipid interactions were assigned a slightly higher interaction strength than standard Leu–Leu interactions (0.184 vs 0.128 kcal $mol^{-1}$; ≈0.31 $k_BT$), reflecting the stronger effective cohesion of hydrocarbon tails while remaining within the native interaction range of the HPS–Urry model and preserving reversible association.

Atomistic simulations together with NMR and FTIR analysis indicated that residues at and near the lipidation site exhibit an increased propensity for β-like conformations and enhanced inter-chain contacts. To consider this lipidation-induced cooperativity in the coarse-grained model without explicitly enforcing secondary structure, residues designated as lipidation-site/linker residues (LYAS) were assigned a modestly strengthened mutual attraction relative to standard aromatic interactions. Accordingly, interactions between LYAS-LYAS residues were set to 0.384 kcal $mol^{-1}$ (≈0.65 $k_BT$), which mimics additional backbone-mediated intermolecular β-sheet hydrogen bonding interactions. This choice selectively enhances lipidation-mediated protein–protein association and promotes β-sheet-like stabilization as an emergent property, while avoiding irreversible aggregation or artificial structural constraints.

#### 2.19 Machine Learning

##### 2.19.1 Sequence-based Transfer-Learning

Protein sequence embeddings were generated using ESM2 (esm2_t33_650M_UR50D), a 650-million parameter protein language model (Lin et al., 2023).[26] Embeddings were extracted from the final transformer layer, yielding 1280-dimensional representations per residue. Two feature encoding strategies were evaluated: (1) mean-pooled representations averaged across all residues, and (2) lipid site-specific representations extracted at positions 2, 3, and 4, and concatenated to form a 3,840-dimensional feature vector, corresponding to the variable lipidation sites.

Structural outcomes (droplet, fiber, metastable) were classified using Random Forest models (100 trees, max depth 8, balanced class weights) with 5-fold stratified cross-validation. Feature standardization was applied within each fold to prevent data leakage. Performance was evaluated using Receiver Operating Characteristic (ROC) curves computed using a one-versus-rest strategy, with 95% confidence intervals for the Area Under the Curve (AUC) estimated via bootstrap resampling (n=1000). Analysis was implemented in Python using scikit-learn and PyTorch. Full implementation details are available at [https://github.com/Rob-Scrutton/LipidationPredictions].

##### 2.19.2 Structure-informed Neural Network

Feature vectors were generated by searching the PDB for resolved structural occurrences matching the lipidation site motif (the variable triad plus the preceding glycine, $GA_2A_3A_4$) using an automated pipeline.[27,28] For each match, residue-level statistics—specifically secondary-structure propensities[29] and backbone dihedral angle distributions (φ/ψ)—were extracted as conformational priors. These structural priors were integrated with primary sequence to create a composite feature space, yielding 204,651 synthetic physically grounded entries. Classification was performed using a feedforward neural network (FNN). The optimal topology was determined via Bayesian Optimization within the MATLAB classification module, minimizing training cross-entropy loss. The final architecture consisted of three hidden layers (264, 171, and 10 neurons) with Sigmoid activation functions. Weights were initialized using the Glorot scheme with zero-initialized biases. Training was performed using the Limited-memory Broyden–Fletcher–Goldfarb–Shanno (L-BFGS) solver without explicit L2 regularization ($\lambda = 0$). Convergence was defined by standard gradient, loss, and step tolerances of $1.0 \times 10^{-6}$. Training was capped at 1000 iterations to prevent overfitting and memorization while maximizing sequence-level generalization, assessed using a sequence-level leave-one-out cross-validation (LOOCV) strategy. (Supplementary Figure 37). The final model was trained on the complete dataset of 64 experimentally validated sequences and evaluated against an external validation set of 11 unseen peptide sequences within the same chemical space.

#### 2.20 Data Analysis and Statistics

Statistical analyses, non-linear regression, and 3D plotting were performed using GraphPad Prism (version 10) and OriginPro 2024b. Python (version 3.12.10) was utilized for additional data processing and figure generation, employing the NumPy, pandas, and stats models libraries. Custom scripts and macros used for analysis are available at https://github.com/Wei0O/lipidation-site. High-throughput analysis of confocal images (spanning >70 protein conditions) was performed in Fiji using a custom macro. The automated pipeline for partition coefficient analysis included background subtraction, droplet segmentation via automatic thresholding (Otsu), and binary mask generation. Results were exported as CSV files, with overlay images generated for quality control.

#### 3. Supplementary Tables

**Supplementary Table 1. Oligonucleotides and PCR Annealing Temperatures Used to Construct the Lipidation-Site Library.**

| **Plasmid** pETDuet_ NMT_ [$\underline{\mathbf{A_2}}A_3A_4$]-$V_{30}$ | **Primers (5′–3′)** | | **$T_a$ (°C)** | **Bridge oligo sequence**[1] |
|---|---|---|---|---|
| | Forward | Reverse | | |
| $A_2$ = Gly | TCTCGTGGTTCTTCCGGTTC | CCCTCCCATATGTATATCTC | 59 | *AGAAGGAGATATACATATGGGA* *GGG*KBGKBG *TCTCGTGGTTCTTCCGGTTCCAGTG* |
| $A_2$ = Ala | TCTCGTGGTTCTTCCGGTTC | TGCTCCCATATGTATATCTC | 58 | *AGAAGGAGATATACATATGGGA* *GCA*KBGKBG *TCTCGTGGTTCTTCCGGTTCCAGTG* |
| $A_2$ = Leu | TCTCGTGGTTCTTCCGGTTC | CAATCCCATATGTATATCTC | 53 | *AGAAGGAGATATACATATGGGA* *TTG*KBGKBG *TCTCGTGGTTCTTCCGGTTCCAGTG* |
| $A_2$= Ser | TCGCGTGGTAGTTCGGGTAG | ACTTCCCATATGTATATCTC | 52 | *AGAAGGAGATATACATATGGG* *AAGT*KBGKBG *TCGCGTGGTAGTTCGGGTAGCTCCG* |

[1] Bridge ssDNA sequences are segmented into three lines for readability. Italicized regions indicate overlap with the vector backbone. Underlined sequences denote the variable lipidation site, encoding either degenerate 'KBG' codons or a deterministic sequence.

**Supplementary Table 2. Sequences of Lipidation-Site and Miscibility Library Constructs.**

| Construct | lipid | Sequence |
|---|---|---|
| m-[$A_2A_3A_4$]-$V_{30}$ | C14:0 | G$\mathbf{A_2A_3A_4}$SRGSSGSS(GVGVP)$_{30}$GWP |
| $V_{30}$ | n.a. | (GVGVP)$_{30}$ GWP |
| m-$V_{30}$ | C14:0 | |
| [LYA]-(V/A/K)$_{80}$ | n.a. | GLYASKLFSNL[(GVGVP GVGVP GAGVP GVGVP GVGVP)$_2$ GGK]$_8$GY |
| m-[LYA]-(V/A/K)$_{80}$ | C14:0 | |
| [LYA]-$V_{40}$ | n.a. | GLYASKLFSNL(GVGVP)$_{40}$ GY |
| m-[LYA]-$V_{40}$ | C14:0 | |
| [LYA]-$V_{80}$ | n.a. | GLYASKLFSNL(GVGVP)$_{80}$ GY |
| m-[LYA]-$V_{80}$ | C14:0 | |

$A_2$,$A_3$, $A_4$ $\in$ {G,A,S,L}; The Z-residue referenced in the nomenclature is underlined.

**Supplementary Table 3. Theoretical and Observed Molecular Weights of the Constructs.**

| Construct | Observed m/z | Theoretical Mw (Da) | Δ (%) |
|---|---|---|---|
| m-[AAA]-$V_{30}$ | 13830.62 | 13829.29 | 0.010 |
| m-[AAG]-$V_{30}$ | 13815.28 | 13815.26 | 0 |
| m-[AAL]-$V_{30}$ | 13869.33 | 13871.37 | −0.015 |
| m-[AAS]-$V_{30}$ | 13846.34 | 13845.29 | 0.008 |
| m-[AGA]-$V_{30}$ | 13808.81 | 13815.26 | −0.047 |
| m-[AGG]-$V_{30}$ | 13807.11 | 13801.23 | 0.043 |
| m-[AGL]-$V_{30}$ | 13855.73 | 13857.34 | −0.012 |
| m-[AGS]-$V_{30}$ | 13840.66 | 13831.26 | 0.068 |
| m-[ALA]-$V_{30}$ | 13866.98 | 13871.37 | −0.032 |
| m-[ALG]-$V_{30}$ | 13863.72 | 13857.34 | 0.046 |
| m-[ALL]-$V_{30}$ | 13910.15 | 13913.45 | −0.024 |
| m-[ALS]-$V_{30}$ | 13888.47 | 13887.37 | 0.008 |
| m-[ASA]-$V_{30}$ | 13844.40 | 13845.29 | −0.006 |
| m-[ASG]-$V_{30}$ | 13835.81 | 13831.26 | 0.033 |
| m-[ASL]-$V_{30}$ | 13884.72 | 13887.37 | −0.019 |
| m-[ASS]-$V_{30}$ | 13862.81 | 13861.29 | 0.011 |
| m-[GAA]-$V_{30}$ | 13819.57 | 13815.26 | 0.031 |
| m-[GAG]-$V_{30}$ | 13803.39 | 13801.23 | 0.016 |
| m-[GAL]-$V_{30}$ | 13853.23 | 13857.34 | −0.030 |
| m-[GAS]-$V_{30}$ | 13829.78 | 13831.26 | −0.011 |
| m-[GGA]-$V_{30}$ | 13804.19 | 13801.23 | 0.021 |
| m-[GGG]-$V_{30}$ | 13775.41 | 13787.21 | −0.086 |
| m-[GGL]-$V_{30}$ | 13845.33 | 13843.32 | 0.015 |
| m-[GGS]-$V_{30}$ | 13829.78 | 13817.23 | 0.091 |
| m-[GLA]-$V_{30}$ | 13860.73 | 13857.34 | 0.024 |
| m-[GLG]-$V_{30}$ | 13846.35 | 13843.32 | 0.022 |
| m-[GLL]-$V_{30}$ | 13895.28 | 13899.42 | −0.030 |
| m-[GLS]-$V_{30}$ | 13873.53 | 13873.34 | 0.001 |
| m-[GSA]-$V_{30}$ | 13830.01 | 13831.26 | −0.009 |
| m-[GSG]-$V_{30}$ | 13817.86 | 13817.23 | 0.005 |
| m-[GSL]-$V_{30}$ | 13871.24 | 13873.34 | −0.015 |
| m-[GSS]-$V_{30}$ | 13851.19 | 13847.26 | 0.028 |
| m-[LAA]-$V_{30}$ | 13864.20 | 13871.37 | −0.052 |
| m-[LAG]-$V_{30}$ | 13860.92 | 13857.34 | 0.026 |
| m-[LAL]-$V_{30}$ | 13915.09 | 13913.45 | 0.012 |
| m-[LAS]-$V_{30}$ | 13888.82 | 13887.37 | 0.010 |
| m-[LGA]-$V_{30}$ | 13848.71 | 13857.34 | −0.062 |
| m-[LGG]-$V_{30}$ | 13846.35 | 13843.32 | 0.022 |
| m-[LGL]-$V_{30}$ | 13898.17 | 13899.42 | −0.009 |
| m-[LGS]-$V_{30}$ | 13872.58 | 13873.34 | −0.005 |
| m-[LLA]-$V_{30}$ | 13912.17 | 13913.45 | −0.009 |

| | | | |
|---|---|---|---|
| m-[LLG]-$V_{30}$ | 13904.58 | 13899.42 | 0.037 |
| m-[LLL]-$V_{30}$ | 13970.35 | 13955.53 | 0.106 |
| m-[LLS]-$V_{30}$ | 13932.61 | 13929.45 | 0.023 |
| m-[LSA]-$V_{30}$ | 13883.82 | 13887.37 | −0.026 |
| m-[LSG]-$V_{30}$ | 13886.42 | 13873.34 | 0.094 |
| m-[LSL]-$V_{30}$ | 13921.43 | 13929.45 | −0.058 |
| m-[LSS]-$V_{30}$ | 13900.83 | 13903.37 | −0.018 |
| m-[SAA]-$V_{30}$ | 13848.50 | 13845.29 | 0.023 |
| m-[SAG]-$V_{30}$ | 13834.37 | 13831.26 | 0.022 |
| m-[SAL]-$V_{30}$ | 13891.54 | 13887.37 | 0.030 |
| m-[SAS]-$V_{30}$ | 13873.31 | 13861.29 | 0.087 |
| m-[SGA]-$V_{30}$ | 13832.57 | 13831.26 | 0.009 |
| m-[SGG]-$V_{30}$ | 13815.49 | 13817.23 | −0.013 |
| m-[SGL]-$V_{30}$ | 13883.92 | 13873.34 | 0.076 |
| m-[SGS]-$V_{30}$ | 13871.55 | 13847.26 | 0.175 |
| m-[SLA]-$V_{30}$ | 13873.31 | 13887.37 | −0.101 |
| m-[SLG]-$V_{30}$ | 13876.26 | 13873.34 | 0.021 |
| m-[SLL]-$V_{30}$ | 13949.93 | 13929.45 | 0.147 |
| m-[SLS]-$V_{30}$ | 13898.74 | 13903.37 | −0.033 |
| m-[SSA]-$V_{30}$ | 13863.97 | 13861.29 | 0.019 |
| m-[SSG]-$V_{30}$ | 13851.80 | 13847.26 | 0.033 |
| m-[SSL]-$V_{30}$ | 13905.17 | 13903.37 | 0.013 |
| m-[SSS]-$V_{30}$ | 13863.93 | 13877.29 | −0.096 |
| $V_{30}$ | 12651.00 | 12642.97 | 0.063 |
| m-$V_{30}$ | 12849.20 | 12853.32 | −0.032 |
| [LYA]-$(V/A/K)_{80}$ | 35673.04 | 35680.86 | −0.022 |
| m-[LYA]-$(V/A/K)_{80}$ | 35961.78 | 35891.16 | 0.197 |
| [LYA]-$V_{40}$ | 17806.02 | 17812.07 | −0.034 |
| m-[LYA]-$V_{40}$ | 18013.05 | 18022.37 | −0.050 |
| [LYA]-$V_{80}$ | 34198.00 | 34191.49 | 0.019 |
| m-[LYA]-$V_{80}$ | 34407.95 | 34401.79 | 0.018 |
| [SAA]-$V_{30}$ ($^{15}N$) | 13813.38 | 13795.07 | 0.133 |
| Linker-$V_{30}$ ($^{15}N$) | 13559.16 | 13562.97 | −0.028 |
| m-[SAA]-$V_{30}$ ($^{15}N$) | 14007.08 | 14005.42 | 0.012 |
| m-[SAL]-$V_{30}$ ($^{15}N$) | 14051.81 | 14055.04 | −0.023 |
| $V_{30}$ ($^{15}N$) | 12787.20 | 12788.68 | −0.012 |
| [LLL]-$V_{30}$ | 13750.80 | 13745.18 | 0.041 |
| [SAA]-$V_{30}$ | 13813.38 | 13634.94 | 1.309 |
| [SAS]-$V_{30}$ | 13655.48 | 13650.94 | 0.033 |
| [SLL]-$V_{30}$ | 13729.24 | 13719.10 | 0.074 |
| [SSS]-$V_{30}$ | 13657.44 | 13666.94 | −0.069 |

**Supplementary Table 4. Theoretical and Observed Molecular Weights of Trypsinized N-Terminal Fragments.**

| Construct | Observed m/z | Theoretical Mw (Da) | Δ (%) |
|---|---|---|---|
| m-[AAA]-$V_{30}$ | 742.473 | 742.633 | −0.022 |
| m-[AAG]-$V_{30}$ | 728.493 | 728.618 | −0.017 |
| m-[AAL]-$V_{30}$ | 784.551 | 784.680 | −0.016 |
| m-[AAS]-$V_{30}$ | 758.543 | 758.628 | −0.011 |
| m-[AGA]-$V_{30}$ | 728.518 | 728.618 | −0.014 |
| m-[AGG]-$V_{30}$ | 714.476 | 714.602 | −0.018 |
| m-[AGL]-$V_{30}$ | 770.632 | 770.665 | −0.004 |
| m-[AGS]-$V_{30}$ | 744.438 | 744.613 | −0.024 |
| m-[ALA]-$V_{30}$ | 784.576 | 784.680 | −0.013 |
| m-[ALG]-$V_{30}$ | 770.556 | 770.665 | −0.014 |
| m-[ALL]-$V_{30}$ | 826.612 | 826.727 | −0.014 |
| m-[ALS]-$V_{30}$ | 800.449 | 800.675 | −0.028 |
| m-[ASA]-$V_{30}$ | 758.543 | 758.628 | −0.011 |
| m-[ASG]-$V_{30}$ | 744.594 | 744.613 | −0.003 |
| m-[ASL]-$V_{30}$ | 800.614 | 800.675 | −0.008 |
| m-[ASS]-$V_{30}$ | 774.480 | 774.623 | −0.018 |
| m-[GAA]-$V_{30}$ | 728.581 | 728.618 | −0.005 |
| m-[GAG]-$V_{30}$ | 714.417 | 714.602 | −0.026 |
| m-[GAL]-$V_{30}$ | 770.517 | 770.665 | −0.019 |
| m-[GAS]-$V_{30}$ | 744.465 | 744.613 | −0.020 |
| m-[GGA]-$V_{30}$ | 714.417 | 714.602 | −0.026 |
| m-[GGG]-$V_{30}$ | 700.570 | 700.586 | −0.002 |
| m-[GGL]-$V_{30}$ | 756.474 | 756.649 | −0.023 |
| m-[GGS]-$V_{30}$ | 730.401 | 730.597 | −0.027 |
| m-[GLA]-$V_{30}$ | 770.467 | 770.665 | −0.026 |
| m-[GLG]-$V_{30}$ | 756.525 | 756.649 | −0.016 |
| m-[GLL]-$V_{30}$ | 812.514 | 812.712 | −0.024 |
| m-[GLS]-$V_{30}$ | 786.579 | 786.660 | −0.010 |
| m-[GSA]-$V_{30}$ | 744.449 | 744.613 | −0.022 |
| m-[GSG]-$V_{30}$ | 730.450 | 730.597 | −0.020 |
| m-[GSL]-$V_{30}$ | 786.426 | 786.660 | −0.030 |
| m-[GSS]-$V_{30}$ | 760.538 | 760.607 | −0.009 |
| m-[LAA]-$V_{30}$ | 784.485 | 784.680 | −0.025 |
| m-[LAG]-$V_{30}$ | 770.581 | 770.665 | −0.011 |
| m-[LAL]-$V_{30}$ | 826.534 | 826.727 | −0.023 |
| m-[LAS]-$V_{30}$ | 800.460 | 800.675 | −0.027 |
| m-[LGA]-$V_{30}$ | 770.535 | 770.665 | −0.017 |
| m-[LGG]-$V_{30}$ | 756.575 | 756.649 | −0.010 |
| m-[LGL]-$V_{30}$ | 812.586 | 812.712 | −0.016 |
| m-[LGS]-$V_{30}$ | 786.517 | 786.660 | −0.018 |

| | | | |
|---|---|---|---|
| m-[LLA]-$V_{30}$ | 826.508 | 826.727 | −0.026 |
| m-[LLG]-$V_{30}$ | 812.514 | 812.712 | −0.024 |
| m-[LLL]-$V_{30}$ | 868.486 | 868.774 | −0.033 |
| m-[LLS]-$V_{30}$ | 842.640 | 842.722 | −0.010 |
| m-[LSA]-$V_{30}$ | 800.419 | 800.675 | −0.032 |
| m-[LSG]-$V_{30}$ | 786.451 | 786.660 | −0.027 |
| m-[LSL]-$V_{30}$ | 842.477 | 842.722 | −0.029 |
| m-[LSS]-$V_{30}$ | 816.491 | 816.670 | −0.022 |
| m-[SAA]-$V_{30}$ | 758.543 | 758.628 | −0.011 |
| m-[SAG]-$V_{30}$ | 744.465 | 744.613 | −0.020 |
| m-[SAL]-$V_{30}$ | 800.449 | 800.675 | −0.028 |
| m-[SAS]-$V_{30}$ | 774.404 | 774.623 | −0.028 |
| m-[SGA]-$V_{30}$ | 744.435 | 744.613 | −0.024 |
| m-[SGG]-$V_{30}$ | 730.412 | 730.597 | −0.025 |
| m-[SGL]-$V_{30}$ | 786.440 | 786.660 | −0.028 |
| m-[SGS]-$V_{30}$ | 760.450 | 760.607 | −0.021 |
| m-[SLA]-$V_{30}$ | 800.614 | 800.675 | −0.008 |
| m-[SLG]-$V_{30}$ | 786.465 | 786.66 | −0.025 |
| m-[SLL]-$V_{30}$ | 842.481 | 842.722 | −0.029 |
| m-[SLS]-$V_{30}$ | 816.456 | 816.670 | −0.026 |
| m-[SSA]-$V_{30}$ | 774.568 | 774.623 | −0.007 |
| m-[SSG]-$V_{30}$ | 760.450 | 760.607 | −0.021 |
| m-[SSL]-$V_{30}$ | 816.566 | 816.670 | −0.013 |
| m-[SSS]-$V_{30}$ | 790.430 | 790.618 | −0.024 |
| [LYA]- $(V/A/K)_{80}$ | 638.340 | 637.730 | 0.096 |
| m-[LYA]- $(V/A/K)_{80}$ | 848.593 | 848.030 | 0.066 |
| [LYA]-$V_{40}$ | 638.294 | 637.730 | 0.088 |
| m-[LYA]-$V_{40}$ | 848.699 | 848.030 | 0.079 |
| [LYA]-$V_{80}$ | 638.363 | 637.730 | 0.099 |
| m-[LYA]-$V_{80}$ | 848.725 | 848.030 | 0.082 |

#### 4. Supplementary Figures

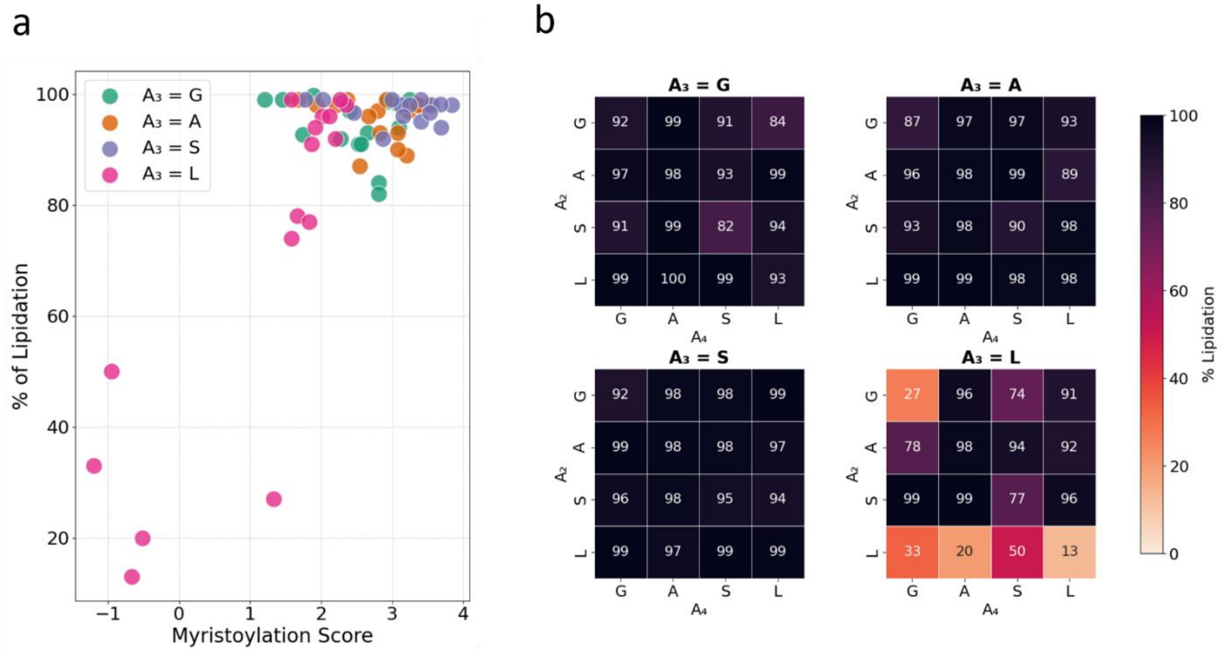

**Supplementary Figure 1. Assessment of lipidation efficiency across the lipidation-site library. a,** Scatterplot of experimentally measured *in vivo* myristoylation efficiency versus predicted myristoylation score,[30] colored by $A_3$ residue identity. **b,** Heatmaps of mean % lipidation for all variants, faceted by $A_3$ residue identity, demonstrating the higher sensitivity of NMT to bulky amino acids (e.g., Leu). These data confirm that the library design effectively samples the permissive landscape of NMT substrate specificity. Irrespective of the lipidation efficiency during expression, all constructs were purified to >95% homogeneity (Supplementary Figure. 2).

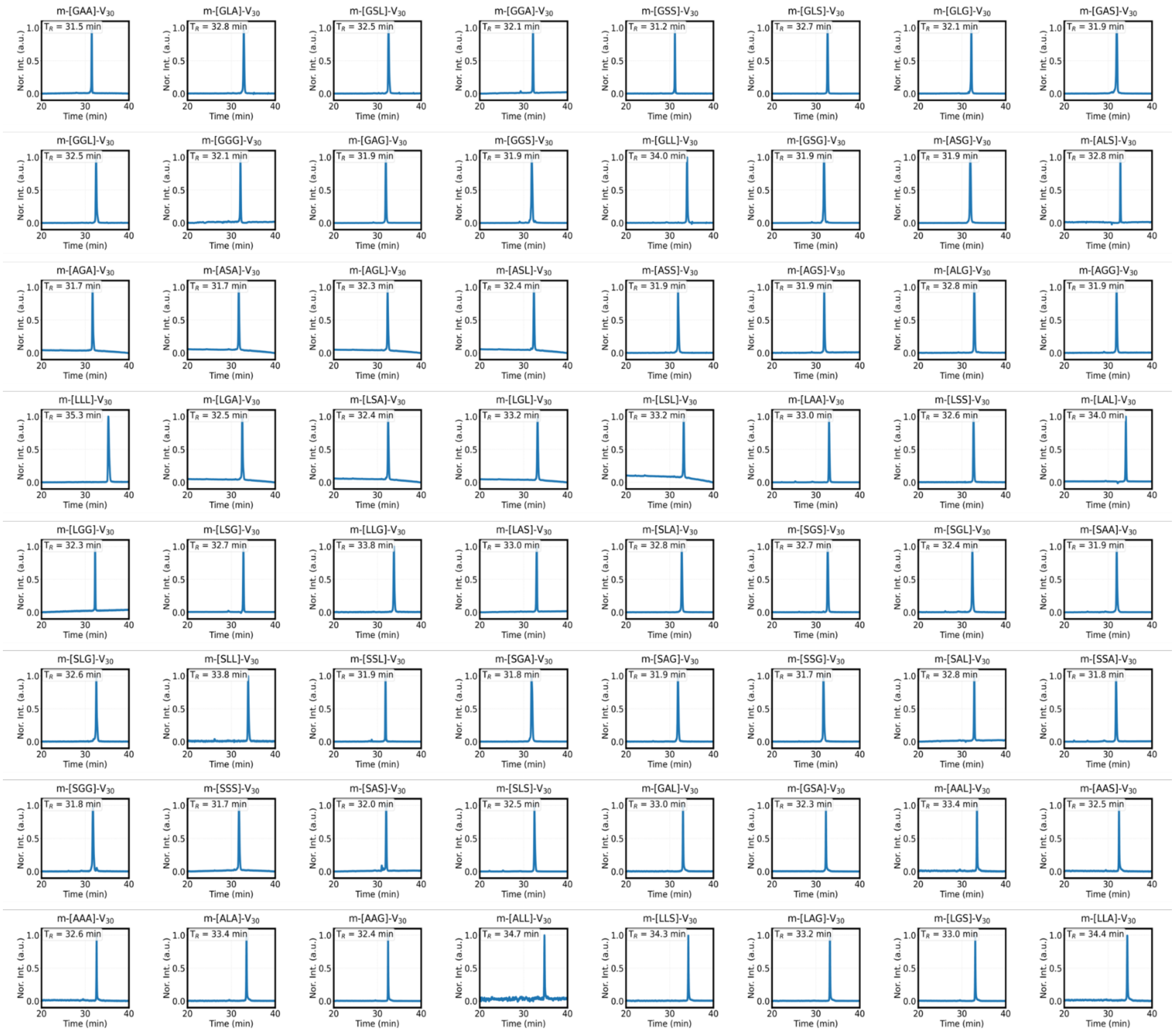

**Supplementary Figure 2. RP-HPLC characterization of the lipidation-site library.** Chromatograms of the 64 lipidated variants (m-[$A_2A_3A_4$]-$V_{30}$. Traces ($A_{210}$) were baseline-subtracted and normalized to the maximum signal within the plotted window. Inset values indicate retention time ($t_R$), reflecting the bulk hydrophobicity of the lipidation site triad.

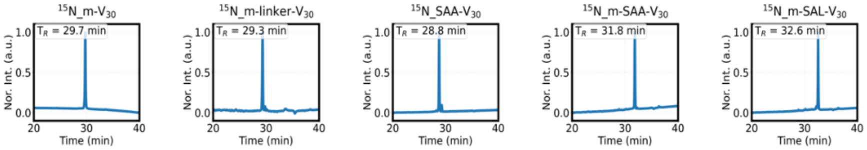

**Supplementary Figure 3. RP-HPLC characterization of $^{15}$N-labeled constructs.** Chromatograms of five constructs isotopically labeled with $^{15}$N. Traces ($A_{210}$) were baseline-subtracted and normalized to the maximum signal within the plotted window. Inset values indicate retention time ($t_R$).

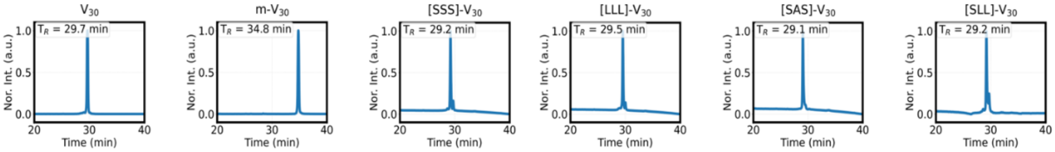

**Supplementary Figure 4. RP-HPLC characterization of control constructs.** Chromatograms of non-lipidated $V_{30}$, lipidated m-$V_{30}$ lacking the lipidation site triad, and non-myristoylated variants used for fuel-driven LLPS experiments. Traces are baseline-subtracted and normalized to the maximum signal within the plotted window. Inset values indicate retention time ($t_R$).

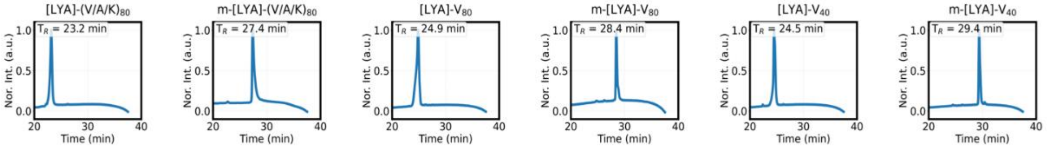

**Supplementary Figure 5. RP-HPLC characterization of miscibility library proteins.** Traces are baseline-subtracted and normalized to the maximum signal within the plotted window. Inset values indicate retention time ($t_R$).

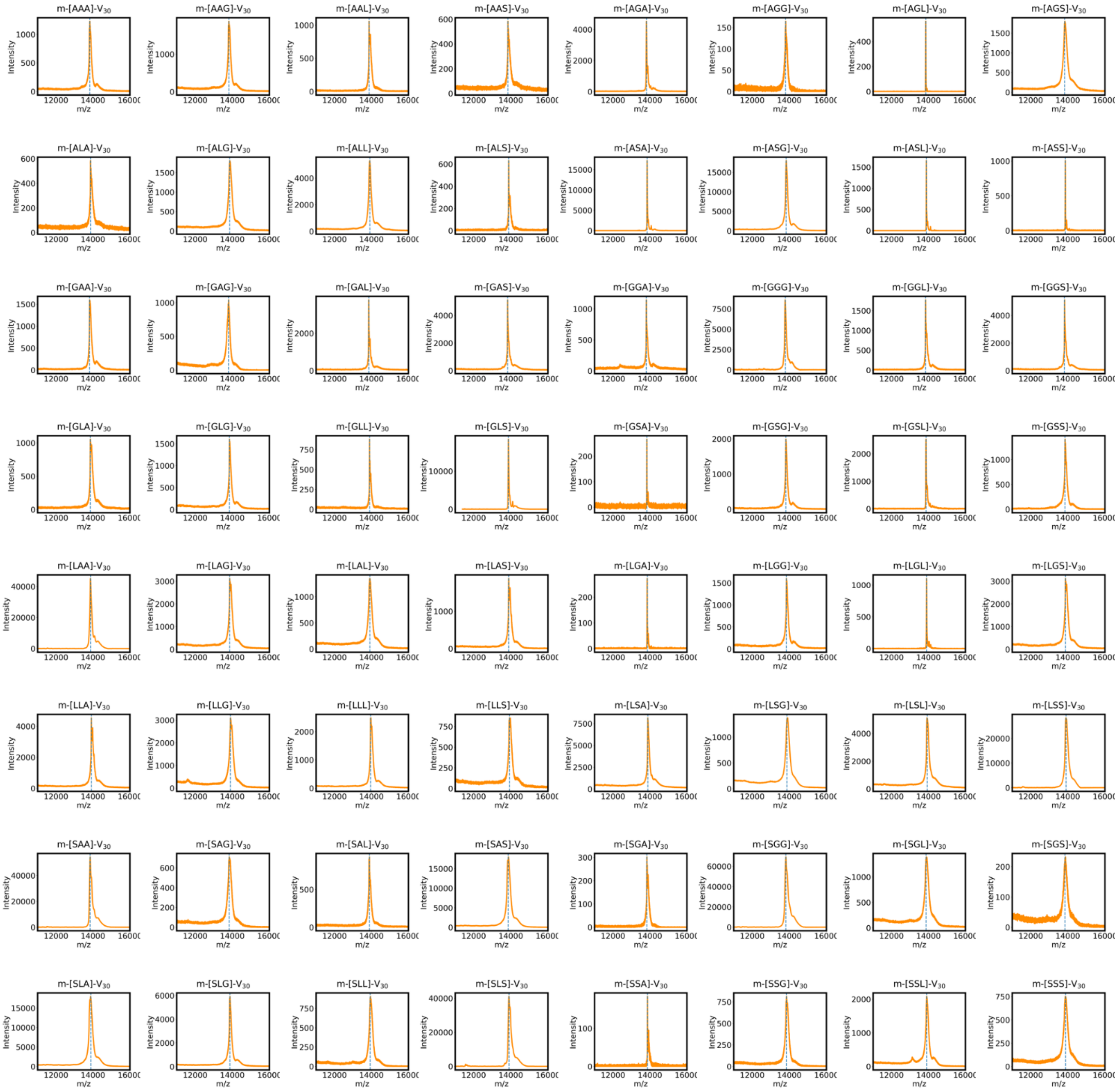

**Supplementary Figure 6. MALDI-TOF characterization of the lipidation-site library.** Mass spectra of the 64 lipidated variants (m-[$A_2A_3A_4$]-$V_{30}$). Vertical lines indicate theoretical m/z values. Detailed observed and theoretical mass values are listed in Supplementary Table 3.

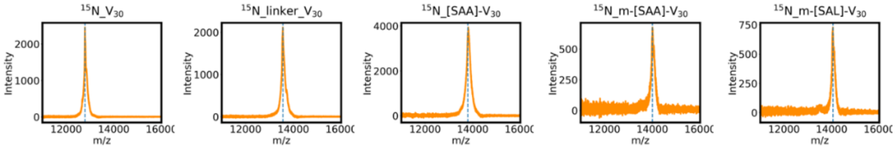

**Supplementary Figure 7. MALDI-TOF characterization of $^{15}$N-labeled proteins.** Vertical lines indicate theoretical m/z values. Detailed observed and theoretical mass values are listed in Supplementary Table 3.

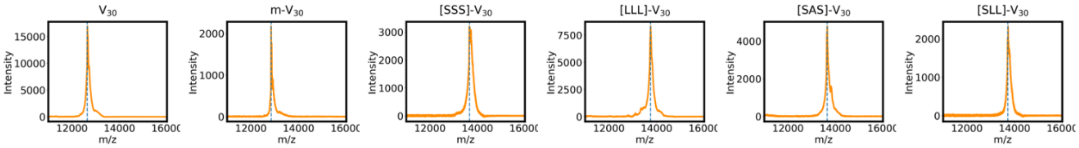

**Supplementary Figure 8. MALDI-TOF characterization of control proteins.** Vertical dashed lines indicate theoretical molecular weights. Detailed observed and theoretical mass values are listed in Supplementary Table 3.

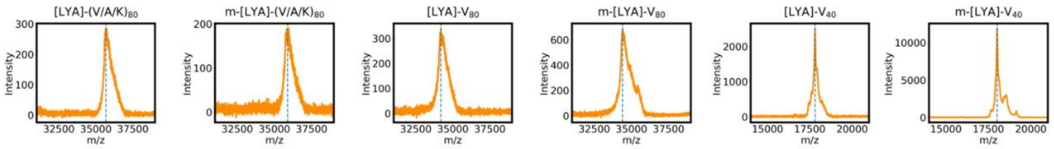

**Supplementary Figure 9. MALDI-TOF characterization of miscibility library proteins.** Vertical dashed lines indicate theoretical molecular weights. Detailed observed and theoretical mass values are listed in Supplementary Table 3.

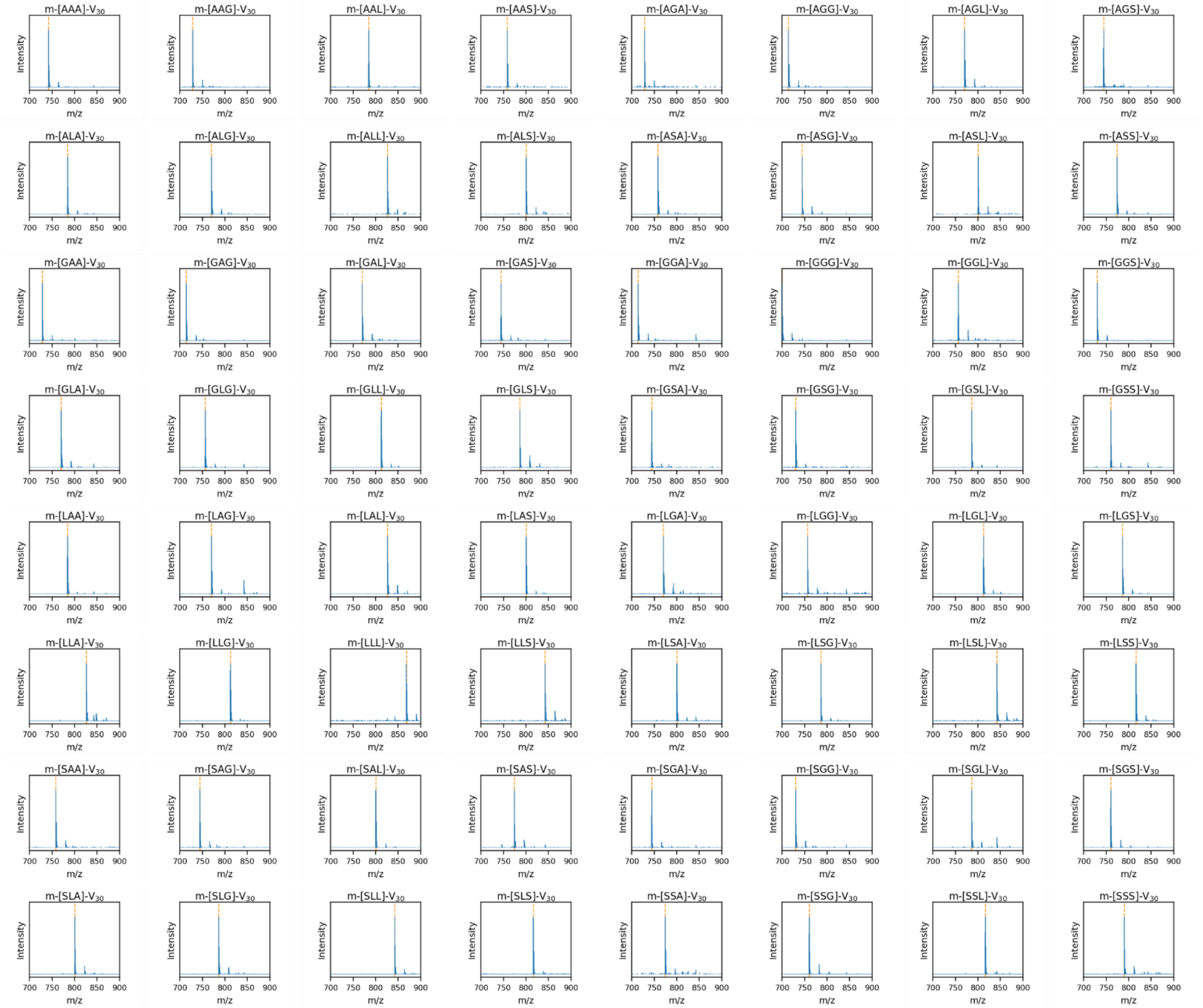

**Supplementary Figure 10. MALDI-TOF spectra of N-terminal fragments of the lipidation-site library.** Spectra of trypsin-digested N-terminal peptides corresponding to the myristoylated hexapeptide (C14:0-$GA_2A_3A_4SR$). Vertical dashed lines indicate theoretical m/z values. Detailed mass values are listed in Supplementary Table 4.

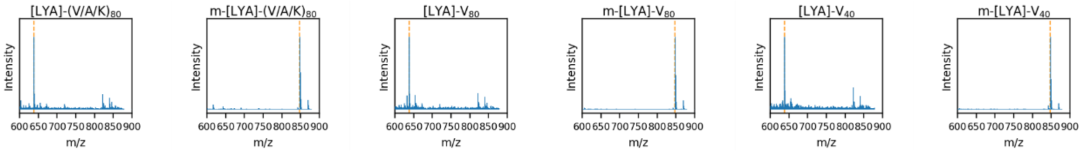

**Supplementary Figure 11. MALDI-TOF spectra of N-terminal fragments of the miscibility library.** Spectra of trypsin-digested N-terminal peptides corresponding to the unmodified or myristoylated hexapeptide (GLYASK). Vertical dashed lines indicate theoretical m/z values. Detailed mass values are listed in Supplementary Table 4.

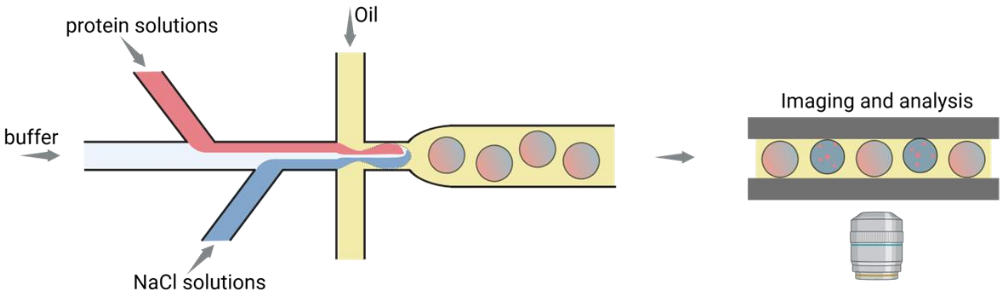

**Supplementary Figure 12. Schematic of the PhaseScan microfluidic platform for high-throughput phase diagram mapping.** Aqueous inputs (protein solution, buffer, and NaCl solution) are mixed on chip and segmented into water-in-oil microdroplets. Flow-rate ratios are modulated to generate droplets with combinatorial protein and salt concentrations. Droplets are equilibrated at desired temperature and imaged to determine the phase separation boundary.

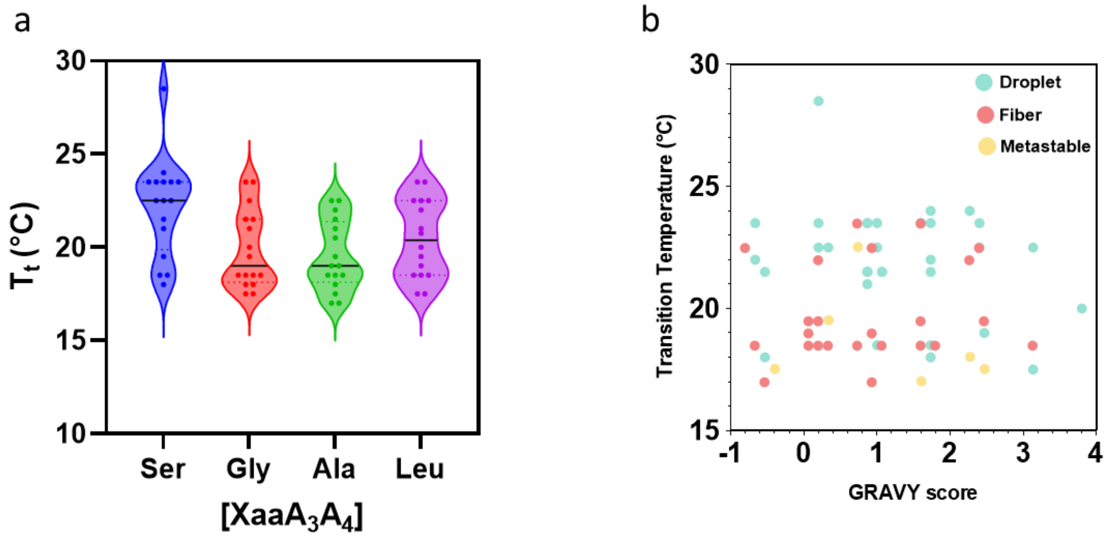

**Supplementary Figure 13. Relationship between transition temperature, sequence identity, and condensate material properties. a,** Violin plots of $T_t$ of lipidation-site library grouped by amino acid identity at position $A_2$. Solid black lines indicate median values. Serine at position $A_2$ elevated $T_t$, suggesting its polarity may counteract and attenuate the hydrophobic contribution of the myristoyl lipid to cohesive forces. **b,** Plot of $T_t$ vs. calculated hydropathy (GRAVY score). Constructs are colored by condensed-phase properties: liquid droplet (teal), solid fiber (red), and metastable gels (yellow). Fiber-forming variants mostly cluster at lower $T_t$ values despite occupying a similar range of GRAVY scores as droplet-forming variants, suggesting a difference in pre-condensation oligomers.

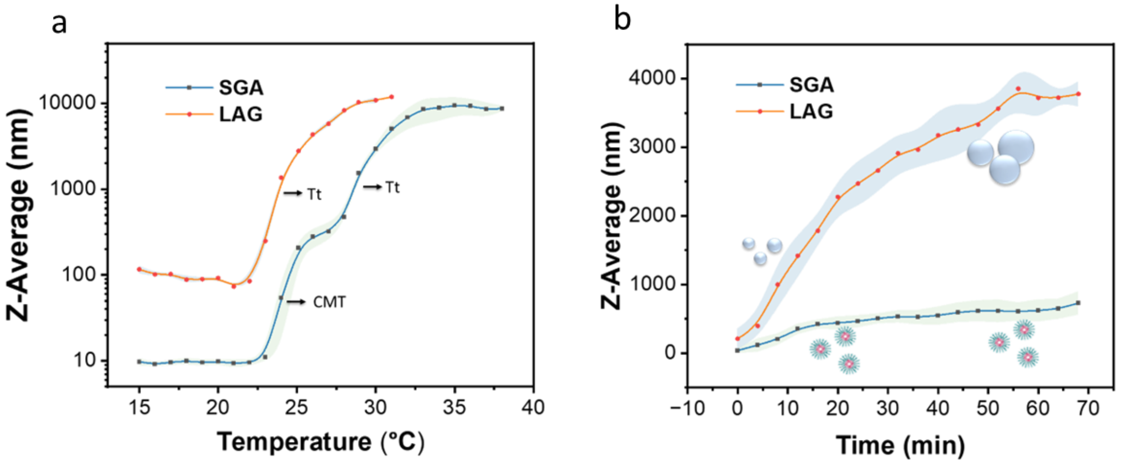

**Supplementary Figure 14. DLS analysis revealing distinct micellar and phase-separation pathways. a,** Temperature-dependent hydrodynamic radius (Z-Average) for m-[LAG]-$V_{30}$ and m-[SGA]-$V_{30}$ variants. LAG displays a single sharp transition to large aggregates, whereas SGA exhibits a two-step transition: an initial increase at the critical micelle temperature (CMT, 24 °C) followed by a macroscopic transition ($T_t$, 29 °C). **b,** Isothermal kinetic stability monitored at 26 °C. LAG undergoes rapid size growth, whereas SGA maintains a stable hydrodynamic radius, consistent with the formation of kinetically stable micelles. Data are mean ± s.d. (n = 3).

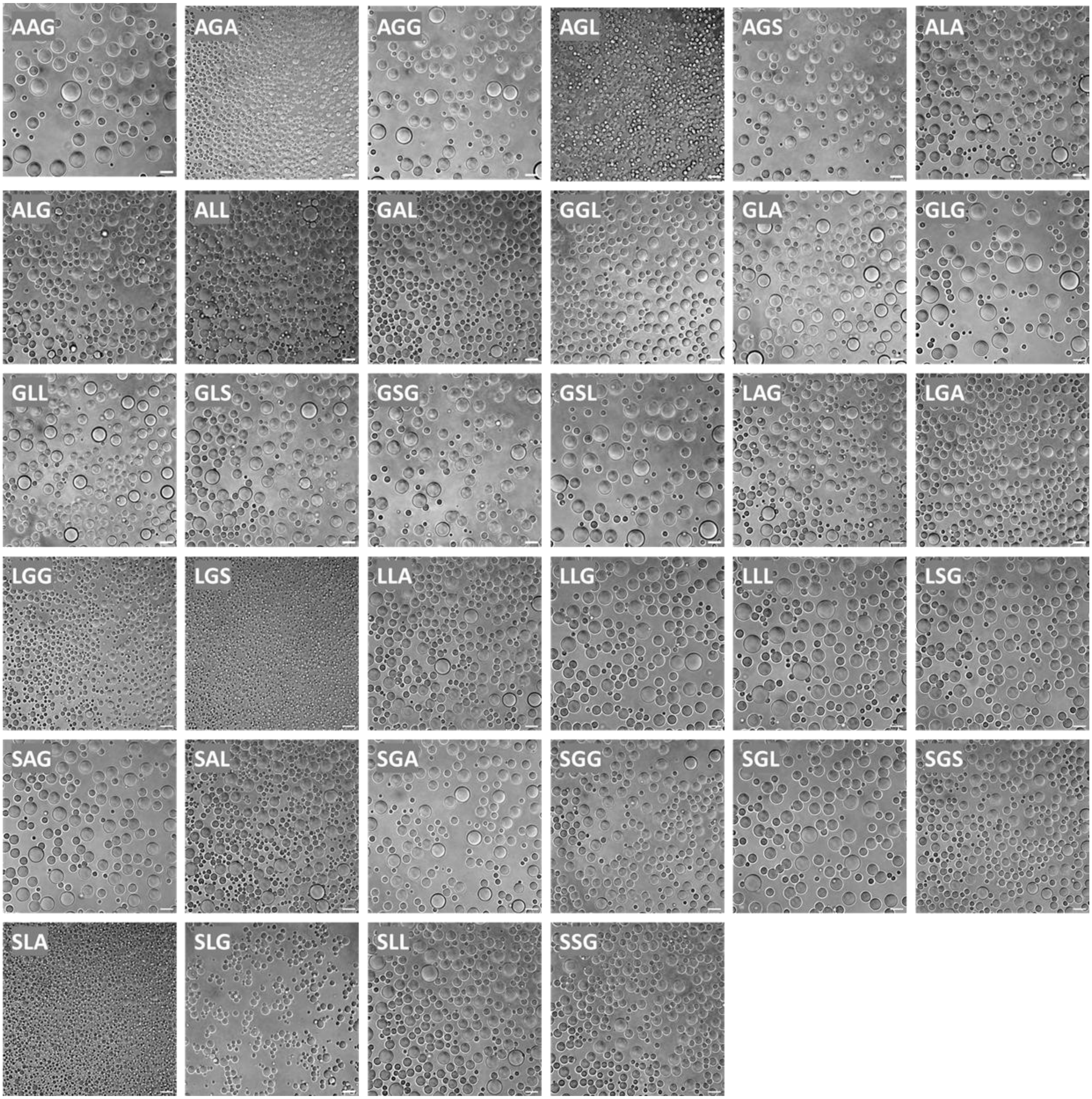

**Supplementary Figure 15. DIC micrographs of liquid condensates.** Images display the 34 library variants classified as liquid condensates, exhibiting the spherical droplets characteristic of dynamic liquid–liquid phase separation. Constructs are labeled by their triad motif ($A_2A_3A_4$). Scale bars, 10 μm.

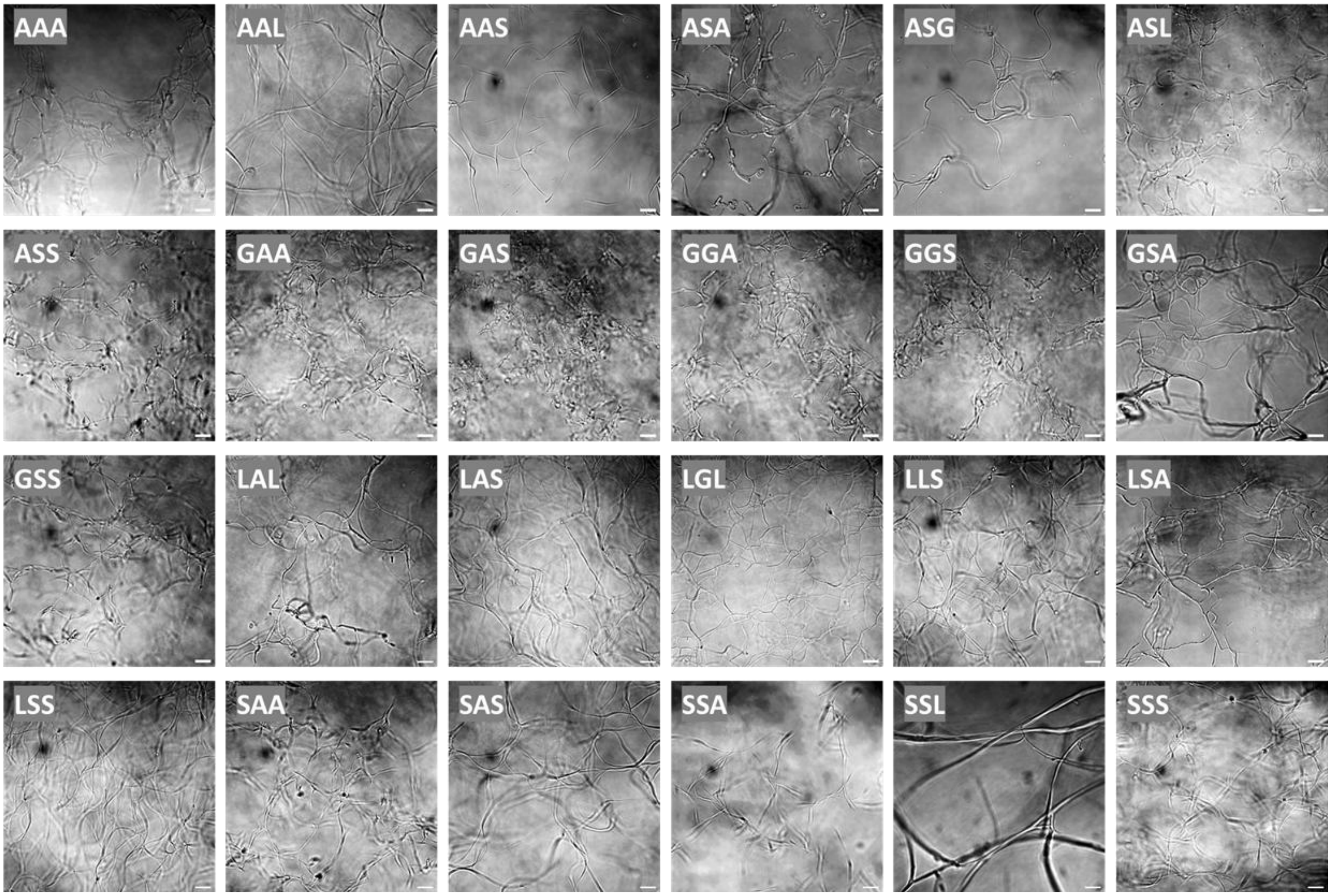

**Supplementary Figure 16. DIC micrographs of solid fibrillar aggregates.** Images display the 24 library variants classified as fiber-formers. These variants exhibit extended, anisotropic fibrillar networks characteristic of liquid–solid phase transition. Constructs are labeled by their triad motif ($A_2A_3A_4$). Scale bars, 10 µm.

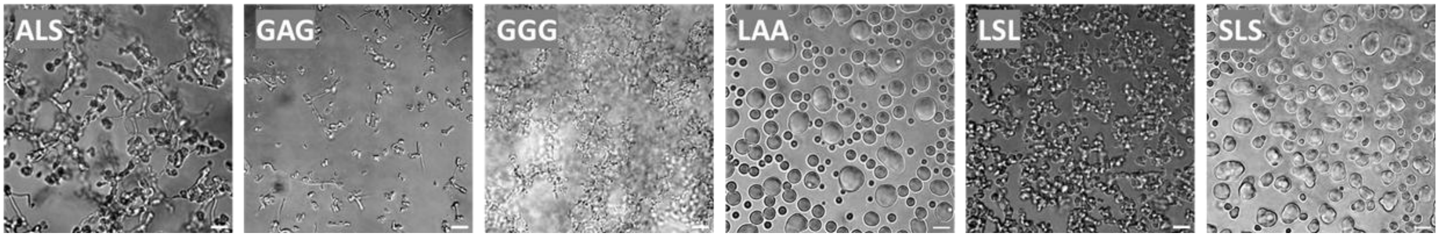

**Supplementary Figure 17. DIC micrographs of metastable assemblies.** Images display the six library variants classified as metastable. These variants exhibit heterogeneous morphologies characterized by irregular metastable gels or amorphous precipitates. Constructs are labeled by their triad motif ($A_2A_3A_4$). Scale bars, 10 μm.

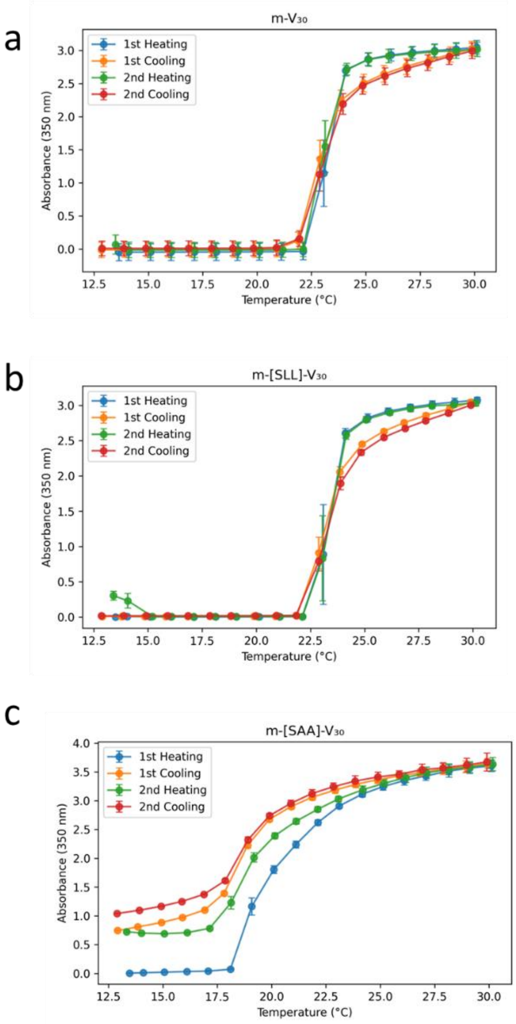

**Supplementary Figure 18. Reversibility of phase transitions for selected constructs monitored over two consecutive heating/cooling cycles. a, b,** Control m-$V_{30}$ (**a**) and droplet-forming m-[SLL]-$V_{30}$ (**b**) exhibit fully reversible transitions with minimal hysteresis, consistent with liquid-liquid phase-separation. **c,** Fiber-forming m-[SAA]-V30 displays significant thermal hysteresis and irreversibility, consistent with liquid-solid phase separation. Data are mean ± s.d. (n = 3).

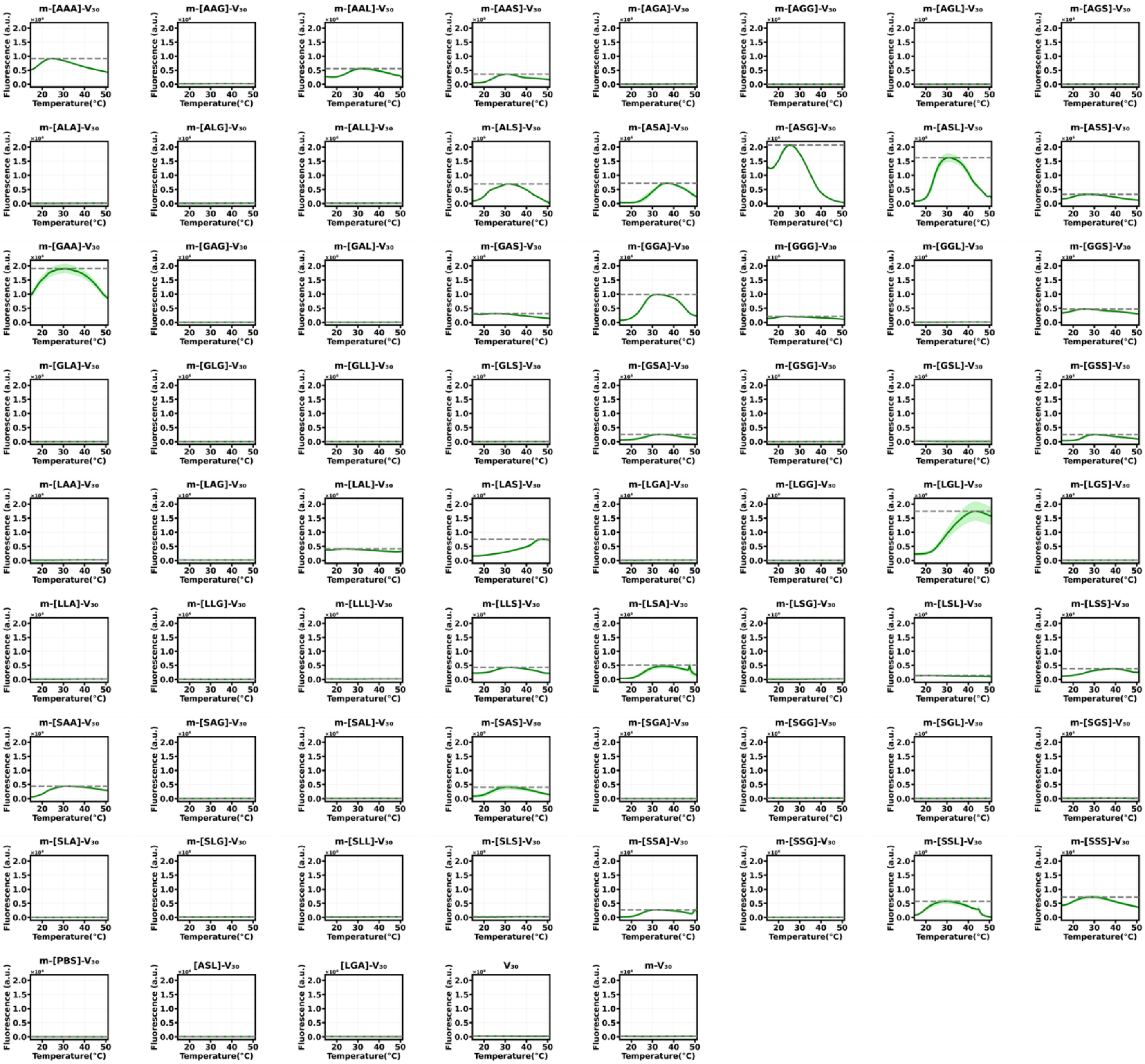

**Supplementary Figure 19. Spectroscopic confirmation of cross-β sheet order.** Thioflavin T (ThT) fluorescence monitored as a function of temperature for the lipidation-site library and control samples. High fluorescence intensity is observed in constructs forming solid fibers, with minimal fluorescence observed in liquid condensates, confirming the presence of ordered, amyloid-like architecture in the solid phase. Data are presented as mean (solid lines) ± s.d. (shaded regions; n = 3). Horizontal dashed lines indicate maximum ThT intensity.

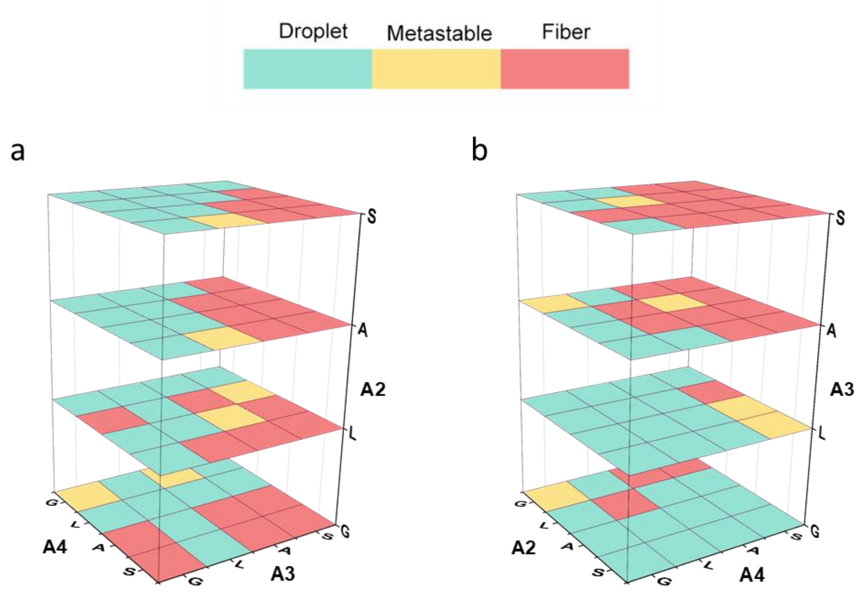

**Supplementary Figure 20. Deconvolution of sequence determinants of condensed-phase material states. a–b,** 3D stacked heatmaps of material states for the 64-member library sliced by position $A_2$ (**a**), $A_3$ (**b**) on Z-axis. Position $A_2$ minimally impacts material properties, whereas positions $A_3$ and $A_4$ (Fig. 3e) exert the strongest influence on condensate properties, with Gly/Leu at these positions strongly favoring liquid droplets and Ala/Ser favoring solid fibers.

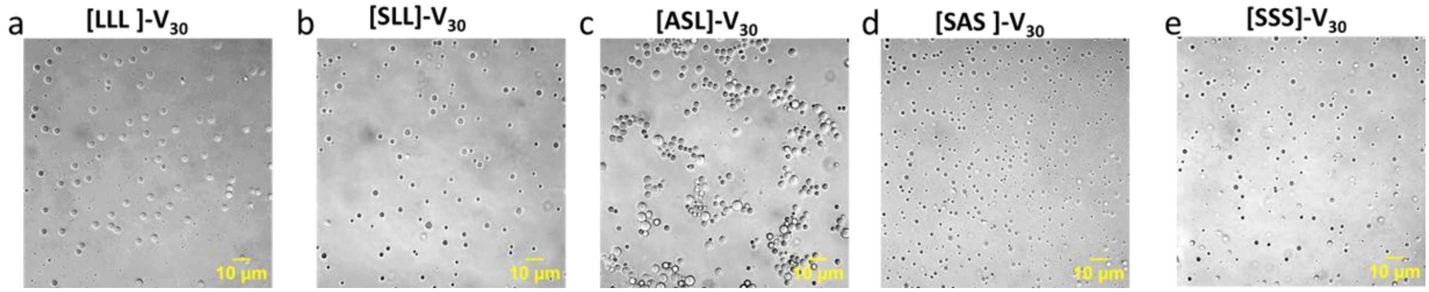

**Supplementary Figure 21. DIC micrographs of unlipidated controls.** **a,** [LLL]-$V_{30}$; **b,** [SLL]-$V_{30}$; **c,** [ASL]-$V_{30}$; **d,** [SAS]-$V_{30}$. **e**, [SSS]-$V_{30}$. In the absence of N-myristoylation, all variants formed liquid droplets, including sequences that drive fiber formation when lipidated. Samples were prepared at 50 µM in PBS supplemented with 1 M NaCl to promote phase-separation at 37 °C. Scale bars, 10 µm.

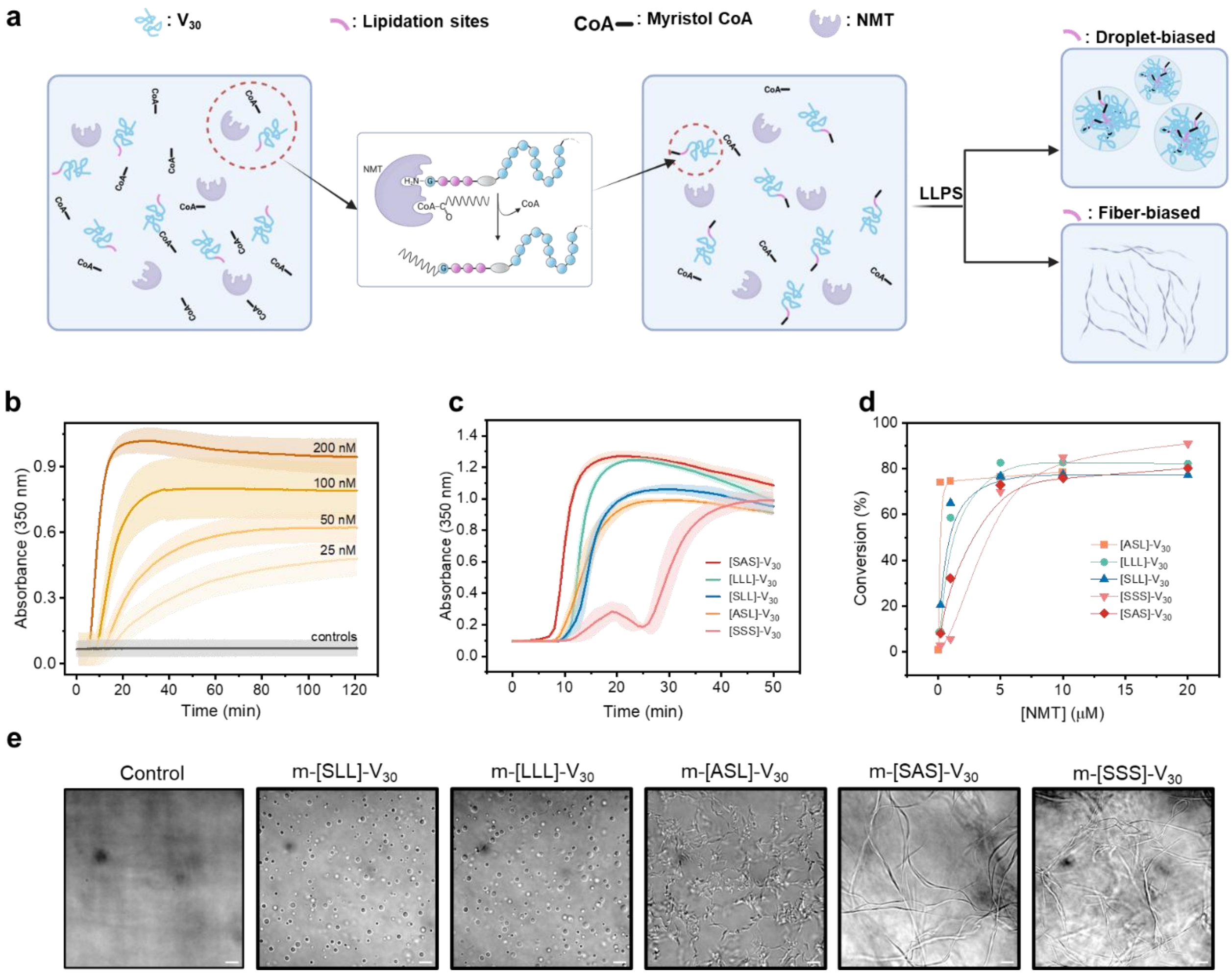

**Supplementary Figure 22. Enzymatic lipidation drives isothermal phase separation and programmable material states. a,** Schematic of fuel-driven assembly. NMT-mediated transfer of myristoyl-CoA to the ELP scaffold triggers an isothermal transition into liquid droplets or solid fibers depending on the lipidation site. **b,** Kinetics for the [ASL]-$V_{30}$ phase separation monitored by turbidity ($A_{350}$) as a function of NMT concentration (25–200 nM). The solid grey line represents the mean value of negative controls lacking either NMT, ELP, or myr-CoA. **c,** Real-time turbidity traces for five representative variants undergoing fuel-driven assembly. **d,** Enzymatic conversion efficiency quantified by RP-HPLC. **e,** End-point DIC micrographs of fuel-driven assemblies showing similar morphologies as thermodynamic assemblies, indicating the condensate properties are sequence-encoded. Control image is for [ASL]-$V_{30}$ without adding NMT. Data in b and c are mean ± s.d. (n = 3 independent experiments). Data in d are from a representative experiment, Solid lines are provided to guide the eye. Scale bars, 10 μm.

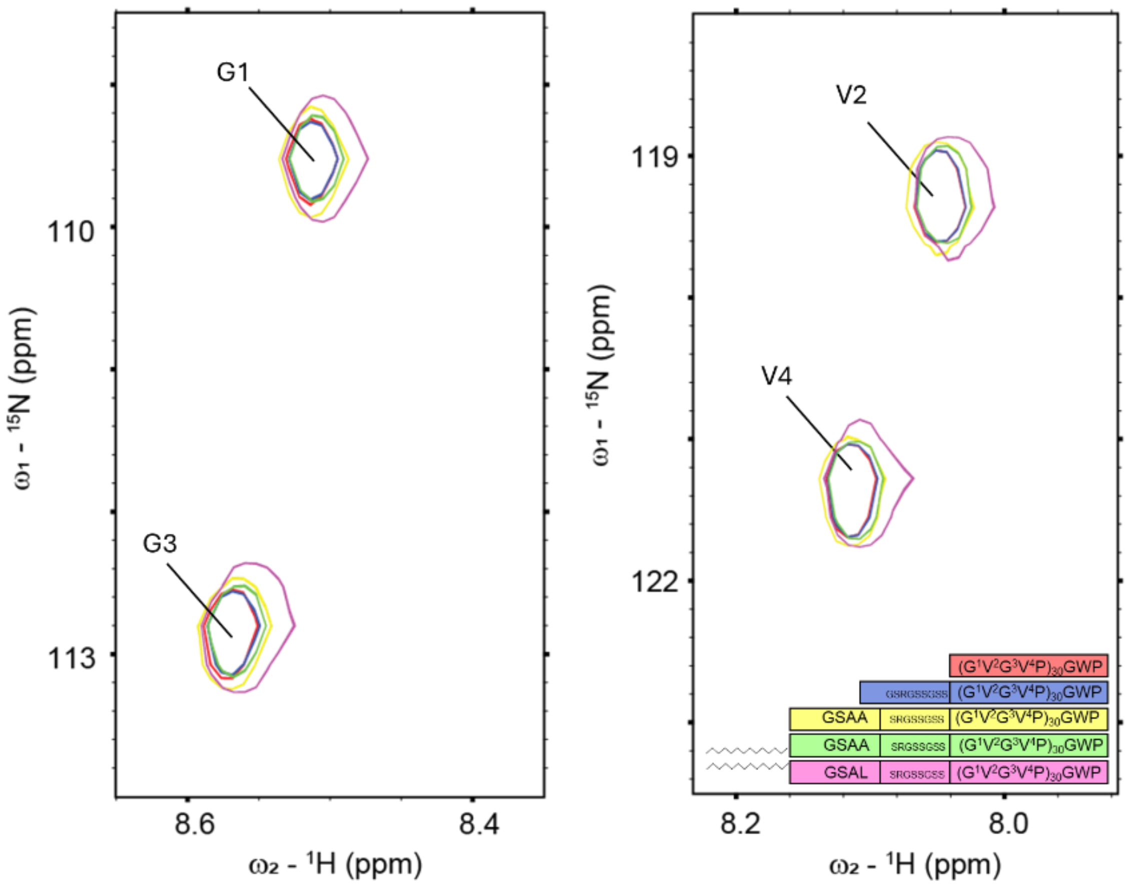

**Supplementary Figure 23. Spectroscopic verification of scaffold conformational invariance.** Superimposed $^{1}H$–$^{15}N$ HSQC spectra (15 °C) of the core scaffold $V_{30}$ (red), Linker-$V_{30}$ (blue), [SAA]-$V_{30}$ (yellow), m-[SAA]$V_{30}$ (lime green), and m-[SAL]-$V_{30}$ (magenta)). Labels denote backbone amides of the $(GVGVP)_{30}$ repeat. The spectral overlap confirms that N-terminal extensions and lipidation do not perturb the global disordered ensembles.

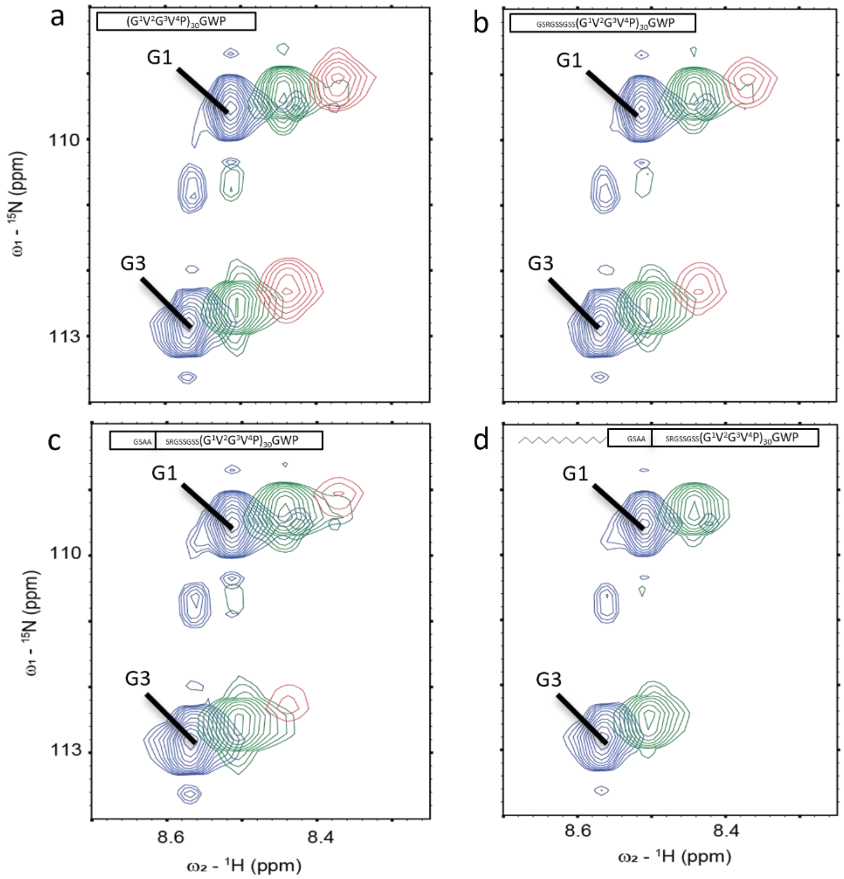

**Supplementary Figure 24. NMR monitoring of phase transition onset.** Overlay of $^1H$–$^{15}N$ HSQC spectra acquired at 15 °C (blue), 25 °C (green), and 35 °C (red) for $V_{30}$ (**a**), Linker-$V_{30}$ (**b**), [SAA]-$V_{30}$ (**c**), and m-[SAA]-$V_{30}$ (**d**). For all constructs, the transition to the phase-separated state results in signal attenuation and eventual disappearance, caused by the slow molecular tumbling of phase-separated assemblies. Consistent with its lower LCST, the lipidated variant (**d**) exhibits complete signal loss at 35 °C, whereas the signal loss for non-lipidated controls (**a**–**c**) occurs at 45 °C.

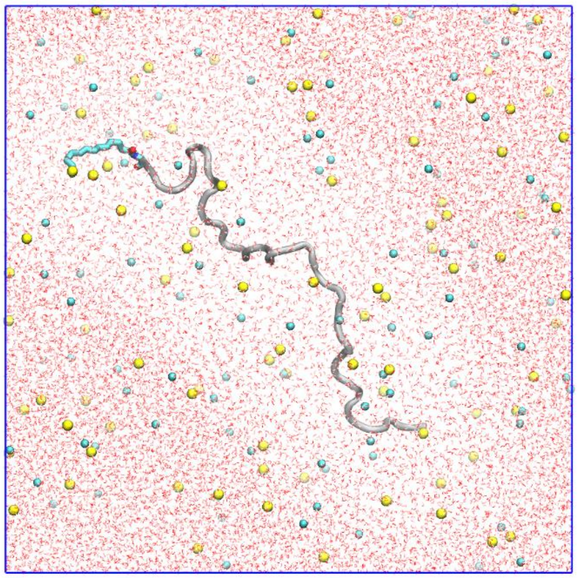

**Supplementary Figure 25. Representative image of the simulation box used for single-chain simulations.** Snapshot of a solvated simulation system containing a myristoylated single-chain protein (shown in silver, with the lipid tail in cyan) in explicit solvent. Water molecules are represented as red and white sticks, $Na^{+}$ and $Cl^{-}$ ions are shown as yellow and cyan spheres. The image illustrates the periodic boundary conditions and the solvation environment in which conformational sampling and structural analyses were performed.

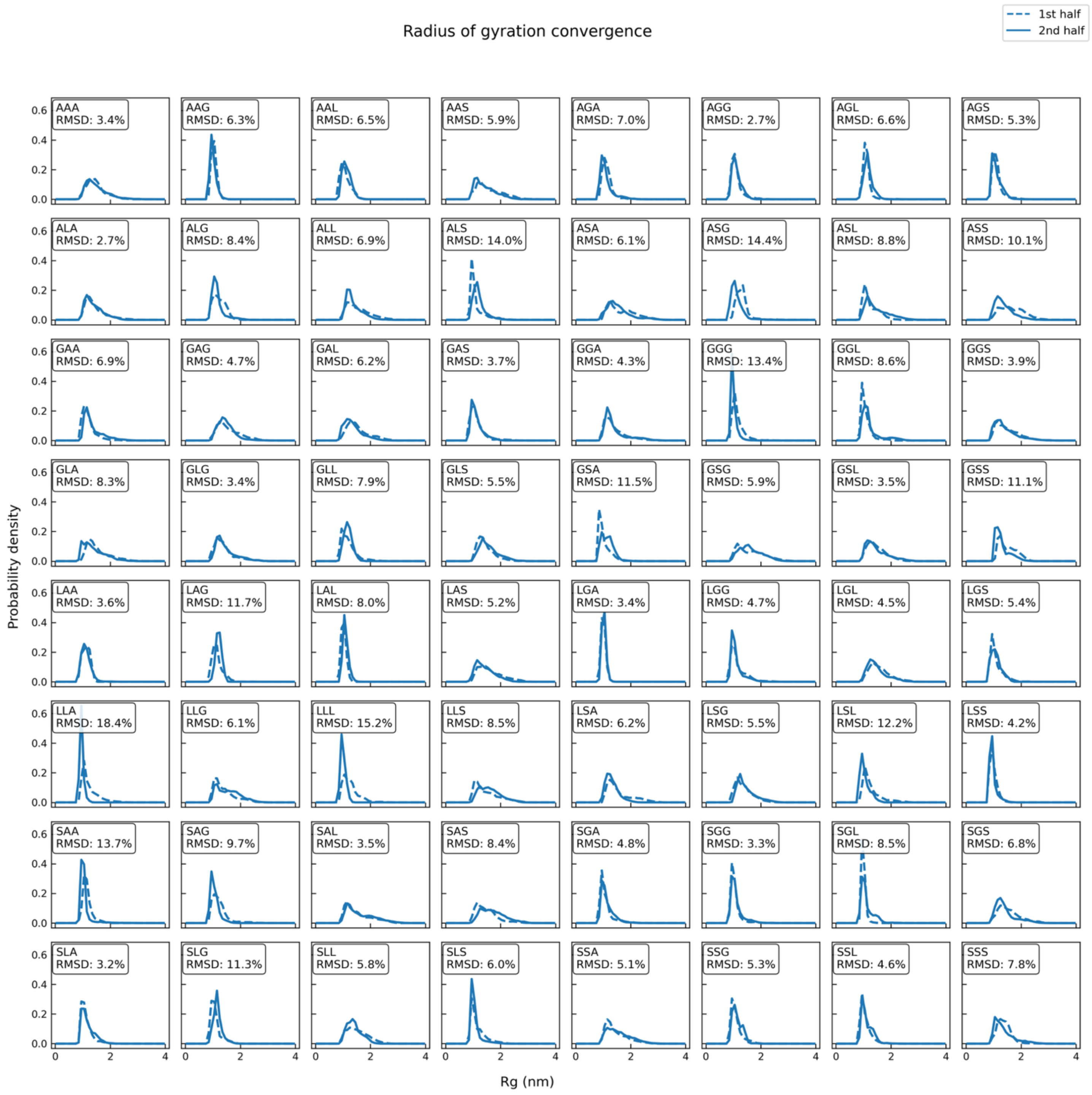

**Supplementary Figure 26. Convergence of radius of gyration (*R*g) distributions across sequences.** Each subplot compares normalized histograms of *R*g values for a single sequence, calculated from into first (100–550 ns, dashed line) and second (550–1000 ns, solid line) halves of a 1µs single-chain MD simulation. The x-axis represents *R*g (nm), and the y-axis denotes normalized probability. The percentage shown in each panel indicates the relative RMSD between the two histograms, providing a quantitative measure of convergence. Lower values reflect better agreement between the two halves.

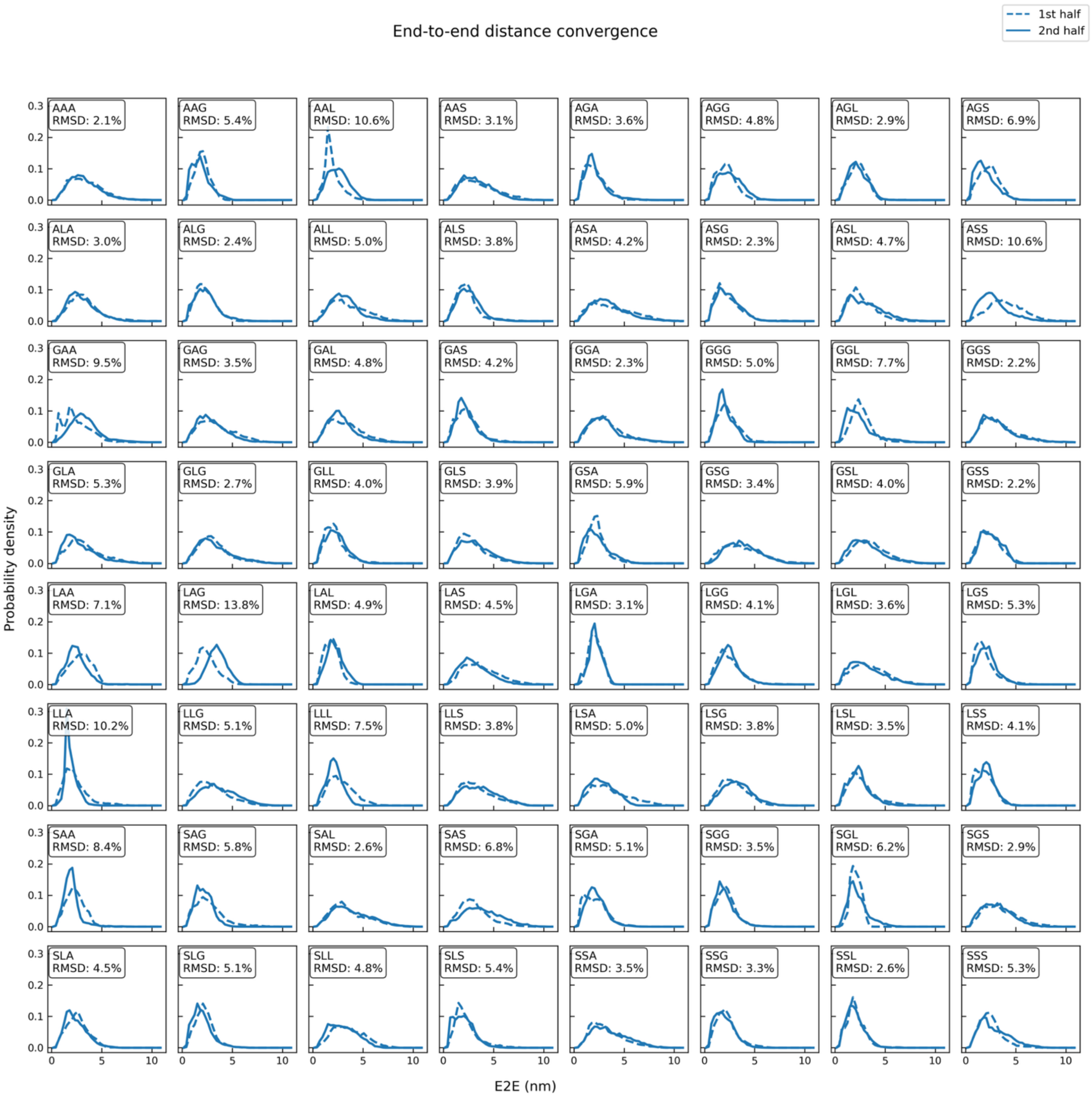

**Supplementary Figure 27. Convergence of end-to-end distance (E2E) distributions across sequences.** Normalized histograms of E2E distances are shown for each sequence, split into first (100–550 ns, dashed line) and second (550–1000 ns, solid line) halves of the simulation. The x-axis corresponds to E2E distance (nm), and the y-axis shows the normalized probability. Relative RMSD values between the two distributions are reported in each subplot to assess convergence.

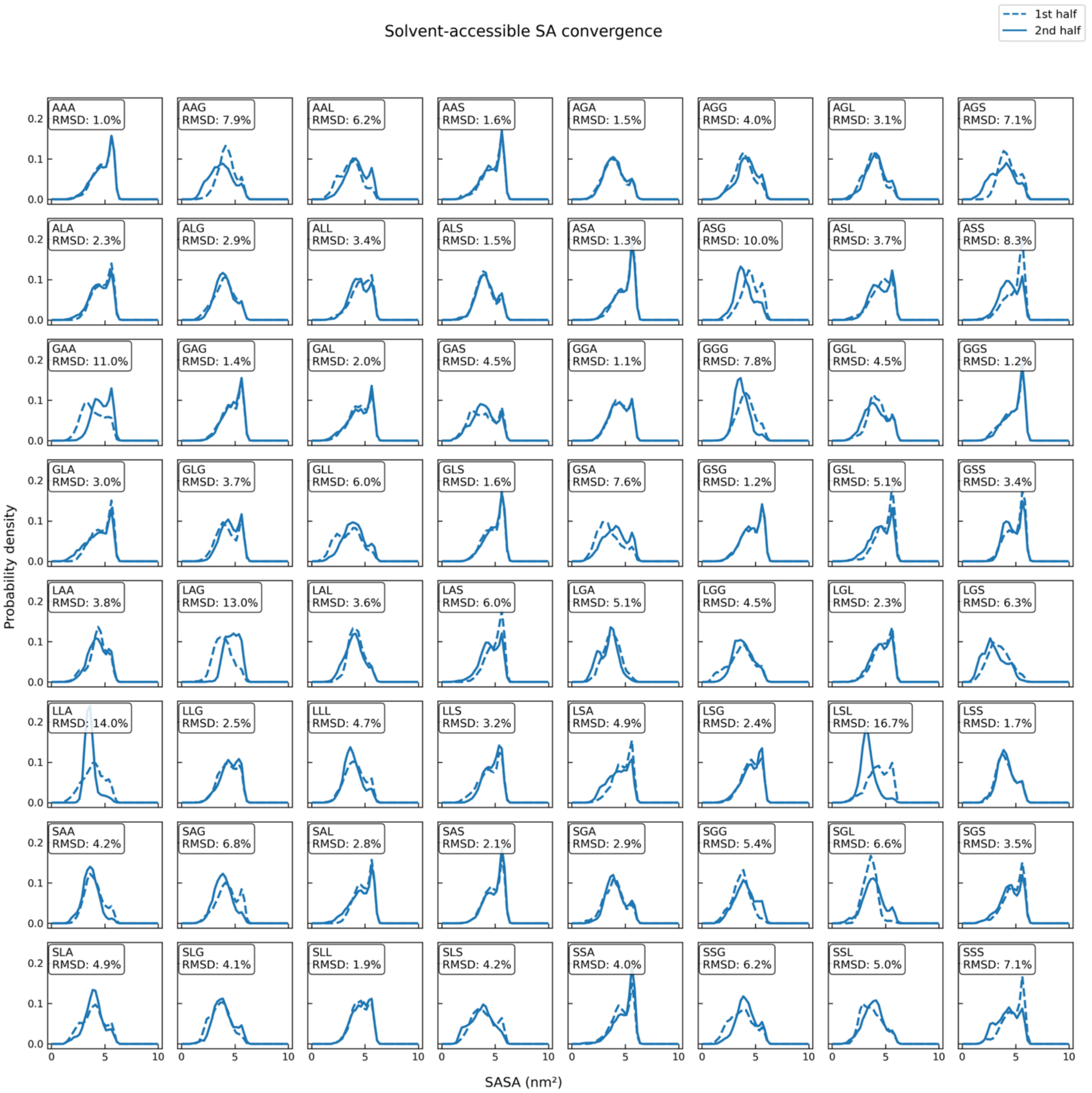

**Supplementary Figure 28. Convergence of solvent-accessible surface area (SASA) distributions across sequences.** Subplots depict comparisons of SASA histograms for the myristoylated N-terminal glycine, computed from into first (100–550 ns, dashed line) and second (550–1000 ns, solid line) halves of the trajectory. The x-axis indicates SASA values (nm²), and the y-axis shows normalized frequencies. The percentage in each panel denotes the relative RMSD between the two histograms for that sequence.

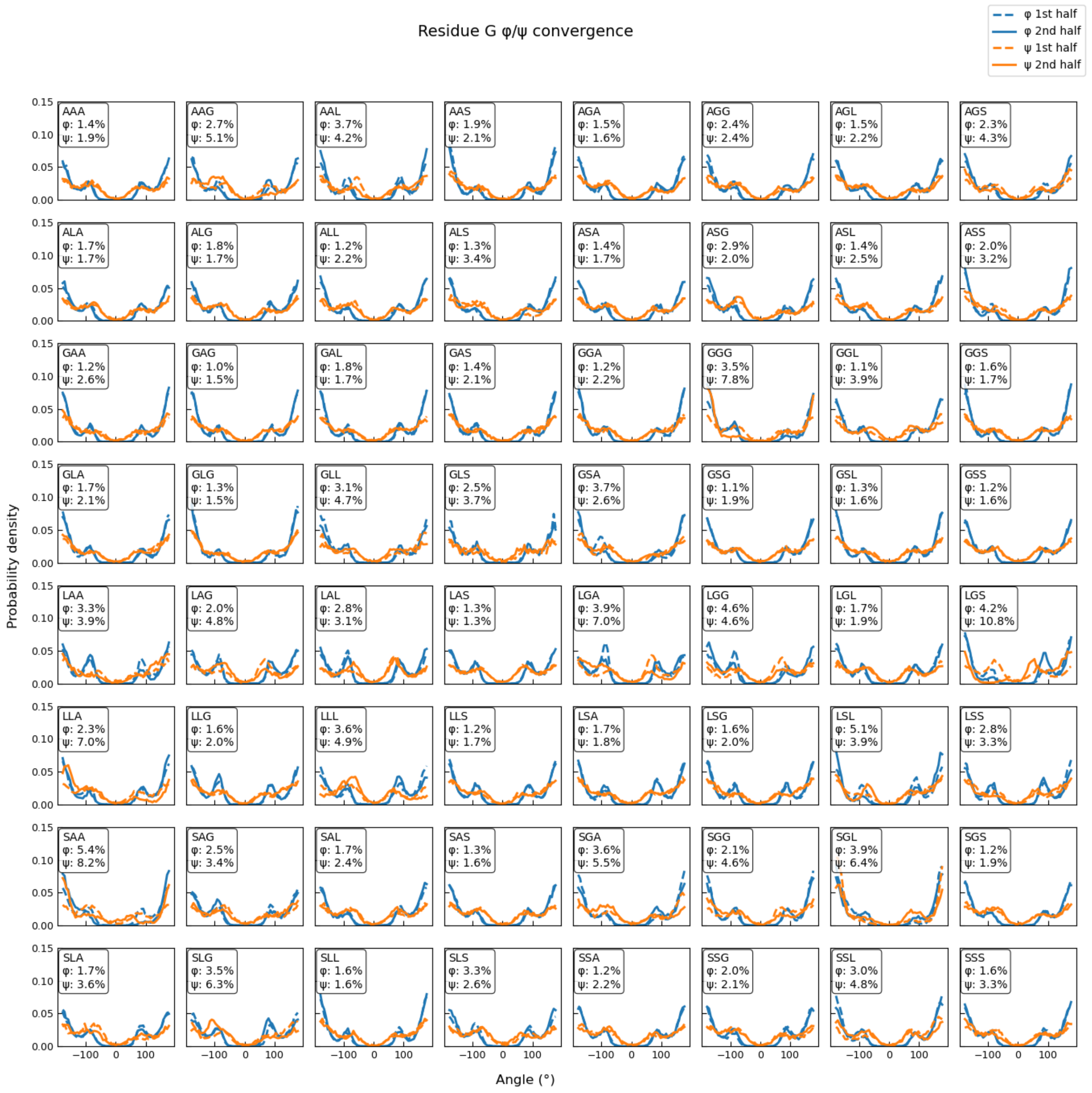

**Supplementary Figure 29. Convergence of backbone dihedral angle distributions for myr-$G_1$ across sequence variants.** Normalized probability density distributions of φ (blue) and ψ (orange) angles for $G_1$ are shown for the first half (dashed lines) and second half (solid lines) of each trajectory. The percentages in each panel quantify the relative RMSD between the halves, as a measure of convergence. The close overlap between dashed and solid curves indicates good convergence of backbone dihedral sampling, while larger deviations highlight sequences with slower conformational equilibration.

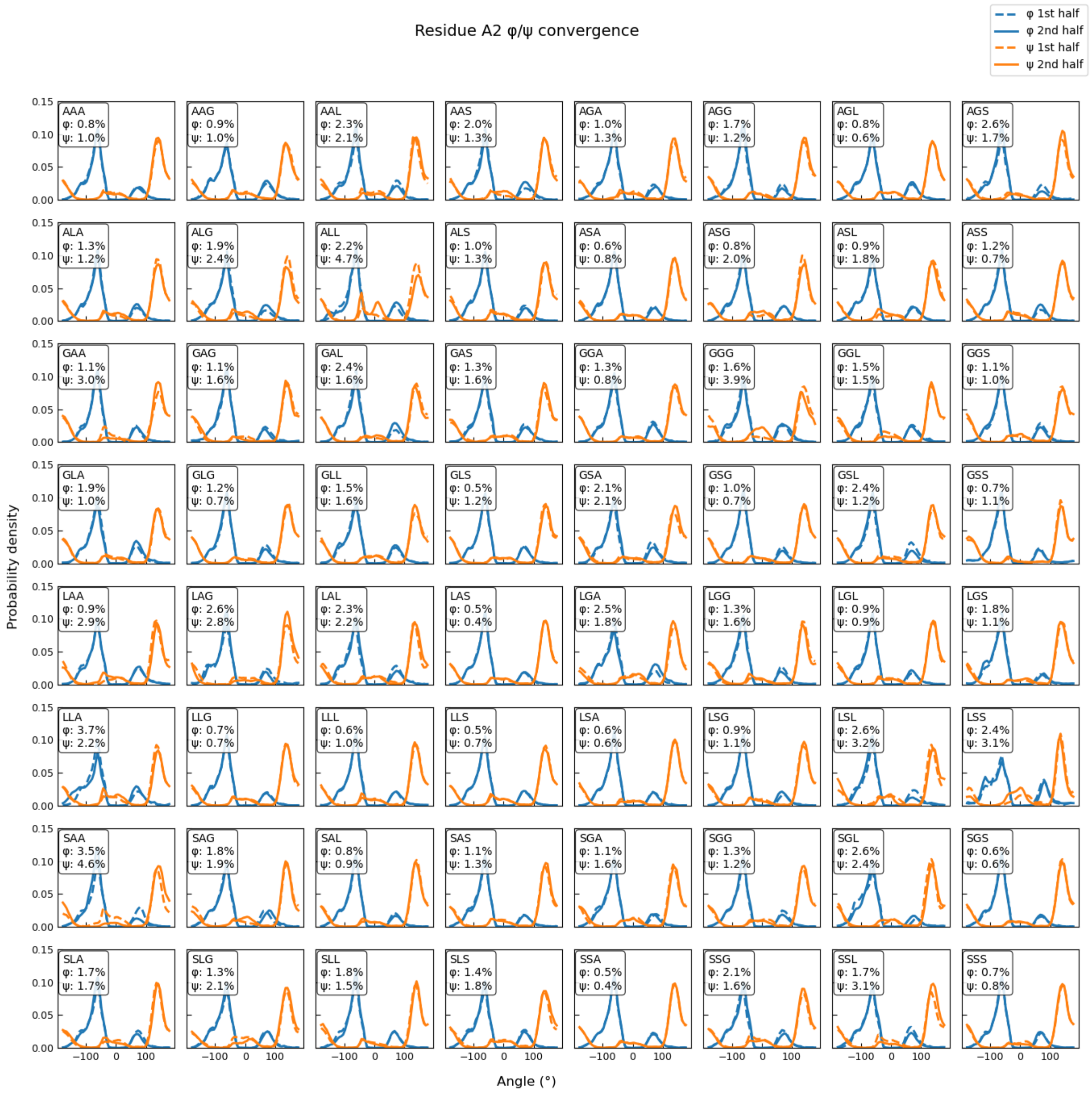

**Supplementary Figure 30**. **Convergence of backbone dihedral angle distributions for $A_2$ across sequence variants.** Normalized probability density distributions of φ (blue) and ψ (orange) angles for residue $A_2$ are shown for the first half (dashed lines) and second half (solid lines) of each trajectory. Each panel corresponds to a distinct tripeptide sequence context (labelled), enabling a systematic comparison of conformational sampling across variants. The percentages reported in each panel quantify the relative RMSD between the first and second halves for φ and ψ, respectively, serving as a measure of convergence. Overall, close overlap between dashed and solid curves indicates good convergence of backbone dihedral sampling, while larger deviations highlight sequences with slower conformational equilibration**.**

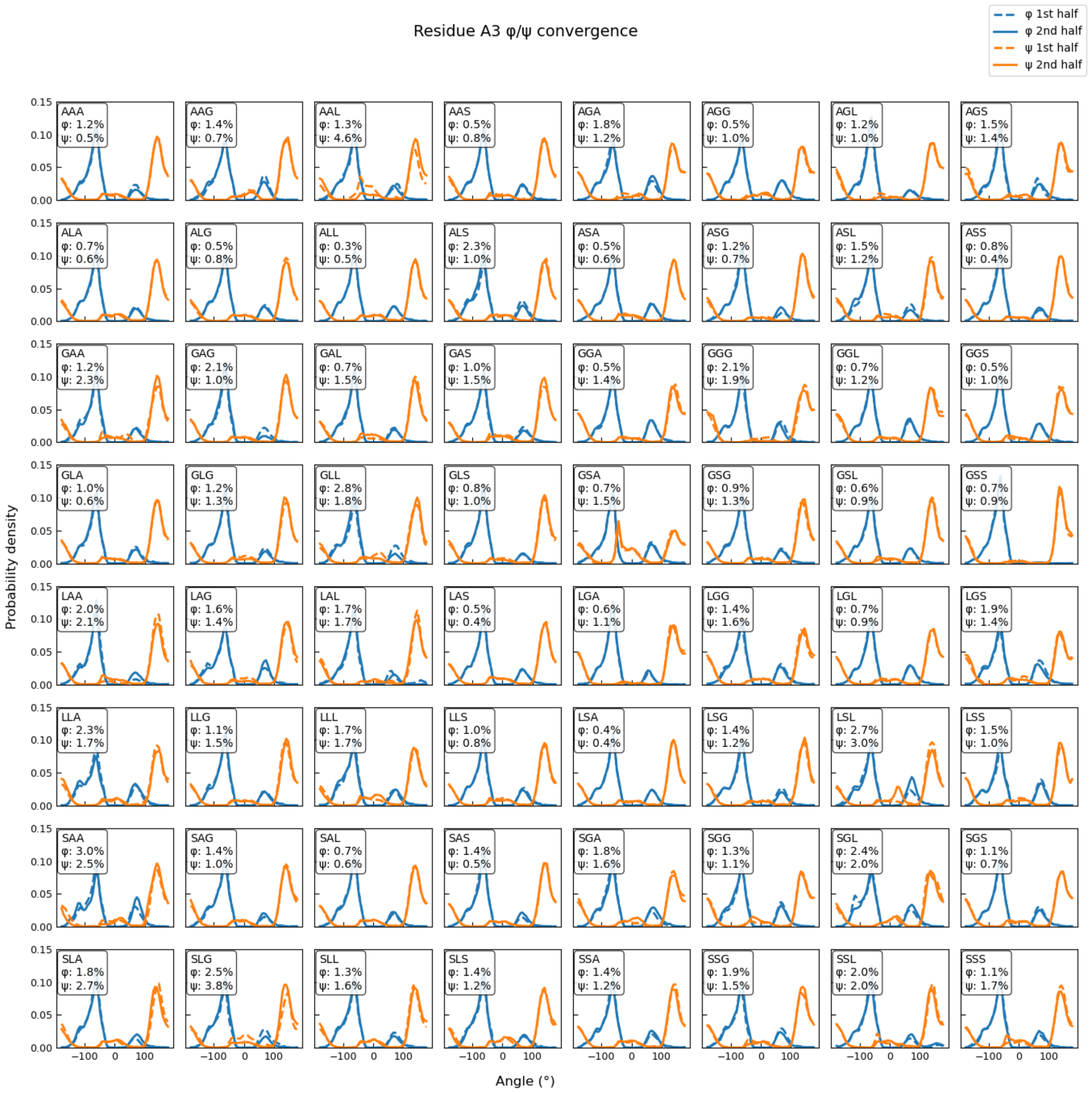

**Supplementary Figure 31. Convergence of backbone dihedral angle distributions for residue A$_3$ across sequence variants.** Normalized probability density distributions of φ (blue) and ψ (orange) angles for residue A$_3$ are shown for the first half (dashed lines) and second half (solid lines) of each trajectory. Each panel corresponds to a distinct tripeptide sequence context (labelled), enabling a systematic comparison of conformational sampling across variants. The percentages reported in each panel quantify the relative RMSD between the first and second halves for φ and ψ, respectively, serving as a measure of convergence. Overall, close overlap between dashed and solid curves indicates good convergence of backbone dihedral sampling, while larger deviations highlight sequences with slower conformational equilibration.

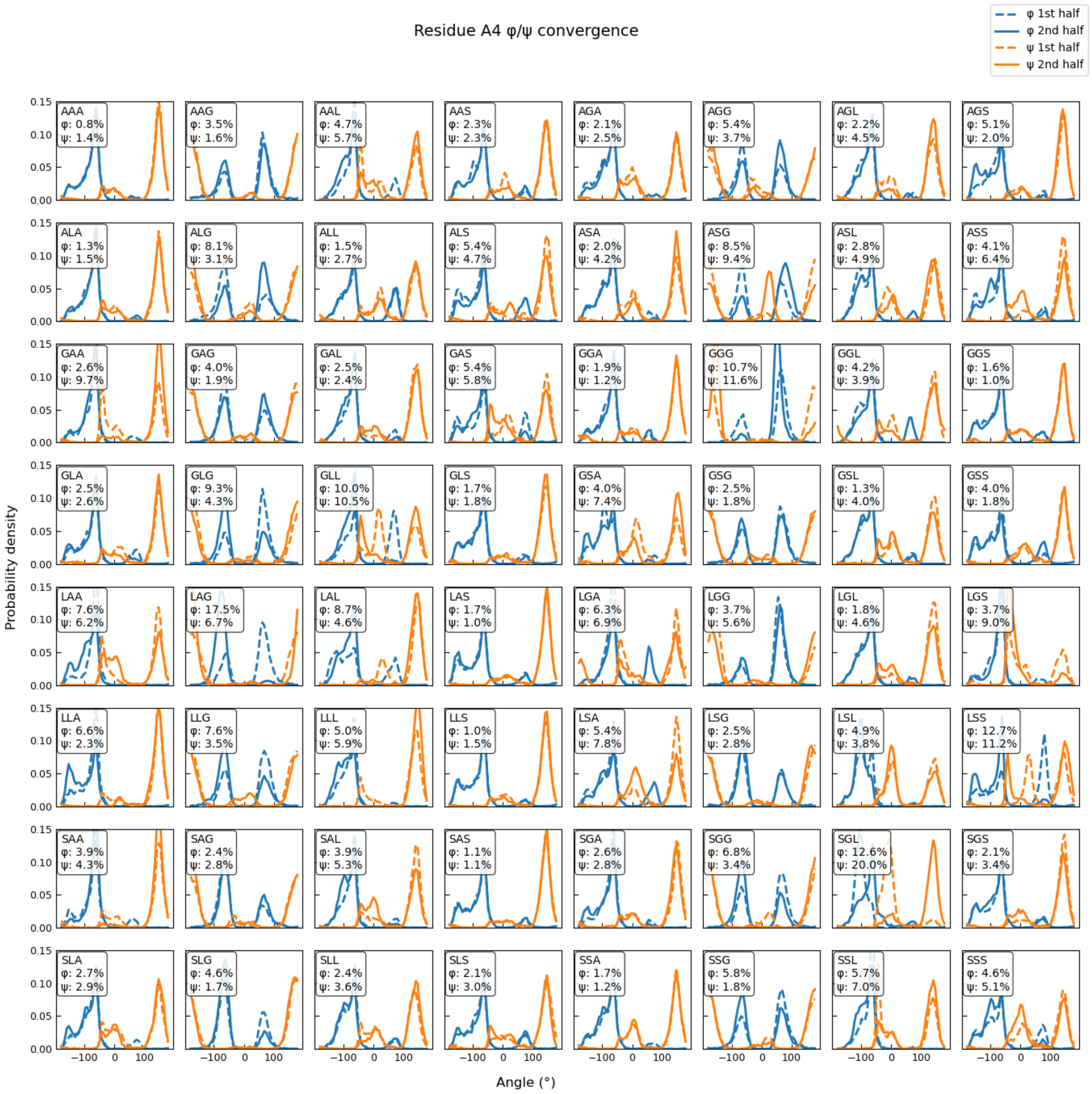

**Supplementary Figure 32. Convergence of backbone dihedral angle distributions A$_4$ across sequence variants.** Normalized probability density distributions of φ (blue) and ψ (orange) angles for residue A$_4$ are shown for the first half (dashed lines) and second half (solid lines) of each trajectory. Each panel corresponds to a distinct tripeptide sequence context (labelled), enabling a systematic comparison of conformational sampling across variants. The percentages reported in each panel quantify the relative RMSD between the first and second halves for φ and ψ, respectively, serving as a measure of convergence. Overall, close overlap between dashed and solid curves indicates good convergence of backbone dihedral sampling, while larger deviations highlight sequences with slower conformational equilibration.

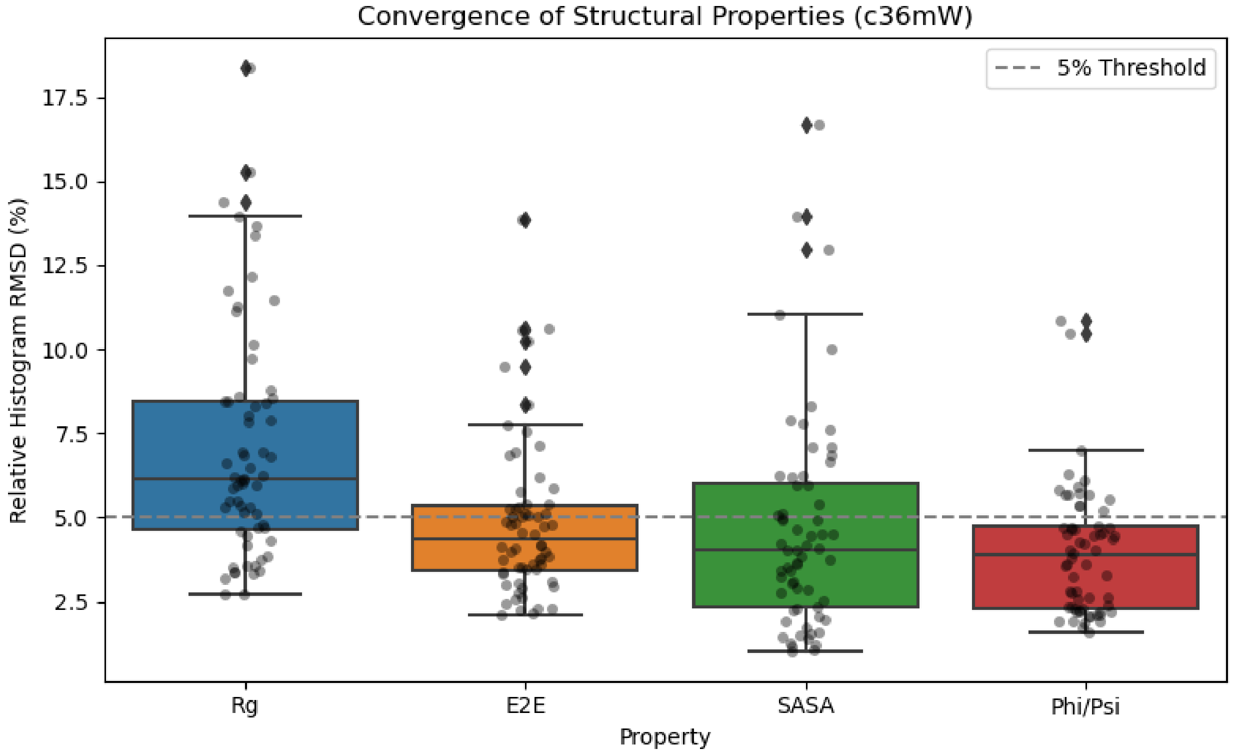

**Supplementary Figure 33. Convergence of key simulated structural properties across the lipidation-site library.** Box-and-swarm plots show the distribution of relative histogram RMSD ($rel_{RMSD}$, expressed as a percentage) between the first-half and second-half trajectories for four key properties: radius of gyration (*R*g), end-to-end distance (E2E), solvent-accessible surface area of the lipidated residue (SASA), and combined φ/ψ angle distributions (averaged over residues 2–11). Individual grey points represent $rel_{RMSD}$ values for each sequence; boxes indicate the interquartile range with the median marked by a horizontal line; whiskers extend to 1.5 × IQR. The dashed grey line denotes a 5% $rel_{RMSD}$ threshold used as the nominal convergence criterion. All properties except RG show median $rel_{RMSD}$ values below this threshold, indicating satisfactory convergence across the dataset.

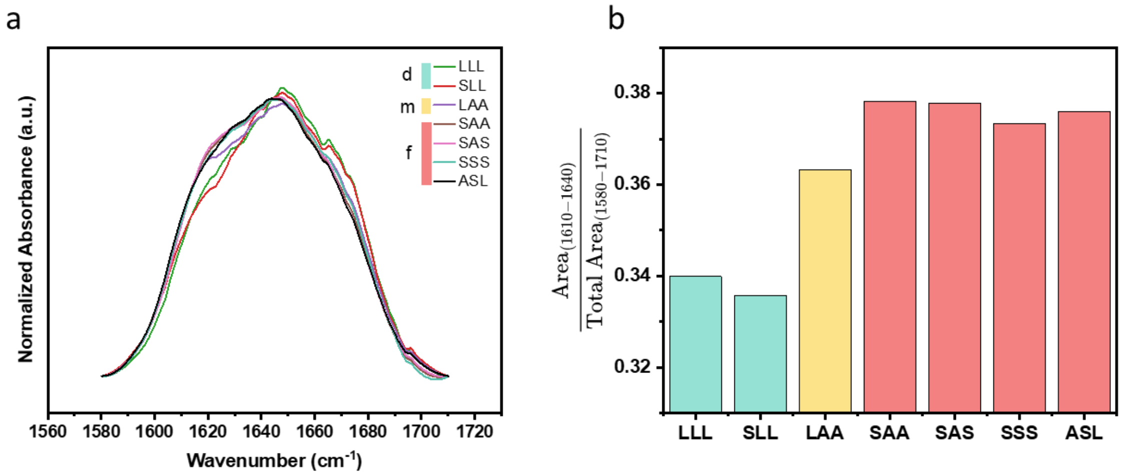

**Supplementary Figure 34. FTIR spectroscopy reveals elevated β-sheet content in fibrillar assemblies. a**, Normalized FTIR spectra (Amide I, 1580–1710 $cm^{-1}$) for representative droplet- (teal), metastable- (yellow), and fiber-forming (red) variants. **b,** Relative β-sheet content quantified as the ratio of 1610–1640 $cm^{-1}$ area to total Amide I area. Fiber-forming variants display significantly higher β-sheet content.

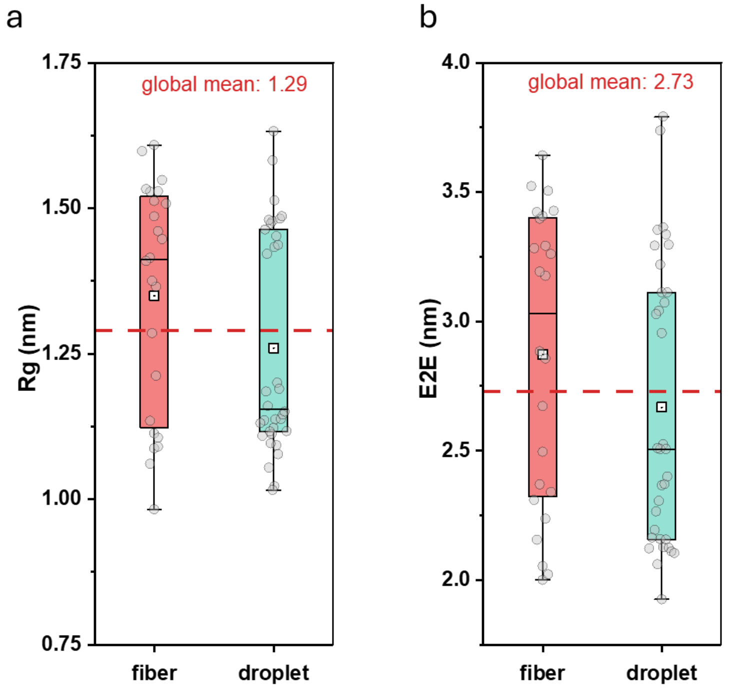

**Supplementary Figure 35. Single-chain conformational properties across distinct phase states.** Box-and-whisker plots show distributions of **a,** radius of gyration (Rg); and **b,** end-to-end distance (E2E) for chains classified as fiber, droplet, and metastable (MS). Individual points represent sequence-resolved measurements. Red dashed lines indicate the global mean value for each metric across all sequences. Boxes denote the interquartile range with the median shown as a horizontal line, mean as square symbol; whiskers extend to 1.5× the interquartile range.

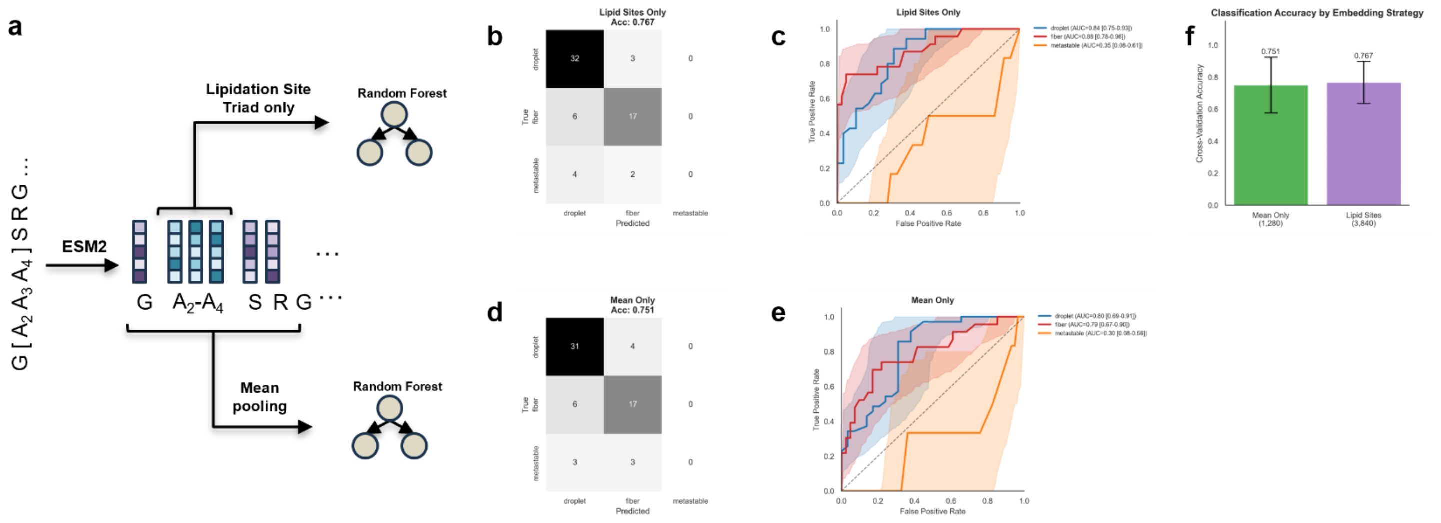

**Supplementary Figure 36. ESM2 embeddings distinguish droplet from fiber states but not metastable intermediates. a,** Classification pipeline. Protein sequences are encoded using ESM2 to generate per-residue embeddings. These are processed using either lipidation-site or mean-pooled sequence representations, then classified using random forest models. **b,c,** Confusion matrix and ROC curves for lipidation-site embeddings; **d,e,** Confusion matrix and ROC curves for mean-pooled embeddings. Both strategies show effective separation of droplet versus fiber states, but poor identification of metastable states. **f,** Comparison of overall accuracy between the two strategies; error bars represent two standard deviations (mean ± 2 s.d.).

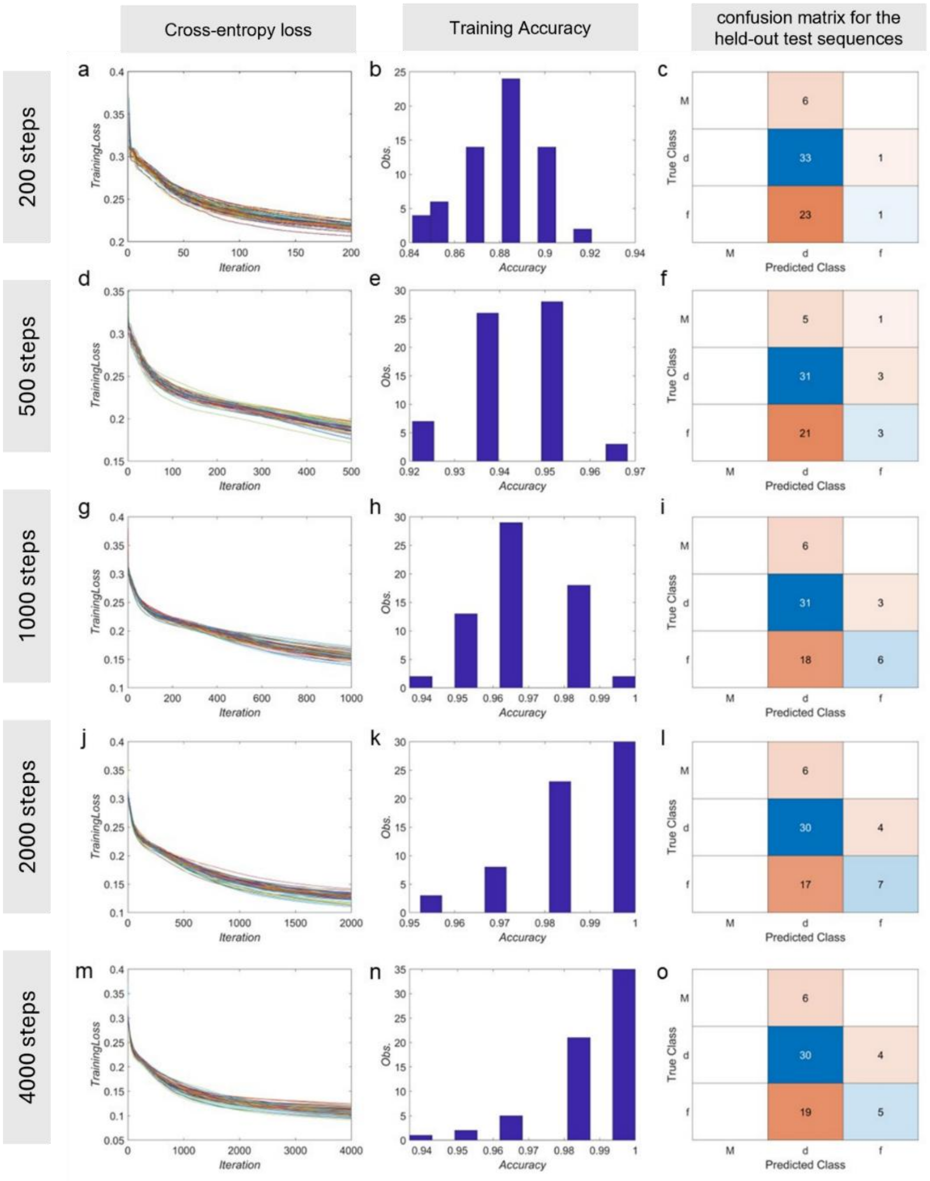

**Supplementary Figure 37. Training dynamics and generalization performance under leave-one-out cross-validation (LOOCV).** Left column (**a, d, g, j, m**): Evolution of training cross-entropy loss as a function of iteration for the 64 independently trained models, each trained on 63 sequences. Middle column (**b, e, h, k, n**): Histograms of the final training accuracy across the 64 models. Right column (**c, f, i, l, o**): Aggregated confusion matrices for the held-out test sequences from the LOOCV evaluation, summarizing sequence-level prediction performance. Together, these results reveal a non-monotonic dependence of generalization performance on training depth, with intermediate iteration limits (1,000–2,000 steps) yielding improved test accuracy relative to both undertrained and over-optimized regimes.

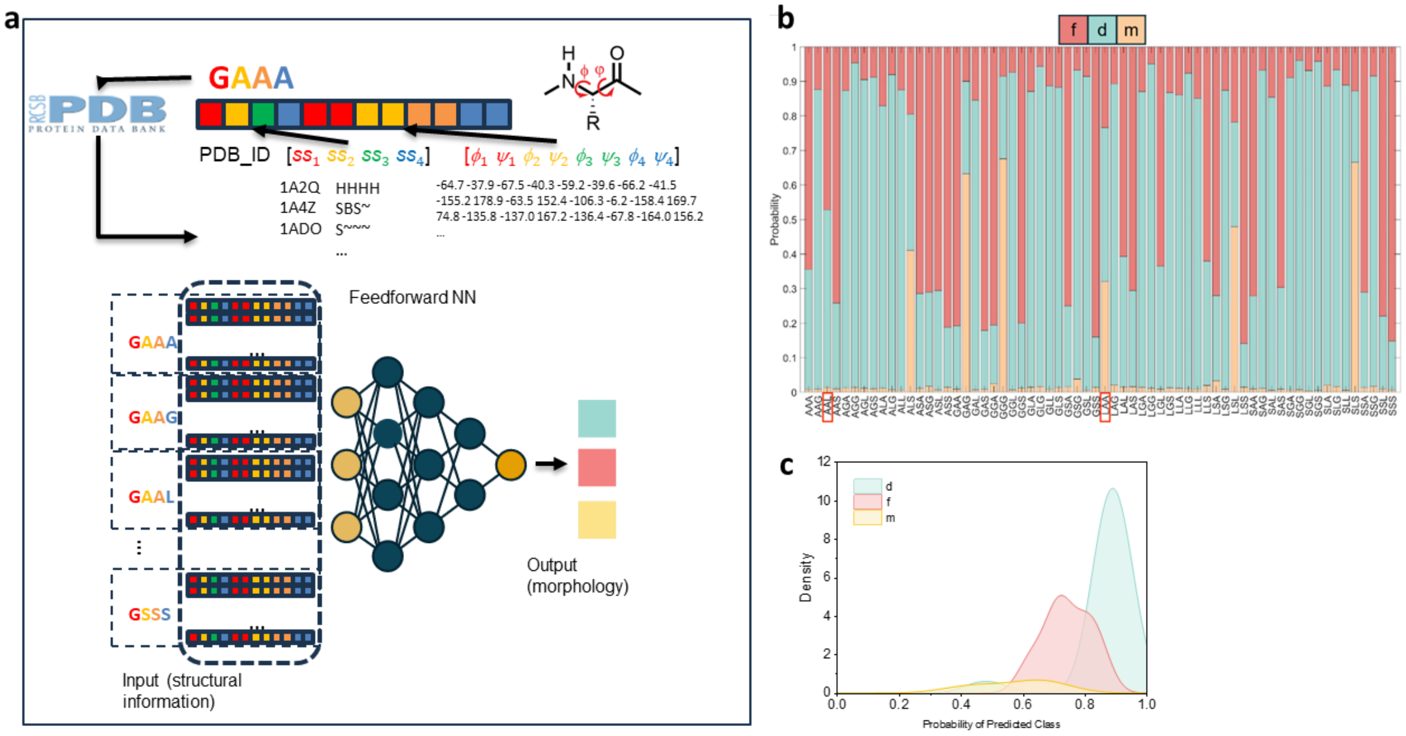

**Supplementary Figure 38. Neural network architecture and training set performance. a,** Schematic of the trilayer feedforward neural network. Input vectors consist of residue-level secondary structure and backbone dihedral angle statistics (ϕ/ψ) derived from PDB data for the lipidation site (Gly1+triad). **b,** Stacked bar chart showing the assembly state probabilities for the 64-sequence training library. The model successfully captures the governing patterns of the dataset, correctly identifying the experimentally observed morphologies for 62 of 64 (~97%) constructs. **c,** Kernel Density Estimate (KDE) of classification probabilities. Distinct phases (liquid droplets (cyan) and solid fibers (red)) exhibit sharp peaks at high probability values, indicating the model learned these features with high confidence. The metastable structures (yellow), however, display a broad, diffuse distribution shifted toward lower probability scores, reflecting the model's greater uncertainty in distinguishing these structures even within the training set.

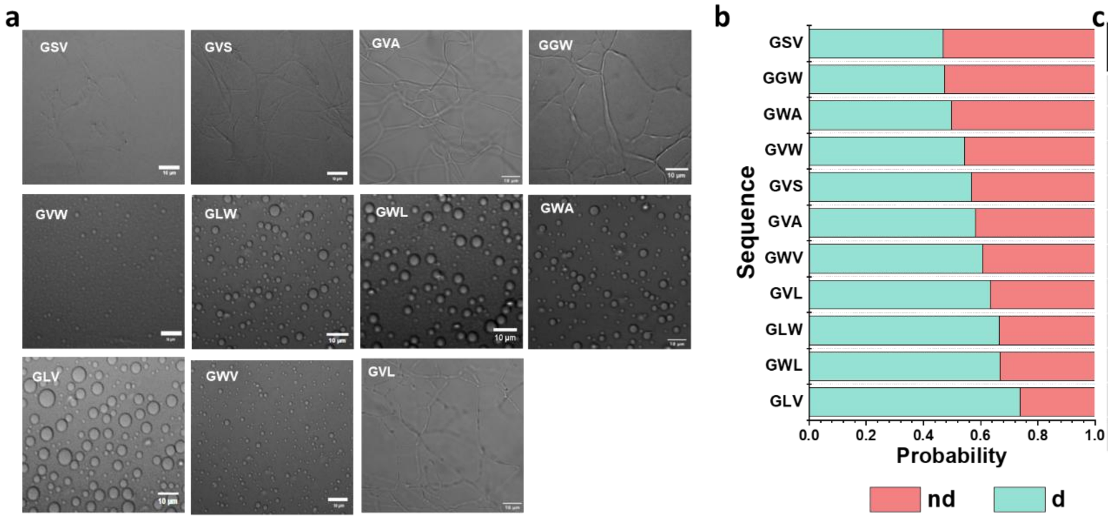

c

| construct | Prediction | Experiments |
|---|---|---|
| GLV | d | d |
| GWL | d | d |
| GLW | d | d |
| GVL | d | d |
| GWV | d | d |
| GVA | d | f |
| GVS | d | f |
| GVW | d | d |
| GWA | d | d |
| GGW | nd | f |
| GSV | nd | f |

**Supplementary Figure 39. Generalization of machine learning predictions to unseen sequences. a,** Representative DIC micrographs for 11 external validation sequences containing V and W residues absent from the training set. **b,c,** Performance of the model trained exclusively on G/A/L/S-containing variants on these unseen sequences. Due to the inherent uncertainty in distinguishing metastable intermediates (Supplementary Figure 38), the predicted probabilities for fibers and metastable states were aggregated into a single "non-droplet" (nd) category.

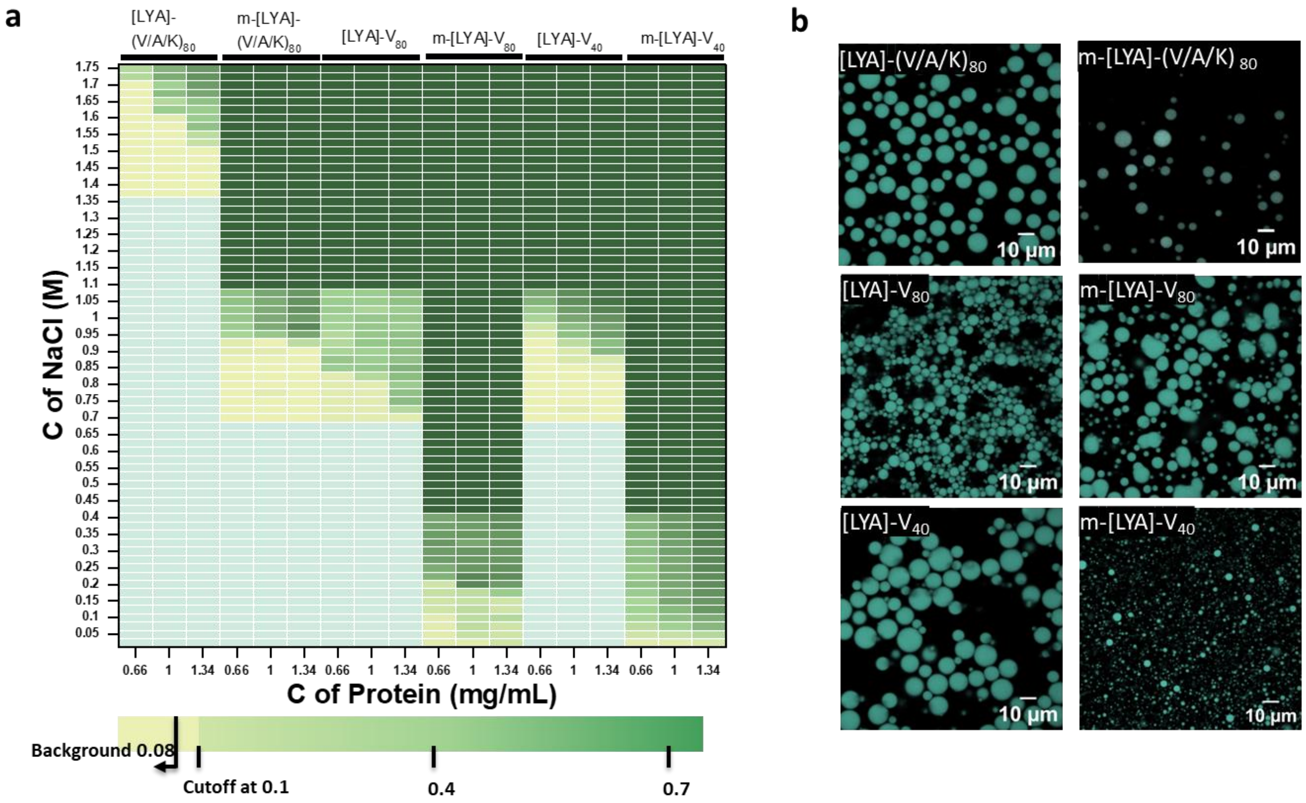

**Supplementary Figure 40. Characterization of phase boundaries and condensate properties of the miscibility library. a,** Partial salt-concentration phase diagrams as a function of protein (0.66–1.34 mg/mL) and NaCl (0.05–1.75 M) concentration, obtained using turbidimetry. The color scale indicates optical density (OD), with phase-boundary defined as salt concentration yielding OD > 0.1. **b,** Representative confocal images of non-lipidated and lipidated constructs, forming liquid condensate at 1.7 M NaCl, above the critical salt concentration for all variants.

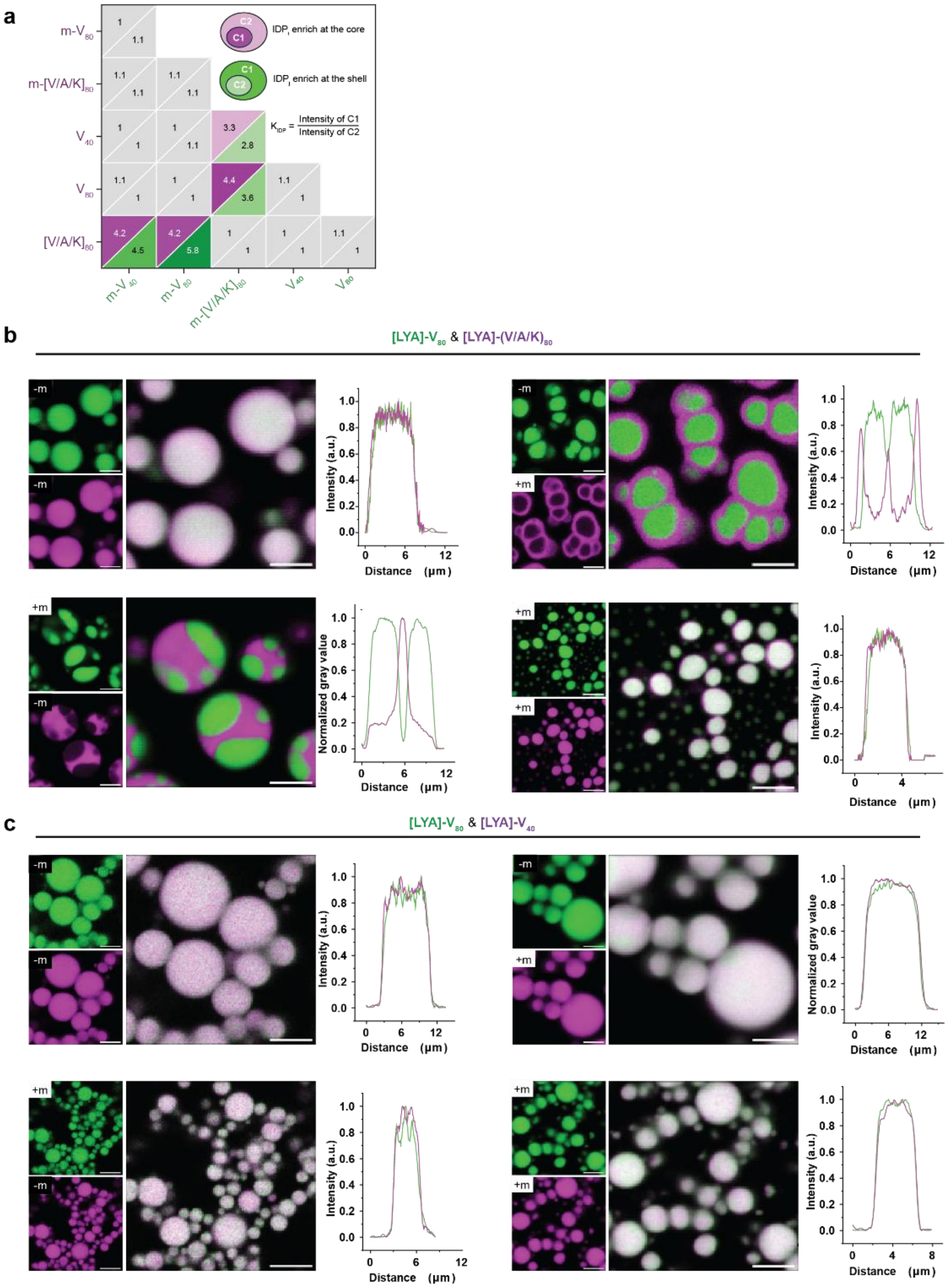

**Supplementary Figure 41. The interplay between the lipid and the guest residue hydropathy regulates condensate miscibility. a,** Heat map of partition coefficients derived from pairwise mixtures across the miscibility library. b, Confocal fluorescence microscopy of pairwise mixtures containing [LYA]-$V_{80}$ (green) and [LYA]-$(V/A/K)_{80}$ (magenta) organized by lipidation state. Symmetric mixtures (both unlipidated or both myristoylated) form miscible condensates with overlapping fluorescence profiles. In contrast, asymmetric mixtures (one component lipidated) demix into multiphase condensates with core-shell architecture. **c,** Confocal fluorescence microscopy of pairwise mixtures containing [LYA]-$V_{80}$ (green) and [LYA]-$V_{40}$ (magenta), organized by lipidation state. In contrast to chemically distinct pairs, chemically similar scaffolds remain miscible across all conditions. Scale bar, 10 μm.

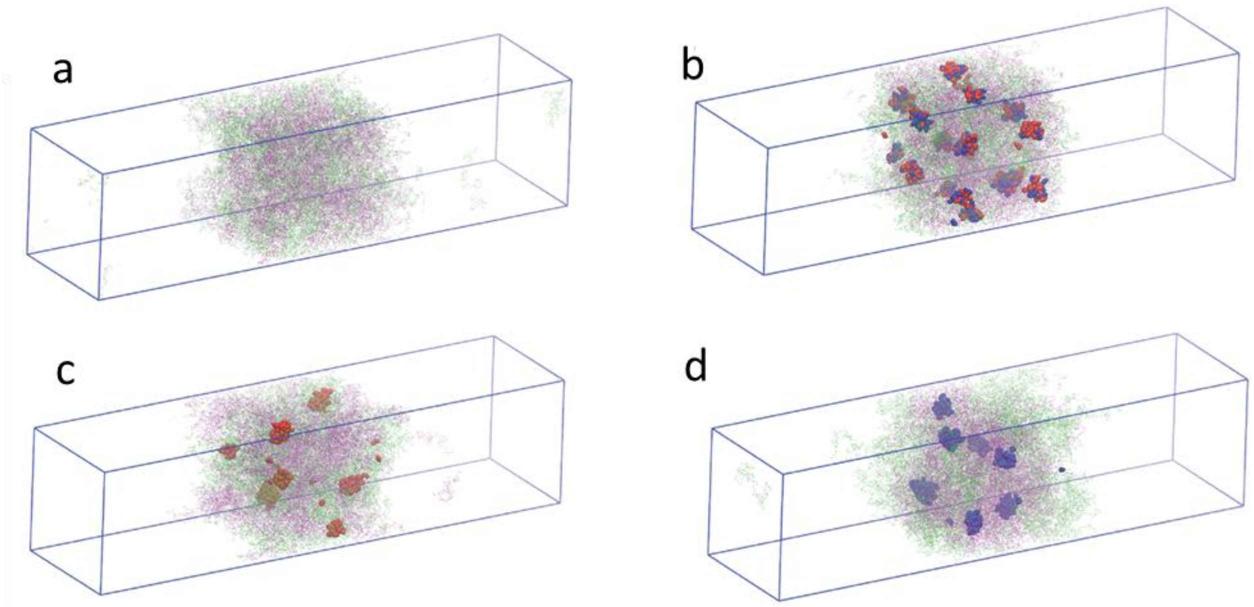

**Supplementary Figure 42. Coarse-grained (CG) simulations of micelle formation in slab geometry for binary mixtures of [LYA]-(V/A/K)$_{40}$ and [LYA]-V$_{40}$ with varied lipidation status. a,** Mixture of unlipidated isoforms: [LYA]-(V/A/K)$_{40}$ and [LYA]-V$_{40}$. **b,** Mixture with both components lipidated: m-[LYA]-(V/A/K)$_{40}$ and m-[LYA]-V$_{40}$. **c, d,** Mixtures with single-component lipidated: m-[LYA]-(V/A/K)$_{40}$ + [LYA]-V$_{40}$ (**c**) and [LYA]-(V/A/K)$_{40}$ + m-[LYA]-V$_{40}$ (**d**). [LYA]-(V/A/K)$_{40}$ chains and their lipid moieties are shown in green and red, respectively, while [LYA]-V$_{40}$ chains and their lipids are shown in purple and blue. Lipidation induces micellar clustering within the dense phase, with the extent and spatial organization of micelles depending on which component is lipidated.

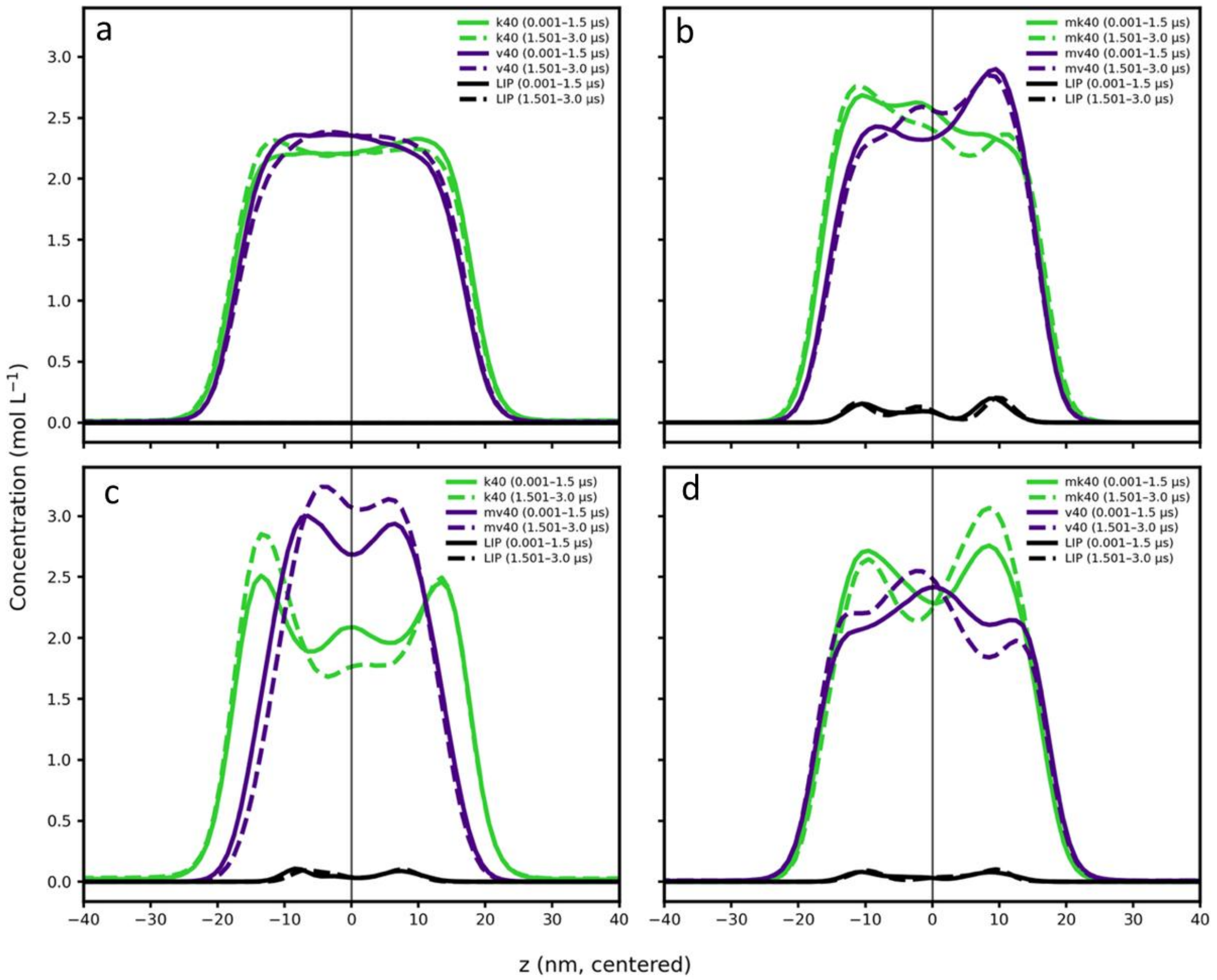

**Supplementary Figure 43. Slab simulation density convergence profiles for binary mixtures of [LYA]-K$_{40}$ and [LYA]-V$_{40}$ with varied lipidation status.** Concentration profiles along the slab normal (z, centered) are shown for: **a**, [LYA]-(V/A/K)$_{40}$ / [LYA]-V$_{40}$; **b,** m-[LYA]-(V/A/K)$_{40}$ / m-[LYA]-V$_{40}$; **c,** [LYA]-(V/A/K)$_{40}$ / m-[LYA]-V$_{40}$; and **d,** m[LYA]-(V/A/K)$_{40}$ / [LYA]-V$_{40}$. Green curves correspond to [LYA]-(V/A/K)$_{40}$ and purple curves to [LYA]-V$_{40}$; solid and dashed lines represent early (0.001–1.5 µs) and late (1.501–3.0 µs) simulation windows, respectively. Black curves show the corresponding lipid density profiles. Overlap and reproducibility of early- and late-time profiles indicate convergence of slab densities, while lipidation-dependent deviations reflect altered spatial organization within the dense phase.

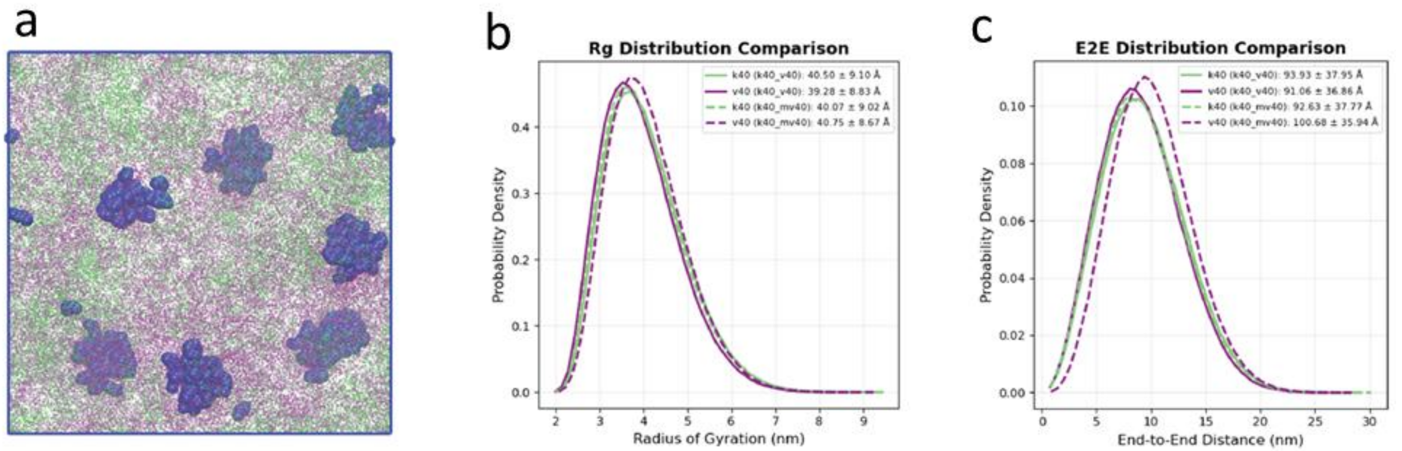

**Supplementary Figure 44. Lipidation-induced micellar organization and peptide conformational expansion in binary mixtures of [LYA]-(V/A/K)$_{40}$ and [LYA]-V$_{40}$. a,** Representative snapshot from coarse-grained simulations of [LYA]-(V/A/K)$_{40}$ / m-[LYA]-V$_{40}$ system, highlighting the formation of lipid-driven micellar patches by lipidated V$_{40}$ chains (blue). The surrounding dense-phase matrix is composed of unlipidated [LYA]-(V/A/K)$_{40}$ chains (green) and [LYA]-V$_{40}$ peptide backbones (purple). **b,** Probability distributions of the peptide radius of gyration (*R*g) for ([LYA]-(V/A/K)$_{40}$ / [LYA]-V$_{40}$) and ([LYA]-(V/A/K)$_{40}$ / m-[LYA]-V$_{40}$) mixtures. *R*g values are computed exclusively from peptide beads, with lipid moieties excluded. Lipidation shifts the *R*g distribution of V$_{40}$ chains toward larger values, consistent with chain expansion. **c,** End-to-end (E2E) distance distributions for peptide chains under the same conditions, computed excluding lipid beads, showing increased sampling of extended conformations upon lipidation. Values in the legends correspond to the mean ± standard deviation of each distribution for the respective sequences and simulation conditions.

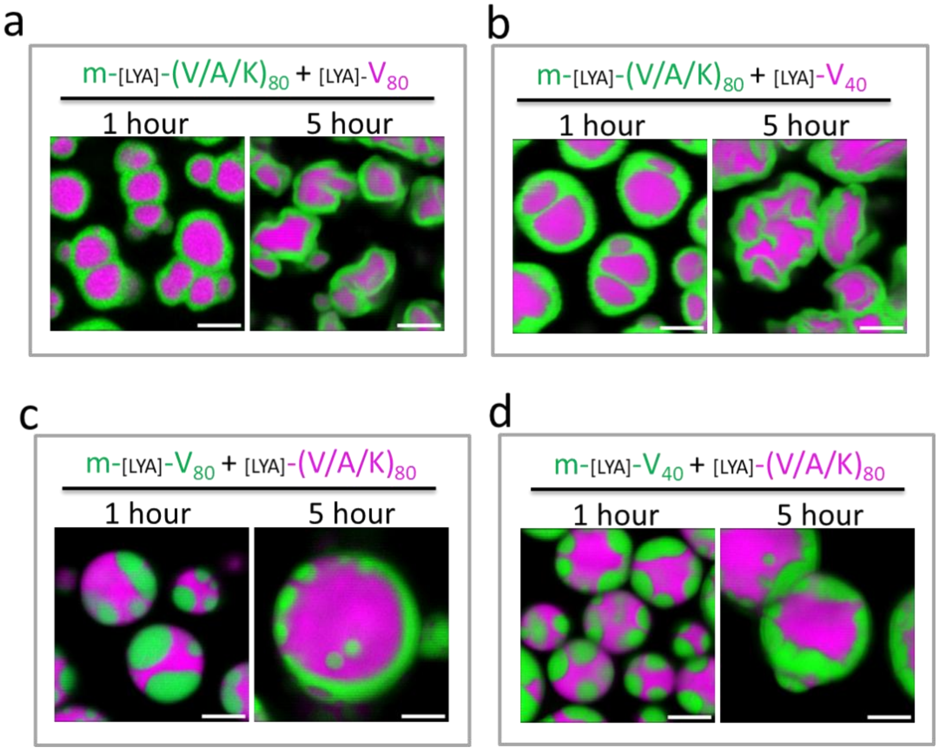

**Supplementary Figure 45. Temporal stability of condensate architecture.** Time-resolved confocal micrographs of select binary mixtures with one lipidated component (green) and one non-lipidated component (magenta) after incubation for 1 and 5 hours. **a,b,** m-[LYA]-$(V/A/K)_{80}$ mixed with [LYA]-$V_{80}$ (**a**) or [LYA]-$V_{40}$ (**b**). **c,d,** [LYA]-$(V/A/K)_{80}$ mixed with m-[LYA]-$V_{80}$ (**c**) or m-[LYA]-$V_{40}$ (**d**). The stability of condensate architectures, notwithstanding minor coarsening of the shell phase, is consistent with the thermodynamic stability of the lipidation-induced immiscibility. Scale bar, 5 μm.

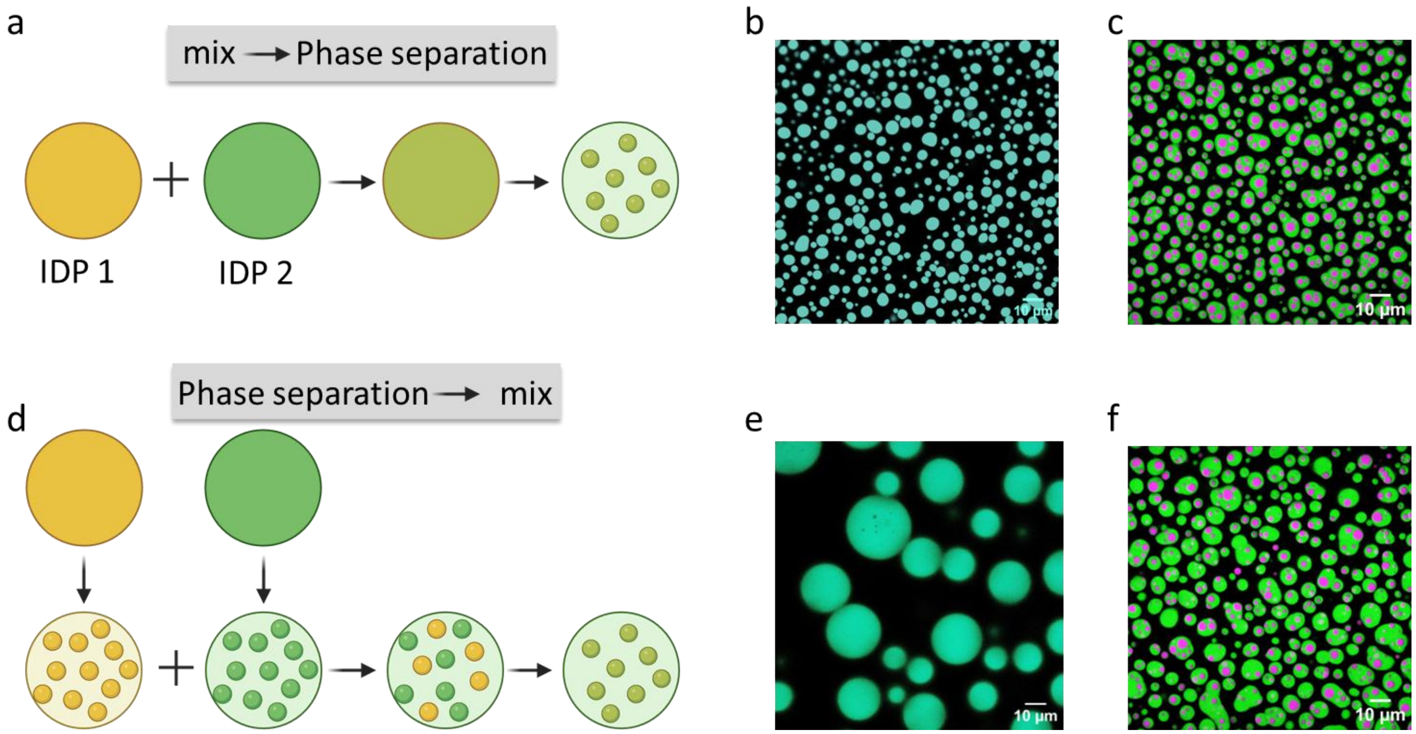

**Supplementary Figure 46. Condensate miscibility is pathway-independent. a–c,** Schematic and results of the co-condensation pathway (mixing soluble precursors). **d–f,** Schematic and results of the condensate mixing pathway. **b, e,** Confocal micrographs of a miscible pair ([LYA]-$(V/A/K)_{80}$ + m-[LYA]-$(V/A/K)_{80}$) formed via co-condensation **(b)** or condensate mixing (**e**). **c, f,** Confocal micrographs of an immiscible pair ([LYA]-$V_{40}$ + m-[LYA]-$(V/A/K)_{80}$) formed via co-condensation (**c**) or condensate mixing (**f**). The identical final states (top vs. bottom) indicate that miscibility is governed by thermodynamic equilibrium rather than the assembly pathway.

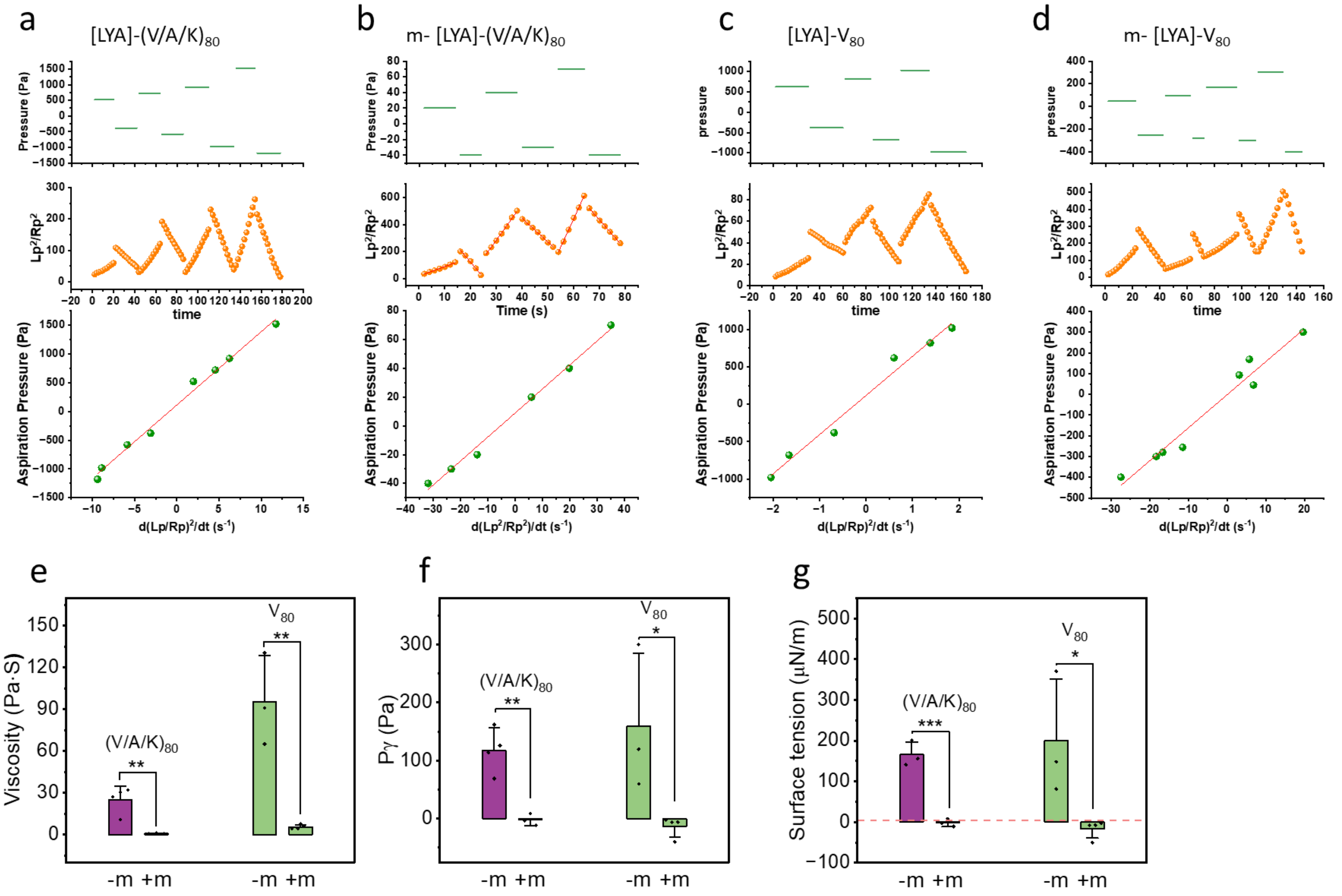

**Supplementary Figure 47. Characterization of condensate material properties via micropipette aspiration (MPA)**. **a–d,** Representative MPA traces for [LYA]-(V/A/K)$_{80}$ (**a**), m-[LYA]-(V/A/K)$_{80}$ (**b**), [LYA]-V$_{80}$ (**c**), m-[LYA]-V$_{80}$ (**d**). Each panel displays the stepwise applied aspiration pressure ($P_{asp}$, top), the time evolution of the normalized projection length squared $(Lp/Rp)^2$, middle), and the linear dependence of aspiration pressure on the expansion rate ($d(Lp/Rp)^2/dt$, bottom). The linearity of the bottom plots confirms Newtonian fluid behavior, where the slope corresponds to dynamic viscosity (4η) and the y-intercept yields the critical pressure ($P\gamma$). (**e–g**) Summary of quantified material properties comparing non-myristoylated (−m) and myristoylated (+m) condensates: **(e)** Viscosity (η), (**f**) Critical pressure ($P\gamma$), and (**g**) Surface tension (γ), dashed line represents theoretical lower limit of the instrument for the dynamic measurement. Myristoylation (+m) significantly reduces viscosity, critical pressure, and surface tension compared to non-lipidated counterparts. Data are mean ± s.d. (n=3). *$P < 0.05$, **$P < 0.01$, ***$P < 0.001$ (one-way ANOVA with Tukey's post-hoc test).

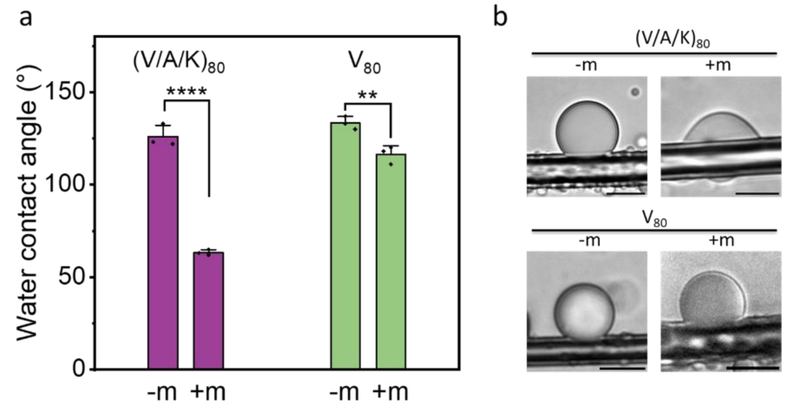

**Supplementary Figure 48. Effect of myristoylation on hydrophilicity and surface wetting of condensates.** **a,** Water contact angle measurements for [LYA]-(V/A/K)$_{80}$ (purple) and [LYA]-V$_{80}$ (green) condensates on a hydrophilic glass surface, comparing non-lipidated (−m) and lipidated (+m) states. The decrease in contact angle indicates increased condensate hydrophilicity, or enhanced interaction with the substrate. **b,** Representative bright-field microscopy images showing the wetting behavior of each sample. Data in (**a**) are mean ± s.d. (n=3). $^{**}P < 0.01$, $^{****}P < 0.0001$ (one-way ANOVA with Tukey's post-hoc test). Scale bars, 10 µm.

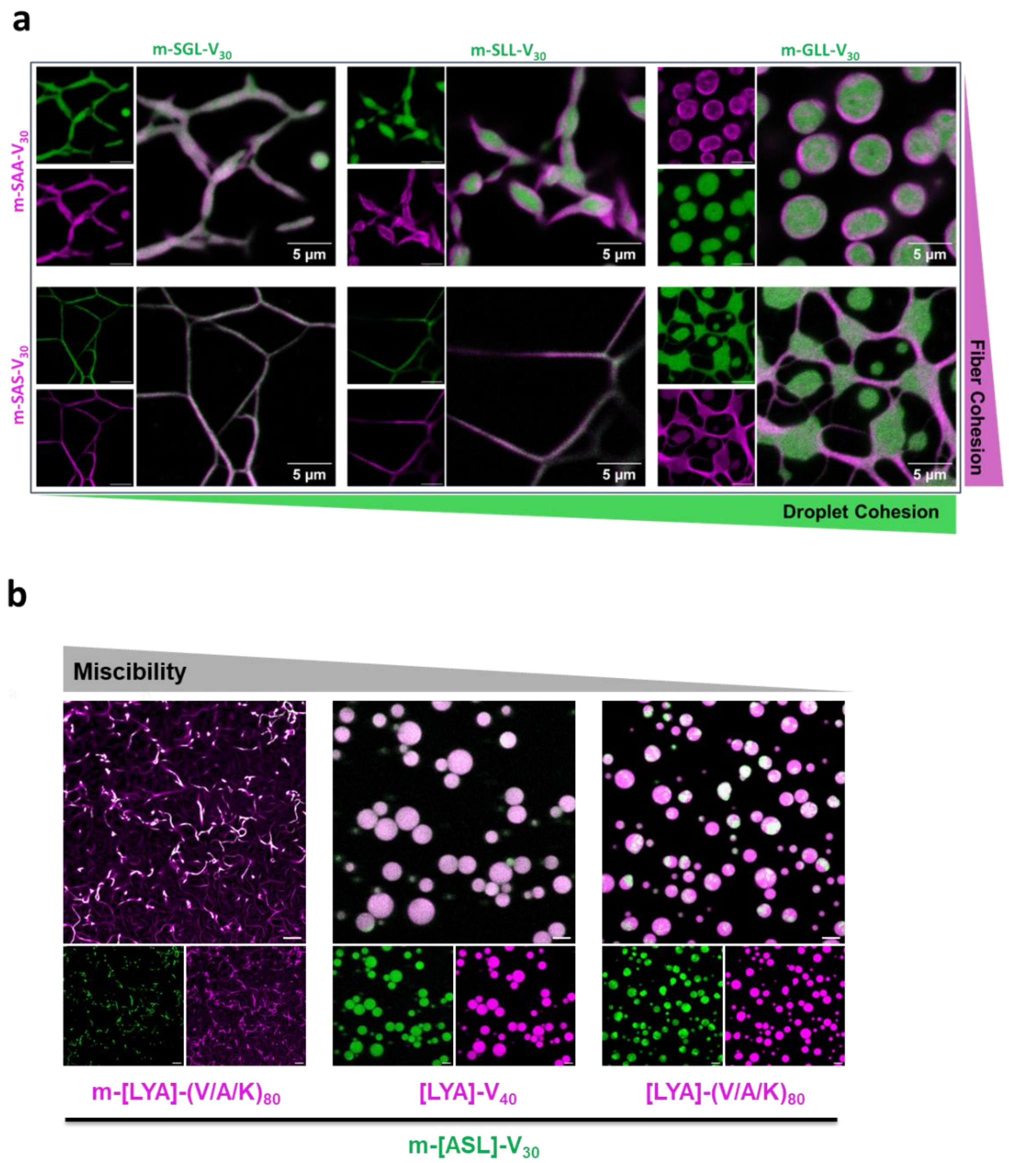

**Supplementary Figure 49. Adhesive interactions regulate the material state and architecture of multiphase condensates. a**, Influence of relative cohesive strength on thermodynamic outcomes in the high-adhesion limit (identical scaffolds). Strong fiber-formers (e.g., m-[SAS]-$V_{30}$) template the conversion of weak droplets into fibrillar networks, whereas stronger droplet-former (e.g., m-[GLL]-$V_{30}$) wet and sequester weak fibers (m-[SAA]-$V_{30}$). Balanced cohesion yields arrested co-assembled networks. Scale bars, 5 µm. **b**, Confocal images of a fiber-forming scaffold (m-[ASL]-$V_{30}$, green) mixed with various droplet-forming scaffolds (red) that differ in IDP sequence and lipidation state. (left) Mixing with a highly miscible, myristoylated partner (m-[LYA]-$(V/A/K)_{80}$) induces droplet-to-fiber conversion, consistent with templated assembly. (middle) Mixing with the unlipidated [LYA]-$V_{40}$ yields compositionally uniform liquid droplets, indicating that strong adhesive interactions override the intrinsic cohesive tendency of the fibers. (right) Mixing with the least miscible partner, [LYA]-$(V/A/K)_{80}$, produces multiphase condensates with segregated liquid (red) and solid (green) sub-compartments. Scale bars, 10 µm.

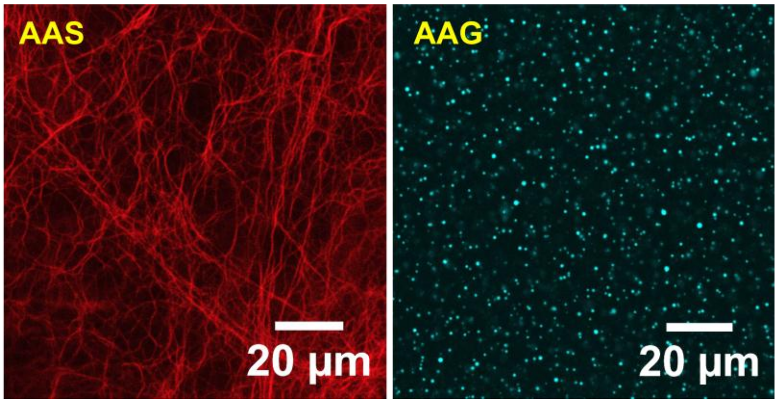

**Supplementary Figure 50. Preservation of condensate material properties in hybrid hydrogels.** Confocal micrographs of AZDye 647–labelled variants in hybrid Matrigel composites show that m-[AAS]-$V_{30}$ forms fibrillar networks (red), whereas m-[AAG]-$V_{30}$ forms spherical droplets (pseudo-colored blue).

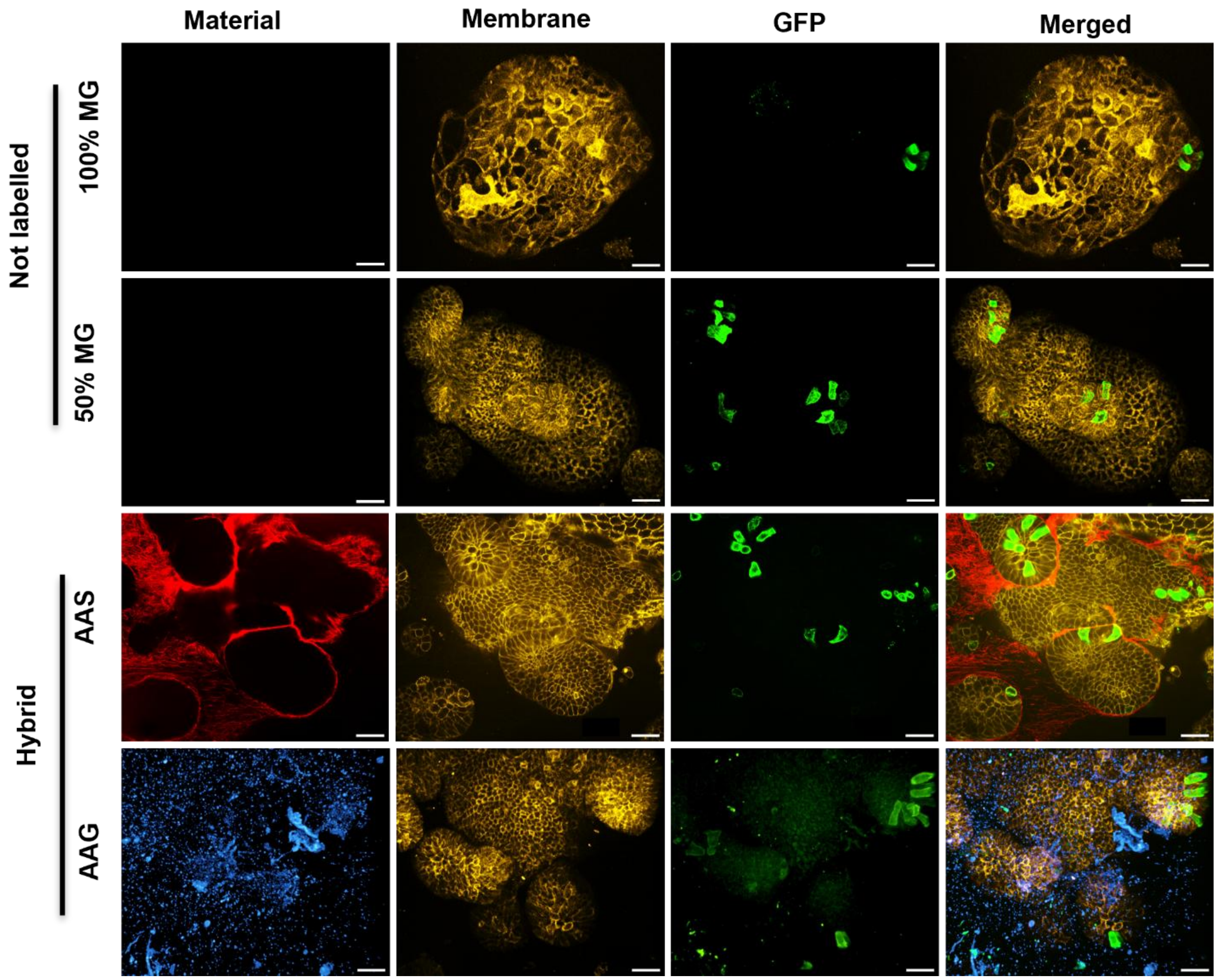

**Supplementary Figure 51. Material distribution and epithelial organization in Matrigel and hybrid ELP–Matrigel matrices.** Confocal images of intestinal organoids (day 4) cultured in Matrigel controls versus hybrid matrices containing fiber-forming m-[AAS]-$V_{30}$ or droplet-forming m-[AAG]-$V_{30}$. Columns display a constitutive membrane reporter (mT; yellow), Defa4-GFP$^{+}$ Paneth cells (mG; green), and labeled material. Controls show smooth epithelia with sparse Paneth cells. Hybrid m-[AAS]-$V_{30}$ matrices form fibrillar networks (red) surrounding Defa4-GFP$^{+}$-enriched crypts, whereas m-[AAG]-$V_{30}$ matrices contain dispersed condensates (blue) with minimal epithelial association. Scale bars, 30 µm.
